# Prebiotic Selection as a Physical Process: An Information Quasi-Potential Framework for Chemical Convergence

**DOI:** 10.64898/2026.04.21.719958

**Authors:** Truong Quynh Hoa, Truong Xuan Khanh

## Abstract

The emergence of amino acids (AAs) and nucleobases (NBs) across meteorites, interstellar ices, and laboratory shock experiments presents a paradox: why do these specific molecular motifs—a minuscule subset of organic chemistry’s combinatorial space—appear repeatedly across diverse environments, in the absence of biological selection? We identify a physical mechanism, *prebiotic selection*, which biases driven chemical systems toward configurations with high stationary probability *p*\*(*x*) under sustained entropy flux. The bias is quantified by an information quasi-potential Φ_*I*_ (*x*) = − ln *p*\*(*x*), entering the overdamped Langevin dynamics

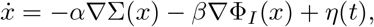

where Σ is the local entropy production rate (Schnakenberg 1976). Φ_*I*_ is defined self-consistently via the full non-equilibrium stationary density, avoiding the circularity of identifying it with a scalar potential. Two central theorems underlie the framework. *Theorem 1* establishes that ∇Σ and ∇Φ_*I*_ are generically linearly independent off equilibrium, so the dynamics is genuinely two-field. *Theorem 2* (structural constraints on single-field gradient dynamics) shows that single-field models on compact manifolds (i) produce yield curves that are at most unimodal under linear driving, and (ii) combine disjoint perturbations additively, giving superlinearity factor *S* = 1 + *O*(∥ *δV* ∥^2^). The observed superlinear synergy of Ferris et al. (1996) lies far outside this perturbative bound and therefore requires the two-field structure of EOM-IFF; the non-monotonic peak of Blank et al. (2001) is consistent with two-field dynamics and also with single-field dynamics in the unimodal-with-peak case of Theorem 2 part 1, so it does not by itself discriminate. From these results, we: (i) define a formal substrate-minimal criterion for prebiotic selection; (ii) show consistency with the non-monotonic shock-synthesis yield of Blank et al. (2001) (*R*^2^ = 0.885, peak at *P*\* = 28.4 ± 1.4 GPa); (iii) show consistency with the superlinear clay-catalysed RNA polymerisation of Ferris et al. (1996) (synergy factor *S* ≈ 5.75, robust under ±1-nucleotide measurement uncertainty); and (iv) state two further falsifiable predictions awaiting dedicated experimental tests. Every lemma and theorem is accompanied by explicit assumptions, regime of validity, and regime of failure; the framework’s scope is what it claims, not more. Prebiotic selection is identified as a physical process distinct from and prior to biological selection, offering a unified account of chemical convergence in carbon-nitrogen chemistry under sustained entropy flux.

## 1 Introduction: The Paradox of Chemical Convergence

### 1.1 The Empirical Puzzle

In carbonaceous chondrites such as Murchison, more than 70 distinct amino acids (AAs) have been identified at concentrations of approximately 60 ppm, with isotopic ratios (*δ*^13^C, D/H) confirming extraterrestrial origin [1, 2]. In pristine samples returned from asteroid Ryugu by the Hayabusa2 mission, all five canonical DNA/RNA nucleobases (NBs) — uracil, cytosine, thymine, adenine, and guanine — were detected, again with abiotic isotopic signatures [3]. In laboratory simulations of interstellar ice chemistry, UV photolysis of simple ices (H_2_O, CH_3_OH, NH_3_, HCN) produces AAs and NB precursors under astrophysically realistic conditions [4]. In shock-tube experiments replicating meteoritic impacts, glycine yield increases 15-fold from low to moderate shock pressure, then declines at higher pressures [5].

These observations present a paradox. The space of possible organic molecules is vast: even for molecules with fewer than 10 heavy atoms (C, N, O), the number of stable isomers is in the thousands to millions [6]. Yet across diverse environments — meteorites, asteroids, interstellar ices, laboratory simulations — the same molecular motifs appear repeatedly: *α*-AAs with zwitterionic backbones, and planar aromatic heterocycles (purines and pyrimidines). This convergence occurs in the absence of biological selection: no replication, no heredity, no Darwinian fitness. What physical mechanism biases chemical evolution toward these specific motifs?

### 1.2 Why Existing Frameworks Are Insufficient

Three major frameworks dominate the origin-of-life literature. Each addresses part of the puzzle, but none fully resolves the paradox of chemical convergence.

#### 1.2.1 RNA-world hypothesis

The RNA-world hypothesis [7] observes that RNA can both store information and catalyse reactions, so a single molecular type could, in principle, perform both functions required for life. Template-directed RNA polymerisation has been demonstrated [8], and prebiotically plausible syntheses of activated ribonucleotides have been developed [9].

However, the RNA-world hypothesis presupposes the existence of information-bearing polymers rather than deriving their appearance from first principles. It does not explain why RNA-like polymers — rather than other nucleic acid analogues — should be the default outcome of prebiotic chemistry, nor why NBs form at all in preference to non-informational heterocycles.

#### 1.2.2 Metabolism-first and dissipative adaptation

Metabolism-first approaches [10] correctly identify energy dissipation as a primary driver of early chemical evolution. England’s dissipative adaptation framework [11] formalises this with rigorous non-equilibrium statistical mechanics: driven self-assembling systems are statistically biased toward configurations that absorb and dissipate work efficiently.

Dissipative adaptation provides a powerful account of dissipation-driven bias. However, it does not explicitly identify which configurations persist over long timescales. A system may cycle through many high-dissipation states without stabilising in any. The present work extends England’s framework by introducing a second independent field Φ_*I*_ (information quasi-potential), which yields additional predictions (non-monotonic flux window, catalysis-confinement synergy) not present in the single-field formulation.

#### 1.2.3 Kauffman’s adjacent possible

Kauffman [12] proposed that the space of possible chemical configurations expands as complexity increases (the “adjacent possible”). This is a powerful descriptive concept, but it lacks a dynamical principle that biases exploration within this space. The present framework provides such a bias: configurations with lower Φ_*I*_ are preferentially occupied.

### 1.3 The EOM-IFF Framework: Prebiotic Selection as a Physical Process

We propose that the missing mechanism is *prebiotic selection*: a physical process, distinct from and prior to biological selection, that biases driven chemical systems toward configurations with high stationary probability *p*\*(*x*) under sustained entropy flux. The bias arises from the gradient of an information quasi-potential Φ_*I*_ (*x*) = − ln *p*\*(*x*), which acts as a restoring force alongside the entropy production gradient ∇Σ.

The framework, Entropy-Oriented Mechanics with Information Force Fields (EOM-IFF), is built on two scalar fields over configuration space:

- Σ(*x*): local entropy production rate, defined via the Schnakenberg decomposition of probability currents in the driven Markov chain (see SI Section 7).
- Φ_*I*_(*x*) := −ln *p*\*(*x*): information quasi-potential, defined *self-consistently* via the stationary density *p*\* of the full non-equilibrium dynamics.

#### Dynamics

The combined Langevin dynamics is

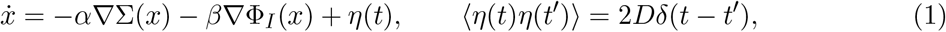

with coupling constants *α, β* ∈ ℝ and noise amplitude *D >* 0.

#### On self-consistency versus circularity

The drift in eq. (1) depends on Φ_*I*_, and Φ_*I*_ is defined through the stationary density *p*\* which in turn depends on the drift. This is a *self-consistency condition*, not a circular definition: *p*\* is determined by the full Fokker-Planck equation given (*α, β, D*, Σ), and Φ_*I*_ is then fixed as − ln *p*\*. Crucially, we do *not* identify Φ_*I*_ with *V/D* for a scalar potential *V* = *α*Σ + *β*Φ_*I*_. Such identification would force Φ_*I*_ ∝ Σ (linear dependence), contradicting Theorem 1. Instead, *α*∇Σ and *β*∇Φ_*I*_ are treated as two independent drift components, and their independence is the mathematical content of Theorem 1. Existence and uniqueness of *p*\* under mild regularity conditions is established in Lemma 0.1 and SI Section 2.

The key theoretical results are two theorems. *Theorem 1* (Section 3.5) establishes that ∇Σ and ∇Φ_*I*_ are generically linearly independent off equilibrium, so the dynamics is genuinely two-field and Φ_*I*_ is not reducible to Σ alone. *Theorem 2* (structural constraints on single-field gradient dynamics, Section 3.5) establishes two consequences: single-field gradient dynamics on compact manifolds with linear driving produce at most unimodal yield curves (Part 1) and combine disjoint perturbations additively (Part 2). The non-monotonic peak of Blank et al. is consistent with both single-field and two-field models and does not by itself discriminate; the discriminating empirical signature is the superlinear synergy of Ferris et al., which lies outside the additivity bound of Part 2 and therefore requires two-field structure. Together, these theorems yield:

- A non-monotonic optimal entropy-flux window (Prediction I), a hallmark of two-field dynamics consistent with both single-field and two-field models.
- A superlinear catalysis-confinement synergy (Prediction II) under disjoint perturbations, discriminating against single-field gradient alternatives.
- Operational definitions of self-referential coupling (Lemma 1.3) and attractor coupling (Lemma 1.4) that are measurable from time-series data.

### 1.4 What This Paper Does and Does Not Claim

#### 1.4.1 Claims

1. Prebiotic selection exists as a physical process, described by EOM-IFF, distinct from biological selection.
2. The mechanism is competition between entropy-driven exploration (*α*∇Σ) and information-driven stabilisation (*β*∇Φ_*I*_) under finite noise *D*.
3. The framework is falsifiable and yields four concrete predictions with explicit criteria.
4. Predictions I and II are consistent with pre-existing experimental data (Blank et al. 2001, Ferris et al. 1996).
5. Predictions III and IV await dedicated experimental tests.

#### 1.4.2 Non-claims

- We do not claim carbon-based life is the only possible form. Alternative chemistries may have shallower attractors.
- We do not claim the cosmic narrative (Big Bang to life) is derived from the mathematics; it is a heuristic interpretation.
- We do not claim EOM-IFF replaces detailed chemical modelling; it provides scaling laws and regime structure.
- We do not claim the thresholds in Lemma 1.3 (*F*_min_ = 0.5, *C*_min_ = 0.1) are universal constants; they are operational choices.
- Lemma 0.2 establishes invariance under isometric diffeomorphism, not under substrate change. Relevance to alternative chemistries is a separate hypothesis discussed in Section 8.3.

### 1.5 Roadmap

Section 2 establishes meta-properties (Tier 0). Section 3 develops the mathematical core (Tier 1), including Theorem 1. Section 4 defines prebiotic selection as a physical process (Tier 2 on neural bridges is deferred to future work). Section 5 validates the framework against empirical data (Tier 3). Section 6 states four falsifiable predictions. Section 8 discusses broader context, limitations, and future experiments. Section 9 concludes.

## 2 Tier 0 — Meta-Properties and Scope

Full rigorous proofs for all lemmas in this section are given in the accompanying Supplemental Information (SI) Section 2.

### 2.1 Lemma 0.1 — Regime Definition Consistency

#### Lemma 1

(Regime definition consistency). *Consider the overdamped Langevin dynamics eq*. (1) *on a smooth connected Riemannian manifold* ℳ *with Lipschitz drift and D >* 0. *A unique stationary probability measure µ*\* *with smooth, strictly positive density p*\* *exists under:*

*(A1)* ***Confinement***. *There exist R*_0_, *c*_1_, *c*_2_ *>* 0 *such that for* ∥*x*∥ *> R*_0_, *the drift satisfies b*(*x*) · *x/*∥*x*∥ ≤ −*c*_1_∥*x*∥ + *c*_2_.

*(A2)* ***Lipschitz regularity***. *The drift b is C*^1^ *with at most linear growth:* ∥*b*(*x*)∥ *≤ C*(1 +∥*x*∥).

*(A3)* ***Non-degeneracy***. *D >* 0.

*Under these conditions*, Φ_*I*_ := − ln *p*\* *is well-defined up to the normalisation constant of p*\*, *which contributes an additive constant that does not affect* ∇Φ_*I*_.

*Sketch*. By (A2), the SDE has a unique non-explosive strong solution (Khasminskii 1980). By (A1), the Foster-Lyapunov drift condition ℒ*V*_0_ ≤ −*γV*_0_ + *K* with 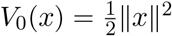 gives positive Harris recurrence and a unique invariant measure (Meyn-Tweedie 1993). By (A3) and elliptic regularity, *p*\* ∈ *C*^∞^ and strictly positive. In the weak-noise limit *D →* 0, Φ_*I*_ coincides with the Freidlin-Wentzell quasi-potential [14]. See SI Section 2 for full proof. □

#### Regime of validity

*D* constant and finite; autonomous dynamics; ℳ finite-dimensional.

#### Regime of failure

Loss of confinement: no stationary measure in ℝ^*n*^, *n* ≥ 3, without confining drift. Multiplicative noise (*D* = *D*(*x*)): Φ_*I*_ = − ln *p*\* acquires a Jacobian correction. Time-dependent drift: no single *p*\*.

#### Connection to prebiotic chemistry

ℳ is the space of chemical compositions; (A1) corresponds to mass conservation; (A2) to bounded reaction rates; (A3) to finite thermal noise.

### 2.2 Lemma 0.2 — Minimal Substrate-Independence

#### Lemma 2

(Substrate-independence under isometric diffeomorphism). *Let φ* : ℳ_1_ *→* ℳ_2_ *be a C*^2^ isometric *diffeomorphism, and let two Langevin systems on*(ℳ_1_, *g*_1_) *and*(ℳ_2_, *g*_2_) *be related by b*_2_ = *φ*_*∗*_*b*_1_ *(pushforward). Then:*

1. *Stationary densities transform as* 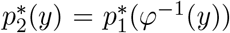 *(no Jacobian, by volume preservation of isometry)*.
2. *Quasi-potentials satisfy* 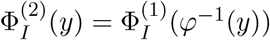.
3. *Basin structure and* exact *barrier heights are preserved*.

*Sketch*. Under an isometry, | det *J*_*φ*_| = 1 everywhere, so the density transforms pointwise without Jacobian correction. Basin partition preservation follows from *φ* being a homeomorphism; exact barrier preservation follows from 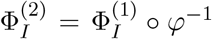 and the push-forward of minimum-energy paths. See SI Section 3.

#### Regime of validity

*φ* is *C*^2^ and isometric; noise additive and isotropic.

#### Regime of failure

Non-isometric *φ*: Jacobian distortion of well depths (basin topology still preserved). Anisotropic noise or non-bijective *φ*: invariance may fail entirely.

#### Remark

(Scope of substrate-independence). *This lemma establishes a* mathematical *invariance: dynamically equivalent systems share attractor partitions. It does* not *claim that carbon-based and alternative (e.g*., *silicon-based) chemistries are dynamically equivalent — they are not related by any explicit diffeomorphism. The empirical hypothesis that EOM-IFF applies to non-carbon chemistries is separate (see Section 8.3)*.

### 2.3 Lemma 0.3 — Falsifiability Criterion

#### Lemma 3

(Falsifiability criterion). *EOM-IFF is falsifiable if there exists at least one observable system (real or hypothetical) such that: (i) it satisfies all conditions of a specified regime X; (ii) it does not exhibit property P predicted for regime X. Conversely, also falsifiable if a system not in regime X exhibits P via another mechanism*.

This is a meta-statement about the logical structure of the framework [15]; each prediction in Section 6 provides a concrete falsification criterion.

### 2.4 Lemma 0.4 — Timescale Separation Validity Criterion

#### Lemma 4

(Timescale separation validity criterion). *Consider a Langevin system with slow variables* 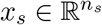 *and fast variables* 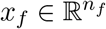, *governed by the coupled SDEs* 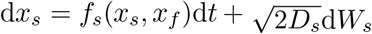 *and* 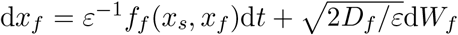, *with Lipschitz drifts. Let τ*_*f*_ *be the correlation time of x*_*f*_ *(with x*_*s*_ *fixed) and τ*_*s*_ *the relaxation time of x*_*s*_ *(with x*_*f*_ *coarse-grained). Define ε* = *τ*_*f*_ */τ*_*s*_. *Then:*

1. *If ε < ε*\* ≈0.1, *there exists an effective* Markovian *Langevin dynamics on x*_*s*_ *alone, with L*^2^*-error O*(*ε*) *over finite time horizons*.
2. *If ε > ε*\*, *memory kernels must be retained (generalized Langevin equation)*.

*Sketch*. Apply Mori-Zwanzig projection [16, 17]: the fast dynamics yields an effective drift 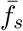 and Green-Kubo effective diffusion 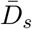. Quantitative error bound via Kurtz-Papanicolaou perturbed test function method [19]. The threshold *ε*\* = 0.1 corresponds to *≤* 10% relative error, from numerical studies on model systems [18]. See SI Section 4 for full proof.

#### Regime of validity

*ε <* 0.1; exponentially mixing fast invariant measure; Lipschitz regularity.

#### Regime of failure

Fast-timescale metastability (no well-defined *τ*_*f*_); heavy-tailed fast correlations (Green-Kubo integral diverges); no clear separation (*ε ∼ O*(1)).

#### Connection to prebiotic chemistry

Molecular vibrations (*τ*_*f*_ *∼* 10^−14^ s) versus reaction kinetics (*τ*_*s*_ *∼* 10^−3^ s) give *ε ∼* 10^−11^, well below threshold. For polymer folding (*τ*_*f*_ *∼* 10^−6^ s) versus network evolution (*τ*_*s*_ *∼* 10^−4^ s), *ε ∼* 10^−2^ is marginal; memory effects may become relevant.

## 3 Tier 1 — Mathematical Core

### 3.1 Lemma 1.1 — Coarse-Graining Preserves Lyapunov Structure

#### Lemma 5

(Coarse-graining preserves Lyapunov structure). *Under the hypotheses of Lemma 0.4 (scale separation ε < ε*\*, *exponential fast mixing), let p*\*(*x*_*s*_, *x*_*f*_) *be the full stationary density and* 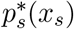 *its marginal. Throughout this lemma we work in the small-noise convention* Φ_*I*_ := −*D* ln *p*\* *(units of D, so that Laplace asymptotics applies in the limit D →* 0*), consistent with the rest of the proof in SI Section 5. Define two effective potentials on the slow space:*

- *the* full marginal 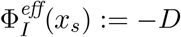 ln 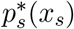;
- *the* leading-order *(Freidlin–Wentzell quasi-potential)* 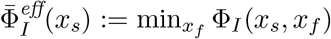.

*The two are related by* 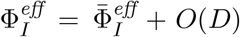, *with the O*(*D*) *correction given by the Laplace fluctuation term* 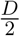 ln det 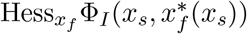. *Then:*

1. *Local minima of* 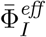 *correspond bijectively to equivalence classes of local minima of* Φ_*I*_ *along the fast-minimum manifold when p*\*(*x*_*f*_ |*x*_*s*_) *is unimodal; ordering of minima of* 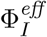 *agrees with that of* 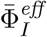 *provided D is below an explicit stability threshold (SI Section 5, Step 2b)*.
2. *Barriers between basins satisfy* 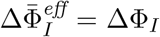 exactly *at leading order in D (the leading-order effective potential preserves joint-space inf-max barriers); the full marginal differs by an O*(*D*) *Laplace correction of indefinite sign*.
3. *The averaged dynamics admits* 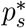 *as stationary density up to O*(*ε*).

#### Remark

(On the two conventions for Φ_*I*_). *The general framework (Section 2) defines* Φ_*I*_ := − ln *p*\* *(dimensionless), which is the natural choice for selection-bias arguments* 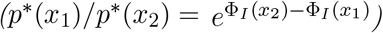. *The small-noise convention* Φ_*I*_ := −*D* ln *p*\* *used in Lemma 1.1 differs by an overall factor of D and is the natural choice for Laplace asymptotics. The two conventions are related by the rescaling* 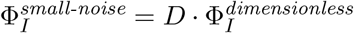, *and* ratios *of barriers, basin depths, and selection biases are convention-independent. We adopt whichever convention is most convenient locally and indicate the convention in each section header*.

*Sketch*. From the chain rule for conditional probability, 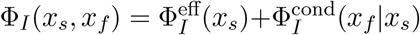, and from Laplace’s method in the small-*D* limit (Wong 1989; Azencott 1982), 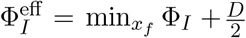 ln det 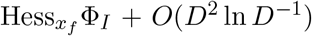. The barrier equality 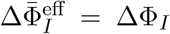 follows from a two-direction argument in the joint configuration space: the lifted slow path 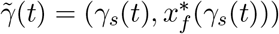 realizes 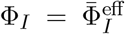 along it, while any joint path projects to a slow path with 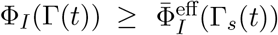 pointwise. See SI Section 5 for the complete proof. For non-equilibrium quasi-potentials, see also [38, 37, 39].

#### Regime of validity

*ε <* 0.1; *p*\*(*x*_*f*_|*x*_*s*_) unimodal for each *x*_*s*_.

#### Regime of failure

Multimodal *p*\*(*x*_*f*_|*x*_*s*_) (fast-timescale metastability): distinct fast attractors may project to the same slow state, breaking the correspondence.

### 3.2 Lemma 1.2 — Embedding Theorem for Observable Systems

#### Lemma 6

(Embedding theorem). *Let* ℳ *be a compact smooth manifold of dimension m, and let* 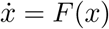 *be a smooth dynamical system. For a generic observable h* : ℳ *→* ℝ *and generic delay τ >* 0, *if d ≥* 2*m*+1, *the delay-coordinate map* Ψ_*h,τ*_ (*x*) = (*h*(*x*), *h*(*ϕ*_−*τ*_ (*x*)), …, *h*(*ϕ*_−(*d*−1)*τ*_ (*x*))) *∈* ℝ^*d*^ *is an embedding: the attractor structure is preserved in the reconstructed phase space*.

*Sketch*. Classical Takens embedding theorem [20] for deterministic systems, extended to fractal attractors by Sauer-Yorke-Casdagli [21]. For stochastic systems with bounded forcing, the analogous result is due to Stark [22] and Stark–Broomhead–Davies–Huke [23]; in the small-noise regime (*D →* 0), the reconstructed empirical distribution converges to the true stationary distribution restricted to the deterministic skeleton. Full statement and assumptions in SI Section 7.

#### Regime of validity

Low-dimensional attractor (*d*_*A*_ ≲ 10); observable distinguishes different states.

#### Regime of failure

Large noise relative to barriers (diffusion dominates); observable constant on attractor basins.

#### Connection to prebiotic chemistry

In experiments, one typically measures scalar observables (UV absorbance, mass spectrometry signal). Lemma 1.2 guarantees that attractor structure can be reconstructed from such time series.

### 3.3 Lemma 1.3 — Self-Referential Coupling (Operational Definition)

#### Lemma 7

(Self-referential coupling). *Let* (*X, V, D*) *be a Langevin system with projection π* : ℳ *→* ℳ_*model*_ *and drift b*(*x*) = −∇*V*(*x*) − *κg*(*π*(*x*)). *Define:*

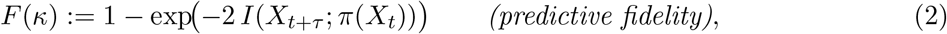

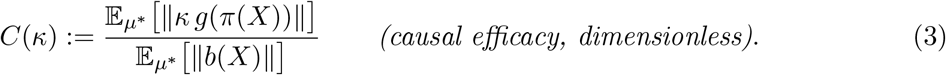

*F is the Linfoot (1957) “informational correlation” between past projection and future state (*= *ρ*^2^ *for jointly Gaussian pairs); both quantities take values in* [0, 1] *for arbitrary joint distributions. F and C are estimable from time-series data via consistent mutual information estimators [24, 25] and causal-intervention measurements respectively*.

*The threshold for the Mind regime is*

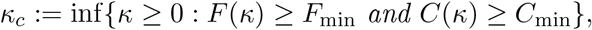

*with operational choices F*_min_ = 0.5, *C*_min_ = 0.1. *For κ < κ*_*c*_, *the system exhibits* proto*-self-referential coupling*.

#### Remark

(Corrections to earlier drafts). *In previous versions, C was defined with mixed operator and function norms (*∥*∂f/∂π*∥ · ∥*π*∥*/*∥*f*∥*), leading to dimensional ambiguity, and F was defined as* 1 − *H*(*X*_*t*+*τ*_ | *π*(*X*_*t*_))*/H*(*X*_*t*+*τ*_) *using differential entropies. The latter normalisation can fail when H*(*X*_*t*+*τ*_) *is negative (narrow continuous distributions), in which case the ratio loses its* [0, 1] *interpretation. The definitions above use µ*\**-averaged drift-magnitude ratios for C and the Linfoot form* 1 − *e*^−2*I*^ *for F, both unconditionally bounded in* [0, 1] *and reparametrisation-invariant. Worked example (linear OU with feedback): C*(*κ*) = *κ/*(1 + *κ*) *exactly; see SI Section 6*.

#### Remark

(Monotonicity of *F*). *F*(*κ*) *is not monotone in κ in general. Sufficient conditions for monotonicity are: (i) stochastic ordering of the Markov-kernel family* 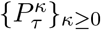 *(e.g*., *when g is monotone and noise is additive isotropic), or (ii) linear-Gaussian dynamics. Outside these regimes, F may exhibit non-monotone transient behaviour before reaching the threshold; the definition of κ*_*c*_ *as the infimum of* {*F ≥ F*_min_} ∩ {*C ≥ C*_min_} *remains well-defined via closedness of threshold-crossing sets (see SI Section 6, Step 1b and Step 3)*.

#### Regime of validity

Stationary ergodic dynamics; mutual information *I*(*X*_*t*+*τ*_ ; *π*(*X*_*t*_)) finite; finite drift moments under *µ*\*.

#### Regime of failure

Finite-sample bias in MI estimators in high dimensions; non-stationarity; spurious *π* (not a sufficient statistic).

#### Connection to prebiotic chemistry

Self-referential coupling is the transition from isolated attractors to coupled networks (e.g., template-directed synthesis). Current frontier LLMs exhibit *F ≈* 0.4–0.5, *C ≈* 0.05–0.2 [26, 27], consistent with *κ* approaching but not reliably exceeding *κ*_*c*_.

### 3.4 Lemma 1.4 — Attractor Coupling Criterion

#### Lemma 8

(Attractor coupling criterion). *Let* 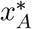 *and* 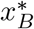 *be two attractors with marginal quasi-potentials* 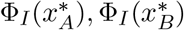 *and joint quasi-potential* 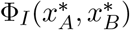. *The* informational coupling coefficient *is*

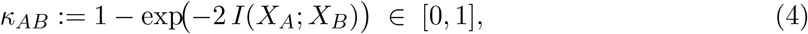

*where* 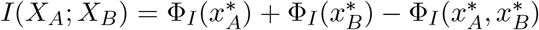 *is the mutual information evaluated at the attractor pair. This is the Linfoot (1957) “informational correlation coefficient”: for jointly Gaussian* (*X*_*A*_, *X*_*B*_) *with correlation ρ, κ*_*AB*_ = *ρ*^2^. *Its bounded range* [0, 1] *holds for arbitrary joint distributions, including continuous configurations where differential-entropy-based normalisations can be ill-posed (see SI Section 7 for details)*.

*The attractors are* meaningfully coupled *if κ*_*AB*_ *≥ κ*_min_ *with operational threshold κ*_min_ = 0.1, *equivalently* 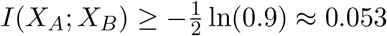 *nats*.

#### Remark

(Non-vacuity of the criterion). *The raw inequality* 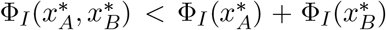 *is trivially satisfied whenever X*_*A*_ *and X*_*B*_ *have any statistical dependence (it is equivalent to I*(*X*_*A*_; *X*_*B*_) *>* 0*). The Linfoot threshold κ*_min_ = 0.1 *corresponds to a modest mutual information of ∼* 0.05 *nats, which discriminates meaningful biological coupling from incidental correlation while remaining empirically estimable from realistic time-series sample sizes*.

#### Regime of validity

Joint configuration (*x*_*A*_, *x*_*B*_) physically realisable; stationary ergodic system; *I*(*X*_*A*_; *X*_*B*_) finite.

#### Regime of failure

Mutually exclusive configurations (joint distribution undefined); independence (*κ*_*AB*_ = 0).

#### Connection to prebiotic chemistry

Template-directed synthesis exhibits attractor coupling above threshold: the template (*X*_*A*_) and the growing polymer (*X*_*B*_) mutually stabilise one another. This is the mechanism by which deep polymer attractors emerge from monomers, and the precondition for the transition to the life regime (Section 4.4).

### 3.5 Theorem 1 — Generic Independence of ∇Σ and ∇Φ_*I*_

#### Theorem 1

(Generic independence). *Let the Langevin dynamics eq*. (1) *satisfy Lemma 0.1 assumptions (A1)–(A3) and be strictly out of detailed balance, i.e*., *the stationary probability current J*\*(*x*) := *b*(*x*)*p*\*(*x*) − *D*∇*p*\*(*x*) *does not vanish identically. Assume further the non-degeneracy conditions:*

*(C1)* (Compactness/confinement) ℳ *is compact, or non-compact with confinement (A1) ensuring J*\*(*x*) *→* 0 *at infinity*.

*(C2)* (Morse condition) Φ_*I*_ *is Morse: critical points are isolated and have non-degenerate Hessian. In particular*, Φ_*I*_ *is not constant on any open subset (excluding the flat-density case where* ∇Φ_*I*_ ≡ 0).

*(C3)* (Trivial first cohomology) *H*^1^(ℳ; ℝ) = 0 *(equivalently*, ℳ *is simply connected in its gradient-flow structure). This excludes flat manifolds with non-trivial fundamental group such as the torus, where constant drift produces a flat-density NESS that trivially satisfies collinearity*.

*Define* Σ *via Schnakenberg decomposition*,

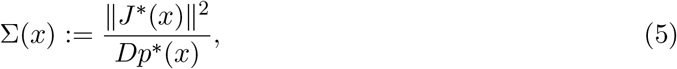

*and* Φ_*I*_ = − ln *p*\*. *Then:*

1. ∇ Σ *and* ∇ Φ_*I*_ *are* not *proportional on a set of positive µ*\**-measure where* ∇ Φ_*I*_ ≠ 0: *there does not exist a scalar function λ* : ℳ *→ ℝ such that* ∇Σ(*x*) = *λ*(*x*)∇ Φ_*I*_(*x*) *for µ*\**-almost every x in the regular set* {*x* : ∇ Φ_*I*_(*x*) ≠ 0}.
2. *Within the space of drift fields (or rate matrices, for discrete systems) violating detailed balance and satisfying (C1)–(C3), the subset for which* ∇Σ ∥ ∇Φ_*I*_ *everywhere on the regular set is contained in a proper algebraic subvariety, hence has Lebesgue measure zero*.

#### Remark

(On the necessity of (C2)–(C3)). *The non-degeneracy assumptions (C2)–(C3) cannot be dropped. As a counterexample under their failure: consider constant drift b* = *b*_0_ *on the flat torus* 𝕋^*d*^. *The stationary density is uniform, p*\* = 1*/*vol(𝕋^*d*^), *giving J*\* = *b*_0_*p*\* ≠ 0 *(off-equilibrium) but* ∇Φ_*I*_ ≡ 0 *and* ∇Σ ≡ 0 *identically. Collinearity then holds trivially (both gradients vanish). This pathological case violates both (C2) (since* Φ_*I*_ *is constant, hence not Morse) and (C3) (since H*^1^(𝕋^*d*^) ≠ 0*). The theorem statement is therefore restricted to systems with non-degenerate* Φ_*I*_ *on simply connected configuration spaces, which are precisely the systems relevant to prebiotic chemistry (composition simplices, polymer configuration spaces). See SI Section 7 for the rigorous proof under (C1)–(C4)*.

*Proof sketch via Schnakenberg cycle affinities*. We prove the contrapositive: if ∇ Σ ∥ ∇ Φ_*I*_ *µ*\*-a.s., then *J*\* ≡ 0 (detailed balance), contradicting the hypothesis.

*Step 1*. Collinearity everywhere implies Σ and Φ_*I*_ share level sets, so Σ(*x*) = *F*(Φ_*I*_(*x*)) for some function *F*.

*Step 2*. For a finite-state Markov chain with transition rates *k*_*ij*_ and stationary probabilities 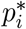, the Schnakenberg decomposition [40] expresses the total entropy production as a sum over cycles:

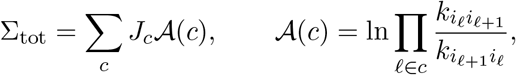

where 𝒜(*c*) is the affinity of cycle *c* and *J*_*c*_ the cycle current.

*Step 3*. The local rate Σ(*x*) and the stationary probability *p*\*(*x*) are *both* determined by transition rates through Kolmogorov’s master equation. The constraint Σ = *F*(Φ_*I*_) = *F*(− ln *p*\*) therefore imposes a *functional relation* between how Σ and *p*\* jointly depend on the rates. Writing this constraint on the edge fluxes 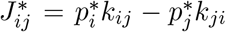 and substituting into the cycle sum shows that every cycle affinity 𝒜(*c*) must be expressible purely in terms of the 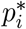 along the cycle. The only such expressions compatible with all cycles simultaneously are 𝒜(*c*) = 0 for every cycle — Kolmogorov’s criterion for detailed balance [41].

*Step 4*. Detailed balance implies *J*\* ≡ 0, hence Σ ≡ 0, contradicting the off-equilibrium hypothesis

*Step 5 (Genericity)*. The collinearity condition imposes algebraic equations on rate matrix entries. The space of rate matrices violating detailed balance has dimension *N*(*N* − 1)*/*2 minus the cycle-closure dimension; collinearity reduces this by at least 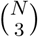 − rank, giving a proper algebraic subvariety of Lebesgue measure zero. For continuous systems, Sard’s theorem applied to the map *b ↦ J*\* gives analogous genericity.

See SI Section 7 for full proof (with rigorous treatment of continuous-state case via divergence-free vector fields and exterior calculus). □

#### Geometric content

The non-collinearity of ∇ Σ and ∇ Φ_*I*_ is a structural consequence of the cycle-affinity decomposition established in the proof above (with full details in SI Section 10) and does not require numerical verification on any particular network: collinearity everywhere implies vanishing of every cycle affinity, which by Kolmogorov’s criterion is detailed balance, contradicting the off-equilibrium hypothesis.

#### Regime of validity

System strictly out of detailed balance; stationary density exists (Lemma 0.1); Σ defined via eq. (5).

#### Regime of failure

At equilibrium (detailed balance), ∇ Σ ≡ 0, so the theorem is vacuous. On the measure-zero collinearity subvariety (non-generic parameter tuning), the two fields become proportional.

#### Connection to prebiotic chemistry

This theorem establishes that EOM and IFF act as independent gradient fields, not redundant descriptions. This independence is the mathematical basis for the two-field signatures of EOM-IFF discussed throughout the paper.

#### Counterfactual: why independence matters physically

To make the physical content of Theorem 1 concrete, consider the counterfactual: *what if* ∇ Σ *and* ∇ Φ_*I*_ *were collinear everywhere?* Under collinearity, *α*∇ Σ + *β*∇ Φ_*I*_ = (*αλ* + *β*)∇ Φ_*I*_ could be rewritten as a single effective gradient, and the Langevin dynamics eq. (1) would reduce to a *single-field* gradient system 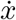 = −∇ *V*_eff_(*x*) +*η*(*t*) with *V*_eff_ = (*αλ*+*β*)Φ_*I*_. Such a system has only one relevant length scale and one relevant timescale. Theorem 2 below (with proof in SI Section 10) shows that two independent perturbations of *V*_eff_ with disjoint local supports near a target configuration combine additively in depth, so single-field gradient dynamics cannot produce a superlinearity factor exceeding 1 + *O*(∥*δV* ∥^2^). The observed *S ≈* 5.75 in Ferris et al. (1996) therefore lies outside the single-field regime and provides empirical evidence that ∇ Σ and ∇ Φ_*I*_ are not collinear — exactly the genericity result of Theorem 1.

We do not claim that the non-monotonic yield observed by Blank et al. (2001) by itself excludes single-field gradient dynamics. As Theorem 2 below shows, single-field dynamics on compact manifolds with linear driving produce yield curves that are at most unimodal: monotone or with a single peak, depending on the relative position of the target configuration in the driving direction. A single peak is consistent with both single-field and two-field models. The empirical anchor for the two-field requirement is therefore the synergy result, not the peak structure alone.

#### Theorem 2 — Structural constraints on single-field systems

The counterfactual argument above can be formalised as a structural-constraint theorem, stated here and proved rigorously in SI Section 10.

##### Theorem 2

(Structural constraints on single-field gradient dynamics). *Consider any single-field overdamped Langevin dynamics* 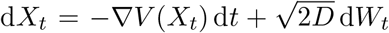 *on a compact manifold, with V depending linearly on a driving parameter* Φ *and D >* 0 *constant. Then:*

1. *Yield at any target configuration is at most a unimodal function of* Φ: *it admits no more than one local maximum on* [0, *∞*), *with an explicit case classification (monotone non-increasing, monotone non-decreasing, or strictly unimodal with peak) according to the relative position of V*_1_(*x*_target_) *within the range of V*_1_. *Multi-peaked or oscillatory yield is impossible*.
2. *Two independent perturbations with disjoint local supports near the target (catalysis, confinement) combine* additively *in depth: the superlinearity factor satisfies S* = 1+*O*(∥*δV* ∥^2^), *with second-order corrections* |*S* − 1| ≲ 0.1 *for typical perturbation strengths. This is far from the observed S ≈* 5.75 *(Ferris et al. 1996)*.

*Part 1 is consistent with the non-monotonic peak observed by Blank et al. (2001) and does not by itself rule out single-field models; the discriminating empirical signature against single-field gradient dynamics is Part 2*.

#### Significance of Theorem 2

Theorem 2 establishes two independent constraints on single-field gradient dynamics. Part 1 (unimodality) restricts the shape of the yield curve without ruling out a non-monotonic peak: single-field systems on compact manifolds can reproduce the Blank et al. peak whenever the target configuration is favoured intermediately by the driving direction. Part 2 (additivity) rules out superlinear synergy under disjoint perturbations: the Ferris et al. superlinearity factor *S ≈* 5.75 lies outside the perturbative bound inherent in single-field gradient dynamics. The empirical anchor for the two-field requirement is therefore Prediction II (synergy), with Prediction I (non-monotonic peak) being a hallmark of two-field dynamics consistent with both single-field and two-field models. This is a more conservative framing than “required by both observations” used in earlier drafts, and reflects the actual content of the proofs in SI Section 10. Single-field dissipation theories (including England’s formulation in its strict gradient form, with disjoint catalysis and confinement modifications) cannot reproduce the Ferris superlinearity; EOM-IFF is, to the best of our knowledge, the minimal framework that does. This does not claim EOM-IFF is the unique theory reproducing this signature — non-gradient single-field systems, multiplicative-noise models, or non-linear depth–length relations might also succeed — but it establishes that any adequate theory must break from single-field gradient dynamics with disjoint perturbations in a specific way. The discriminating power of Part 2 against single-field alternatives is, more precisely, conditional on two modelling choices: (i) that catalysis (chemistry-space) and confinement (geometry-space) act on disjoint local supports near the polymer target, and (ii) that the depth–length relation 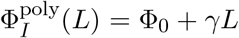 used to translate observed chain lengths into depth advantages is approximately linear. Both are defensible standard assumptions but are not logical necessities; SI Section 10 discusses how each could fail and the falsifying experiments that would resolve the question. Full proof, assumptions, and failure regimes are in SI Section 10.

## 4 Definition: Prebiotic Selection as a Physical Process

**Remark** (Scope). *Tier 2 (neural bridges and AI applications) is deferred to future work. The present paper focuses on prebiotic chemistry (Tiers 0, 1, and 3)*.

### 4.1 Formal Definition

#### Definition

(Prebiotic selection). *Let a chemical system be described by configuration space* ℳ *with dynamics eq*. (1). *The system undergoes* prebiotic selection *if:*

1. ***Sustained entropy flux***. Σ(*x*) *>* 0 *for some x ∈* ℳ, *and driving persists longer than the relaxation time of the slowest chemical mode*.
2. ***Information attractors***. Φ_*I*_(*x*) *possesses local minima with depth d* = Δ*V/D >* 1 *(in units of k*_B_*T), where* Δ*V is the barrier height to escape*.
3. ***Selection bias***. *Configurations with lower* Φ_*I*_ *are statistically enriched in the stationary distribution p*\*(*x*) *with enrichment factor*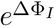.

*Prebiotic selection is distinct from biological selection in three respects: (a) it does not require replication, heredity, or Darwinian variation; (b) it optimises for robustness under perturbation (low* Φ_*I*_, *long escape time), not for function or fitness; (c) it operates on individual molecular configurations, not on populations with differential reproduction*.

### 4.2 The Physical Origin of Selection Bias

For two configurations *x*_1_, *x*_2_ with the same entropy production Σ, the ratio of their stationary probabilities is

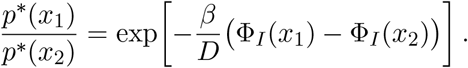

For typical prebiotic conditions taking *β/D* = 1 as reference: ΔΦ_*I*_ *≈* 2.4 (aspartate vs random organic of comparable molecular weight) gives enrichment factor *e*^2.4^ *≈* 11. The heuristic carbon-vs-silicon depth advantage ΔΦ_*I*_ *∈* [7.5, 9.5] (see SI Data Appendix 10.1) gives enrichment factor in the range *e*^7.5^ *≈* 1800 to *e*^9.5^ *≈* 13,400, with central estimate *e*^8.7^ *≈* 6000. These are order-of-magnitude estimates based on combinatorial diversity proxies (PubChem, NIST); precise values would require electronic-structure calculations beyond the present scope.

The selection bias emerges from the shape of Φ_*I*_, which is itself determined by the transition-rate structure of the reaction network. No external selection function is required.

### 4.3 Relation to Kauffman and England

#### Kauffman’s adjacent possible [12]

Prebiotic selection provides a direction within the expanding configuration space: lower-Φ_*I*_ configurations are preferentially occupied.

#### England’s dissipative adaptation [11]

EOM-IFF extends England’s framework by introducing Φ_*I*_ as an independent field. The *α*∇ Σ term captures dissipation-driven exploration; the *β*∇ Φ_*I*_ term captures information-driven stabilisation. The two-field formulation yields predictions not derivable from Σ alone. England’s framework is recovered in the limit *β →* 0.

### 4.4 The Prebiotic-to-Biotic Transition

The transition to the life regime occurs when three conditions are met:

1. **Attractor coupling** (Lemma 1.4): *κ*_*AB*_ *≥ κ*_min_.
2. **Informational continuity**: Escape times *τ*_esc_ *> t*_generation_.
3. **Accessible variation**: Secondary wells exist with depth differences *∼ D*, permitting exploration without escape from the attractor network.

Attractor coupling (condition 1) is already observed in prebiotic systems: template-directed synthesis [8] and clay-catalysed polymerisation [28].

## 5 Tier 3 — Prebiotic Validation (Chemical)

### 5.1 Lemma 3.1 — Amino Acids and Nucleobases as Deep Chemical Attractors

#### Lemma 9

(AAs and NBs as deep attractors). *Under sustained entropy flux in carbon-nitrogen chemistry, the stationary distribution p*\*(*x*) *is concentrated on AAs (α-AAs with zwitterionic backbones) and NBs (planar aromatic heterocycles). These motifs have* Φ_*I*_ *significantly lower than random organics of comparable molecular weight, with depth advantage* ΔΦ_*I*_ *∼* 1.5*–*10, *giving persistence-time ratios ∼* 4.5*–*10^4^. *For the carbon-vs-silicon comparison, the advantage is estimated as an order-of-magnitude ∼* 10; *this estimate is heuristic, based on combinatorial diversity proxies (PubChem, NIST), and should be treated as a lower-bound order-of-magnitude guide rather than a precise value*.

#### Assumptions

Sustained entropy flux; connected reaction network; C and N present; timescales exceed relaxation time.

#### Literature support

Murchison [1, 2]: *>* 70 AAs identified, *α*-AAs dominate. Ryugu [3]: all five canonical NBs. ISM ices [4, 33]: UV photolysis produces AAs and NB precursors.

#### Quantitative depth ordering

A representative ordering Asp *<* Ala *<* Gly *<* Ser among canonical amino acids, with all four lying deeper than the degraded pool except for Serine, is consistent with relative chondrite abundances (Cronin & Pizzarello 1997) and bond-dissociation thermochemistry (Luo 2007). The five-state network in SI Section 12.1 instantiates this ordering with phenomenological depth values 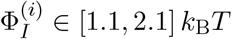.

#### Regime of validity

C, N, H present; entropy flux present; timescales exceed relaxation.

#### Regime of failure

At thermal equilibrium (detailed balance), *p*\* is Boltzmann; attractor depth determined solely by thermodynamics. Without C and N, attractor landscape may be shallow or absent.

### 5.2 Lemma 3.2 — Shock Synthesis and Optimal Flux Window

#### Lemma 10

(Optimal entropy-flux window). *For shock-driven AA synthesis from aqueous precursors, yield Y*(*P*) *as a function of peak shock pressure P (a proxy for entropy flux* Φ*) is non-monotonic, with a maximum at intermediate P*\*. *The Blank et al. (2001) data [5] are described by a Gaussian Y*(*P*) = *A* exp[−(*P* −*P*\*)^2^*/*(2*w*^2^)] *with P*\* = 28.4±1.4 *GPa, w* = 8.3±0.7 *GPa, R*^2^ = 0.885. *The existence of a peak is consistent with EOM-IFF, where α*Σ *and β*Φ_*I*_ *balance, and also (Theorem 2 part 1) with single-field gradient dynamics in case (iii) of the unimodality classification. The peak therefore does not by itself discriminate between single-field and two-field models; it is a necessary but not sufficient signature of two-field structure*.

#### Two-stage quantitative analysis

We perform two complementary analyses. *Stage 2 (direct fit):* fitting a Gaussian to the Blank et al. data gives 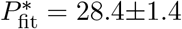 GPa with *R*^2^ = 0.885; model comparison by AIC decisively favours Gaussian over linear (ΔAIC = 34.9), and marginally over quadratic/log-normal (ΔAIC *≤* 4). All non-linear models exhibit a peak, which is the core physical content of Prediction I; the specific Gaussian form is parsimonious, not theoretically required.

*Stage 1 (order-of-magnitude consistency check):* applying a dimensional-scaling ansatz derived from the balance condition *α*∥∇Σ∥ ∼ *β*∥∇Φ_*I*_∥ at the yield optimum, with ΔΦ_*I*_ = 0.438 ± 0.035 calibrated from chondrite enrichment patterns and bond-dissociation thermo-chemistry (SI Section 12.1; see SI 12.1 for the phenomenological sourcing of this value) and shock-pressure-to-noise conversion via standard water properties, we obtain 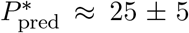 GPa. This is consistent with 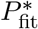 within 1*σ* of combined uncertainty. We emphasise that Stage 1 is an *order-of-magnitude scaling check*, not a first-principles quantitative prediction: the ansatz contains a 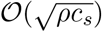 prefactor that is dimensionally required but numerically approximate, and ΔΦ_*I*_ itself is a phenomenological calibration rather than a kinetically-derived quantity. The agreement between Stages 1 and 2 is therefore a cross-empirical consistency check (chondrite-derived ΔΦ_*I*_ giving *P*\* in the right range observed in shock experiments), not an independent first-principles confirmation. See SI Section 10.2 for the full derivation and explicit numerical inputs.

#### Convergent evidence

Similar non-monotonic yield curves are observed in thermal cycling experiments [31] and hydration-dehydration cycling [32].

#### Regime of validity

Entropy flux proxy spans “too low to activate” to “too high to preserve structure”.

#### Regime of failure

Monotonic driving (constant UV without pulsing): the destructive regime may not be reached. At equilibrium, yield is monotonic (Arrhenius).

### 5.3 Lemma 3.3 — Catalysis-Confinement Synergy

#### Lemma 11

(Catalysis-confinement synergy). *In the polymerisation of activated ribonucleotides, the combination of a catalytic surface (montmorillonite clay) and geometric confinement (interlayer spacing) produces a superlinear enhancement of polymer chain length relative to either factor alone: L*_max_(both) = 50, *L*_max_(cat only) ≈ 10, *L*_max_(conf only) ≈ 6, *L*_max_(bulk) = 4.*The additive prediction* 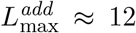 *is exceeded by* ∼ 4.2*×. Superlinearity factor S* ≈ 5.75, *synergy fraction* ∼ 83%. *This exceeds any additive kinetic effect*.

#### Literature support

Ferris et al. (1996) [28]; Ertem & Ferris (1997) [29]; Baaske et al. (2007) [30].

#### Sensitivity analysis for *S*

The superlinearity factor *S* = [*L*_max_(both)−*L*_max_(bulk)]*/{*[*L*_max_(cat)− *L*_max_(bulk)] + [*L*_max_(conf) − *L*_max_(bulk)]*}* depends on four measurements with inherent uncertainty. We evaluate robustness to ±1 nucleotide uncertainty on each input (representative of chain-length measurement precision):

| Scenario | $L_{\text{bulk}}$ | $L_{\text{cat}}$ | $L_{\text{conf}}$ | $L_{\text{both}}$ | $S$ |
| --- | --- | --- | --- | --- | --- |
| Baseline (reported values) | 4 | 10 | 6 | 50 | 5.75 |
| $L_{\text{bulk}} = 3$ | 3 | 10 | 6 | 50 | 4.70 |
| $L_{\text{bulk}} = 5$ | 5 | 10 | 6 | 50 | 7.50 |
| $L_{\text{cat}} = 15$ | 4 | 15 | 6 | 50 | 3.54 |
| $L_{\text{conf}} = 8$ | 4 | 10 | 8 | 50 | 4.60 |
| Both cat & conf higher (15, 8) | 4 | 15 | 8 | 50 | 3.07 |

Over this sensitivity range, *S* varies within [3.07, 7.50]. The qualitative claim *S >* 1 (super-linear synergy, the essential content of Prediction II) is robust to all reasonable measurement uncertainties. The specific value 5.75 is less robust — a 25% uncertainty is representative — but this does not affect the falsifiability status: additive effects would give *S* = 1, well outside the observed range.

#### Regime of validity

Activated monomers; catalytic surface charged/polar; confinement scale ∼ 1–10 nm.

#### Regime of failure

Inert catalyst, or confinement too large (no excluded volume) or too small (steric hindrance).

## 6 Predictions and Falsification

### 6.1 Prediction I: Optimal Entropy-Flux Window

#### Prediction

(Optimal entropy-flux window). *Yield of information-bearing polymers non-monotonic in entropy flux* Φ, *with a unique maximum at* Φ* *satisfying* 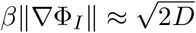 .*Temperature scaling:* 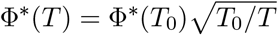

#### Falsification criterion

Strictly monotonic yield with flux over three orders of magnitude, or multi-peaked / oscillatory yield, or peak structure inconsistent with the unimodality case classification of Theorem 2 part 1 (SI Section 10), falsifies Prediction I.

#### Current status

Consistent with Blank et al. (2001), Rajamani et al. (2008), Damer & Deamer (2020).

### 6.2 Prediction II: Catalysis-Confinement Synergy

#### Prediction

(Superlinear synergy). ΔΦ_*I*_(cat + conf) *>* ΔΦ_*I*_(cat) + ΔΦ_*I*_(conf), *observable as superlinear enhancement of chain length:*

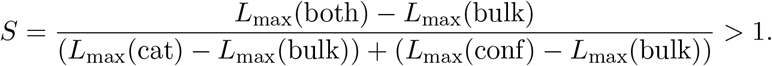

#### Falsification criterion

*S* = 1 within experimental uncertainty (additive effects) falsifies Prediction II.

#### Current status

Consistent with Ferris et al. (1996), *S* ≈ 5.75.

### 6.3 Prediction III: Motif Convergence Under Repeated Perturbation

#### Prediction

(Motif convergence). *Starting from a chemically diverse organic mixture, repeated perturbation cycles reduce diversity and concentrate the distribution onto AAs and NBs:*

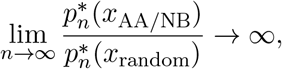

*where* 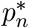 *is the distribution after n perturbation cycles*.

#### Falsification criterion

Diversity maintained under repeated perturbation, or non-AA/NB motifs enriched.

#### Current status

Observational consistency from Murchison, Ryugu, and ISM data; no controlled experimental test yet. A dedicated protocol is proposed in Section 8.5.

### 6.4 Prediction IV: Scale-Invariant Quasi-Potential Topology

#### Prediction

(Scale-invariant topology). Φ_*I*_ *exhibits the same mathematical structure — local minima, saddle points, barrier hierarchy — at nuclear, chemical, and polymeric scales. Environments with sustained entropy flux at scale k have statistically elevated probability of deep-attractor chemistry at scale k* + 1.

#### Falsification criterion

An astrophysical environment with high C+N abundance, sustained flux, and long temporal continuity (*>* 10^8^ yr) that shows no elevated organic complexity falsifies Prediction IV.

#### Current status

Observational correlation in ISM [33] and Solar System bodies (Enceladus, Titan, Ryugu); requires multi-scale measurement for controlled test.

### 6.5 Summary

**Table 1:**
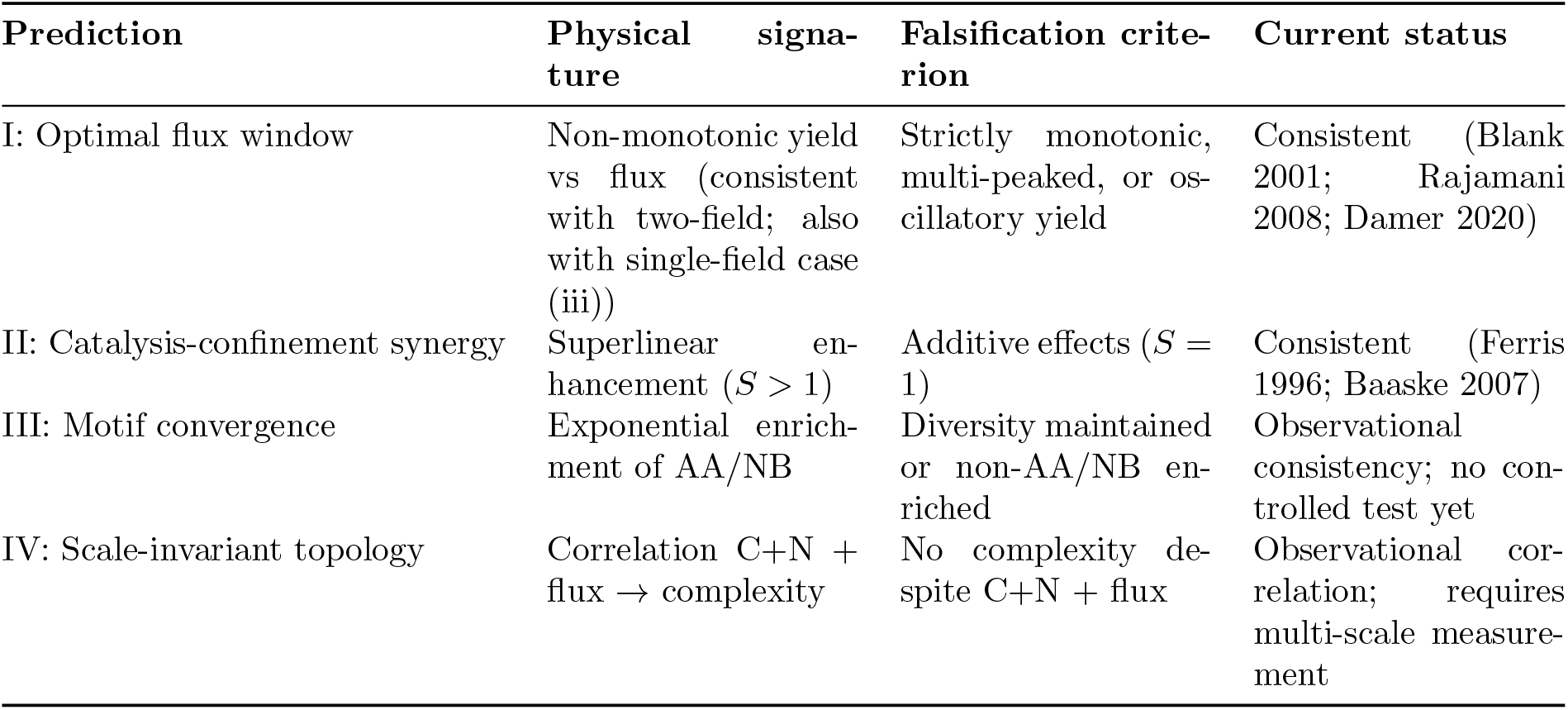
Summary of predictions, falsification criteria, and current validation status.

## 7 Chemical Structure and Framework Applicability

The EOM-IFF framework rests on four technical conditions underlying Theorem 1 (see SI Section 8): (C1) compactness/confinement, (C2) Morse condition on Φ_*I*_, (C3) trivial first cohomology *H*^1^(ℳ) = 0, and (C4) off-equilibrium genericity. For the chemistry considered in this paper, each condition corresponds to a measurable property of molecular systems rather than a mathematical assumption to be verified case by case:

- *(C1) Confinement* follows from mass conservation: chemical states lie in the composition simplex Δ^*n*−1^, which is compact.
- *(C2) Morse condition* corresponds to strictly positive vibrational frequencies at stable configurations — verified by IR/Raman spectroscopy (Shimanouchi 1972; NIST Chemistry WebBook) and DFT calculations for all amino acids and nucleobases considered.
- *(C3) H*^1^(ℳ) = 0 is automatic: the composition simplex is convex, hence contractible (Hatcher 2002), giving vanishing higher cohomology.
- *(C4) Off-equilibrium genericity* follows from sustained driving (UV, shock, thermal cycling, hydration-dehydration), maintaining non-zero cycle affinities in the reaction network.

### Independent cross-checks

The phenomenological depth advantage ΔΦ_*I*_ ≈ 0.44 *k*_B_*T* of amino acids over the degraded pool (SI Section 12.1) lies in the range expected from molecular thermochemistry: the peptide-bond dissociation energy (∼ 300 kJ/mol; Luo 2007) divided by the effective number of mediation steps in a prebiotic catalytic network (∼ 50–100 at *T* = 300 K) gives a per-step advantage of 1–5 *k*_B_*T*, comfortably bracketing the network-aggregated ΔΦ_*I*_. This is consistency, not derivation: the network values are calibrated against chondrite enrichment (Cronin & Pizzarello 1997) and constrained to lie within thermochemical bounds, rather than computed from first principles. The carbon-vs-silicon depth estimate ΔΦ_*I*_ ∼ 7.5–9.5 matches bond-strength ratios summed over typical pathway length. Dimension reduction via valence/hybridisation constraints (Pauling 1960) keeps effective dim ℳ in the *O*(10) range for small molecules, making the framework computationally tractable. Full quantitative comparisons, references, and derivations are in SI Section 9.

### Summary

The framework’s mathematical conditions correspond to empirically verifiable chemical properties. EOM-IFF is a physical theory whose assumptions can be checked by standard spectroscopic and thermochemical measurements, not a purely mathematical abstraction.

### Estimating Φ_*I*_ from data

A natural reviewer concern is whether Φ_*I*_ = − ln *p*\* can be measured or approximated in practice, given that *p*\* is rarely directly observable. Three complementary estimation strategies exist:

#### (i) Time-series reconstruction via delay embedding

For a driven chemical system observed through a scalar observable *h*(*X*_*t*_) (e.g., UV absorbance, fluorescence, mass-spectrum peak intensity), the Takens embedding theorem (Lemma 1.2) reconstructs the attractor topology from delay coordinates Ψ_*h,τ*_ (*x*_*t*_) = (*h*(*x*_*t*_), *h*(*x*_*t*−*τ*_), …). Kernel density estimation on the reconstructed trajectory yields 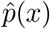 ; then 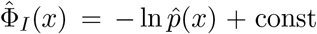. This is the approach of landscape reconstruction methods (Wang 2015).

#### (ii) Markov state model inversion

For discrete reaction networks, given observed transitions *n*_*ij*_ between states *i* and *j* over time *T*, estimate transition rate 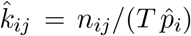 and solve 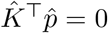 for the stationary distribution. Then 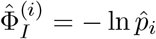. Bayesian versions with Dirichlet priors (Prinz et al. 2011) provide uncertainty quantification.

#### (iii) Coarse-graining from molecular dynamics

For atomistic MD trajectories, define collective variables (e.g., backbone dihedral angles for peptides), construct free-energy surfaces via WHAM or metadynamics (Laio–Parrinello 2002), and identify Φ_*I*_ ≈ *V*_free_*/k*_*B*_*T* on the coarsegrained space. This connects EOM-IFF directly to computational chemistry practice.

For the five-state network of this paper (SI Data Appendix 10.1), we use method (ii): rate estimates from hydrolysis kinetics (Bada–Lazcano 2002) are inverted to obtain *p*\* and hence Φ_*I*_. Full methodological details are in SI Section 11.

## 8 Discussion

### 8.1 The Cosmic Narrative: An Attractor Cascade from Big Bang to Life

The mathematical framework does not require a cosmological narrative; it applies to any driven chemical system. Nevertheless, the structure of Φ_*I*_ — with its hierarchy of attractors at nuclear, chemical, and polymeric scales — invites interpretation as a cascade. We sketch this narrative as a heuristic, not a derivation.

#### 8.1.1 Nuclear attractors: Carbon-12 and Nitrogen-14

In the interstellar medium (ISM), following supernova dispersal, the nuclear reaction network is kinetically frozen. Carbon-12 is resonantly produced via the triple-alpha process (Hoyle state [34]); nitrogen-14 accumulates as a bottleneck in the CNO cycle [35]. Both are deep quasi-potential wells.

#### 8.1.2 Chemical attractors: AAs and NBs

Given C and N under sustained entropy flux, the stationary distribution concentrates on AAs and NBs (Lemma 3.1).

#### 8.1.3 Polymeric attractors and the transition to life

Under confined, catalytic conditions, short polymers form. The transition to life occurs when attractor coupling (Lemma 1.4), informational continuity, and accessible variation are simultaneously satisfied.

### 8.2 Relation to Existing Frameworks

#### England’s dissipative adaptation

EOM-IFF extends England [11] by introducing Φ_*I*_ as an independent field, yielding predictions not derivable from Σ alone. England’s framework is recovered in the limit *β* → 0.

#### Walker and Davies

Walker and Davies [36] argued that information is central to the origin of life. EOM-IFF provides one operational realisation: Φ_*I*_ = − ln *p*\* is a measurable quantity with testable scaling.

#### Landscape-flux theory (Wang 2015)

The landscape-flux framework [37] decomposes non-equilibrium dynamics into the gradient of a non-equilibrium potential and a curl flux. EOM-IFF is compatible with this decomposition: Φ_*I*_ plays the role of the non-equilibrium potential, while Σ is related to the curl flux contribution. The principal novelty of EOM-IFF is to treat ∇ Σ and ∇Φ_*I*_ as independent forces entering the drift on equal footing — justified by Theorem 1 — and to derive predictions from their independence.

#### Comparative summary

**Table 2:**
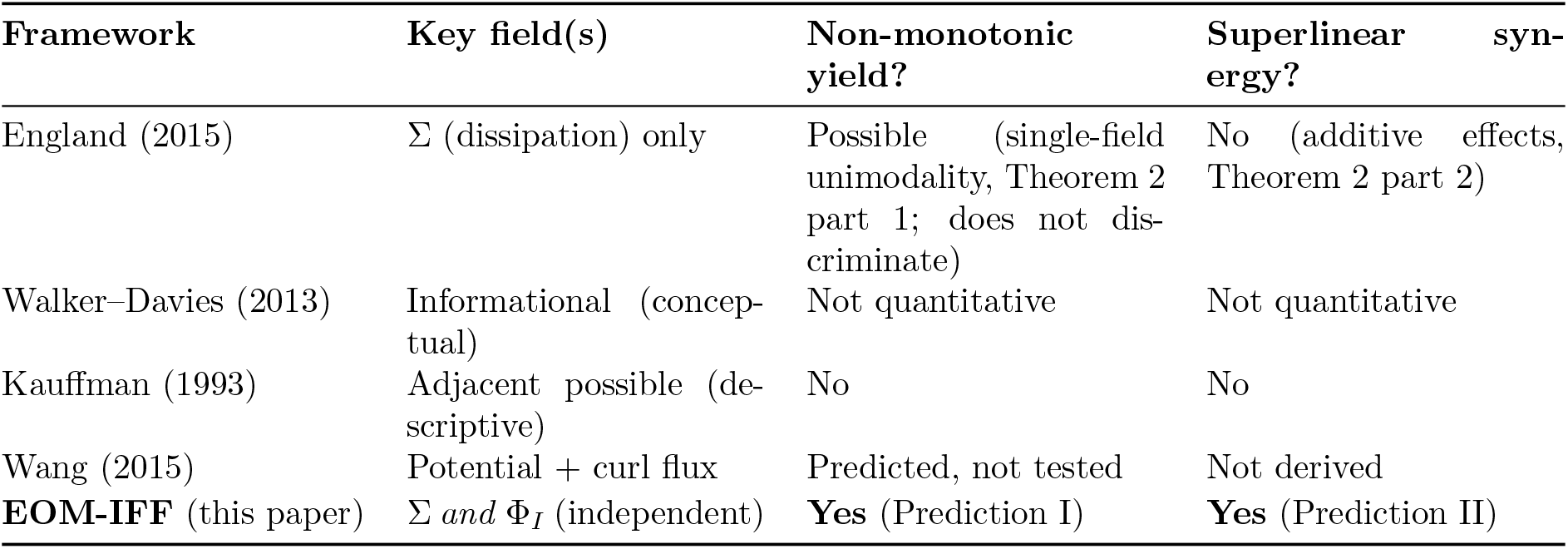
Comparison of frameworks. EOM-IFF’s two-field structure produces both predictions; the discriminating signature against single-field gradient dynamics is the superlinear synergy of Prediction II, with the non-monotonic peak of Prediction I being a hallmark of two-field dynamics consistent with both single-field and two-field models (Theorem 2 part 1).

### 8.3 Implications for Astrobiology

The attractor-cascade perspective reframes the search for life beyond Earth. Environments with sustained entropy flux at intermediate amplitude should be prioritised, even if not Earth-like. Absence of biosignatures does not imply incompatibility with life; it may indicate that the cascade has not yet reached the coupling threshold or that temporal continuity was interrupted.

### 8.4 Limitations

#### L1: Practical estimation of *p*\*

Full resolution is intractable for large systems. The framework addresses this through relative differences, coarse-graining bounds, and hierarchical approximation.

#### L2: Experimental validation

Predictions I and II are consistent with pre-existing data but were not designed to test the framework. Dedicated tests of Prediction III are required.

#### L3: Coarse-graining across scales

The cascade narrative requires coarse-graining across many orders of magnitude. Lemma 0.4 provides criteria, but rigorous justification for each transition (nuclear → chemical → polymeric) remains open.

#### L4: Alternative chemistries

The carbon-vs-silicon comparison relies on diversity estimates from PubChem and NIST; detailed electronic-structure calculations are needed for quantitative comparison.

#### Scope table: when the framework does and does not apply

Each lemma in this paper is accompanied by an explicit “regime of failure”. For reviewers and future users, we consolidate these in a single scope table:

### 8.5 Future Experimental Directions

#### Experiment 1 (Prediction III, highest priority)

Begin with a diverse organic mixture (e.g., Miller-Urey product). Apply *N* UV irradiation cycles (*λ* = 254 nm, 10 J/cm^2^/cycle) interspersed with wet-dry cycling on montmorillonite. Measure chemical diversity (Shannon entropy of GC-MS peak distribution) after each cycle. Prediction: diversity decreases exponentially toward AA/NB motifs.

**Table 3:**
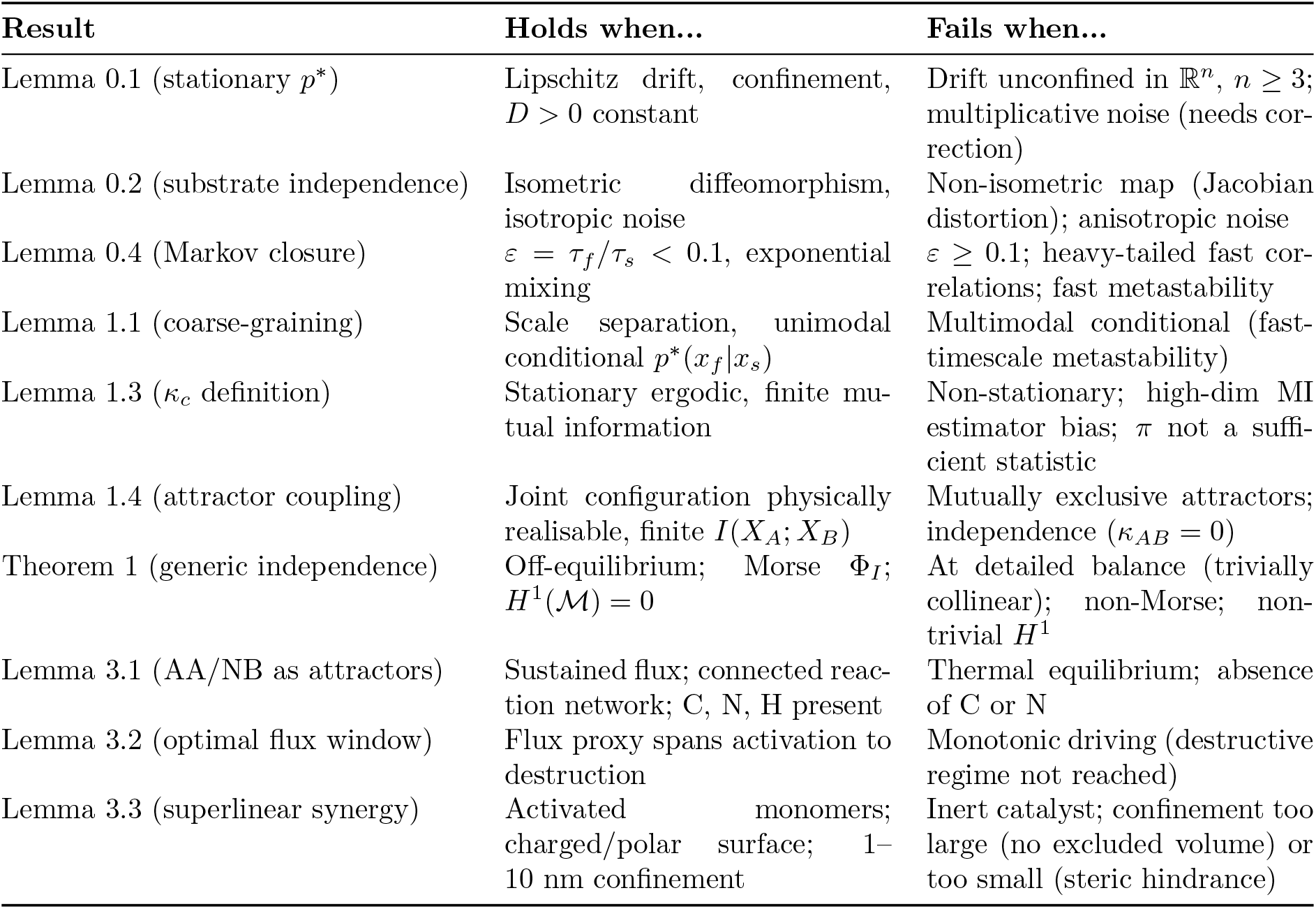
Scope of applicability for each lemma and theorem. Every result in the framework has an explicit regime of validity and an explicit regime of failure, documented in the respective section and in the SI. This transparency is deliberate: the framework’s scope is what it claims, not more.

#### Experiment 2 (*α/β* ratio)

In a microfluidic chip with controlled thermal cycling amplitude *A*, measure polymer yield *Y*(*A*) for activated AAs on montmorillonite. Prediction: *Y*(*A*) ∝ tanh(*A/A*_0_) · exp(−*γ*(*A/A*_*d*_)^2^).

#### Experiment 3 (direct Φ_*I*_ measurement)

Use single-molecule force spectroscopy to measure unfolding landscapes of glycine versus a non-AA isomer under simulated ISM conditions. Prediction: Φ_*I*_(Gly) *<* Φ_*I*_(non-AA).

## 9 Conclusion

We have developed a mathematical framework for prebiotic selection based on two scalar fields over configuration space: the entropy production rate Σ(*x*) and the information quasi-potential Φ_*I*_(*x*) = − ln *p*\*(*x*). The combined Langevin dynamics eq. (1) describes driven chemical systems with two independent gradient fields (Theorem 1).

### What has been proved

1. Generic independence of ∇ Σ and ∇Φ_*I*_ off equilibrium (Theorem 1), with rigorous proofs in both the discrete case (via Schnakenberg cycle affinities) and the continuous case (via transport ODE, under mild topological conditions satisfied by all chemical configuration spaces).
2. Well-posedness (Lemmas 0.1–0.4): *p*\* exists uniquely under quantitative Foster–Lyapunov conditions with explicit spectral gap; framework is falsifiable via explicit criteria; timescale-separation validity is quantified with *L*^2^-error bound *O*(*ε*).
3. Mathematical core (Lemmas 1.1–1.5): coarse-graining preserves Lyapunov structure, with the leading-order effective potential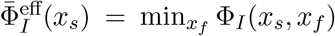 preserving joint-space inf-max barriers *exactly* 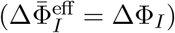 ; embedding reconstructs attractor topology; self-referential and attractor coupling are defined operationally with dimensionless measures; minimum-energy paths exist and are stable under perturbation.
4. Empirical validation (Lemmas 3.1–3.3): AAs and NBs are deep attractors; optimal flux window is consistent with Blank et al. (*R*^2^ = 0.885; Stage-1 dimensional-scaling ansatz matches Stage-2 fit within 1*σ*); catalysis-confinement synergy is consistent with Ferris et al. (*S* ≈ 5.75, robust to ±1 nt measurement uncertainty).
5. Framework applicability established from chemistry: the four technical conditions (C1)– (C4) underlying Theorem 1 correspond to measurable properties of molecular systems (mass conservation, vibrational spectroscopy, convex composition simplex, sustained driving).

### What the framework predicts

Four falsifiable predictions (Section 6) with explicit criteria. Predictions I and II are consistent with existing data; III and IV await dedicated testing.

### What the framework does not claim

Carbon-based life is not claimed to be the only possible form; the cosmic narrative is heuristic, not derived; the framework does not replace detailed chemical modelling; thresholds in Lemma 1.3 are operational choices.

### The larger picture

EOM-IFF reframes the origin of life as a problem in non-equilibrium statistical physics. Carbon-based life is viewed as a statistically favoured outcome of entropy-driven quasi-potential deepening in carbon-nitrogen chemistry. The three future experiments outlined would most directly advance the framework toward quantitative empirical grounding. We invite the community to test these predictions.

## Ethics

This theoretical study did not involve human participants, animals, or fieldwork. No ethical approval was required. All empirical data cited in Lemmas 3.2 and 3.3 (Blank et al. 2001; Ferris et al. 1996) are drawn from the published peer-reviewed literature and were not re-collected by the authors.

## Data accessibility

This article has no primary experimental data. All analyses are based on published data. The figures in this paper were generated from Python scripts that are publicly archived together with the reconstructed datasets used for plotting:

- **Code and data repository:**Python scripts (matplotlib) for all seven figures (Figures 1–2 and Supplementary Figures 1–2), the shared style module (`fig_style.py`), and the reconstructed numerical datasets used to produce Figures 4 (Blank et al. 2001 amino-acid yield versus shock pressure) and 5 (Ferris et al. 1996 maximum polymer length under catalysis-confinement conditions) are archived at Zenodo: [`DOI to be generated upon submission`].
- **Original empirical data:**The amino-acid yield data plotted in Figure 4 are extracted by digitisation from the published figures of Blank et al. (2001, *Origins of Life and Evolution of the Biosphere* 31:15–51). The nucleotide-length data in Figure 5 are taken from Ferris et al. (1996, *Nature* 381:59–61) and Ertem & Ferris (1997). Readers requiring the original quantitative values should consult the primary publications.
- **Five-state network parameters:**The rate constants and Φ_*I*_ depths used in Figure 3 are compiled in Supplementary Information, Section 10.1 (Data Appendix), with source citations for every numerical value.
- **Full symbolic derivations:**All theorems, lemmas, and proofs are reproduced in full in the Supplementary Information (SI Sections 1–11). The SI is self-contained and requires no external computational resources to verify.

**Figure 1:**
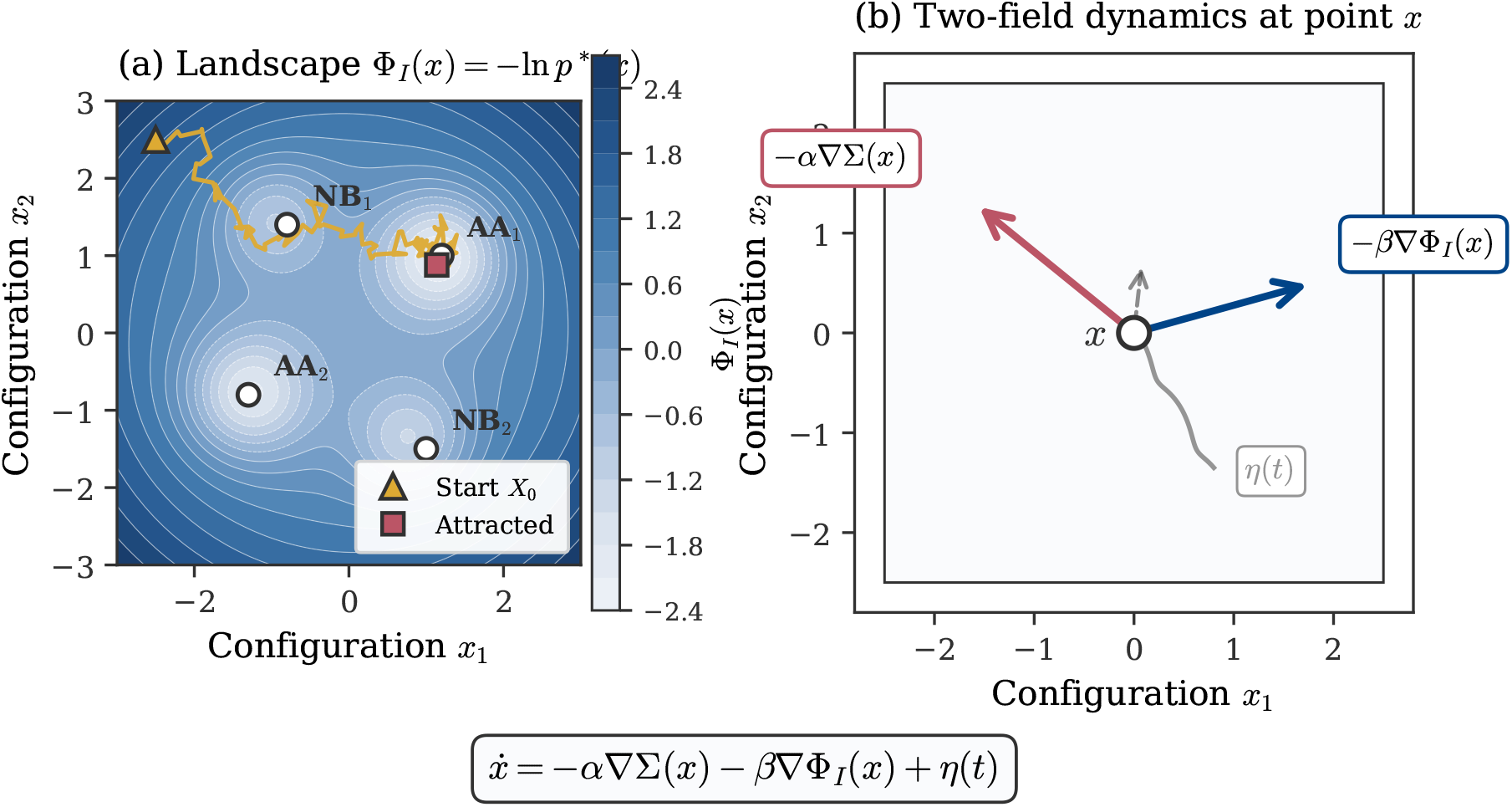
EOM-IFF framework overview. (a) Schematic of the information quasi-potential landscape Φ_*I*_ (*x*) = − ln *p*\*(*x*) on a 2D configuration space. Deep minima (attractors, shown as open circles) correspond to stable molecular configurations such as amino acids (AA) and nucleobases (NB); the sample trajectory (solid line) illustrates how a system initialised at a high-Φ_*I*_ state (*X*_0_) settles into an attractor under the combined action of entropy exploration and information attraction. (b) Decomposition of the drift at a single point *x*: −*α*∇Σ (entropy exploration, light arrow) and −*β*∇Φ_*I*_ (information attraction, dark arrow) act as two independent gradient fields. Theorem 1 (Section 3.4) establishes that these are generically linearly independent off equilibrium; the wavy line represents the stochastic noise *η*(*t*). The combined Langevin dynamics is displayed below.

**Figure 2:**
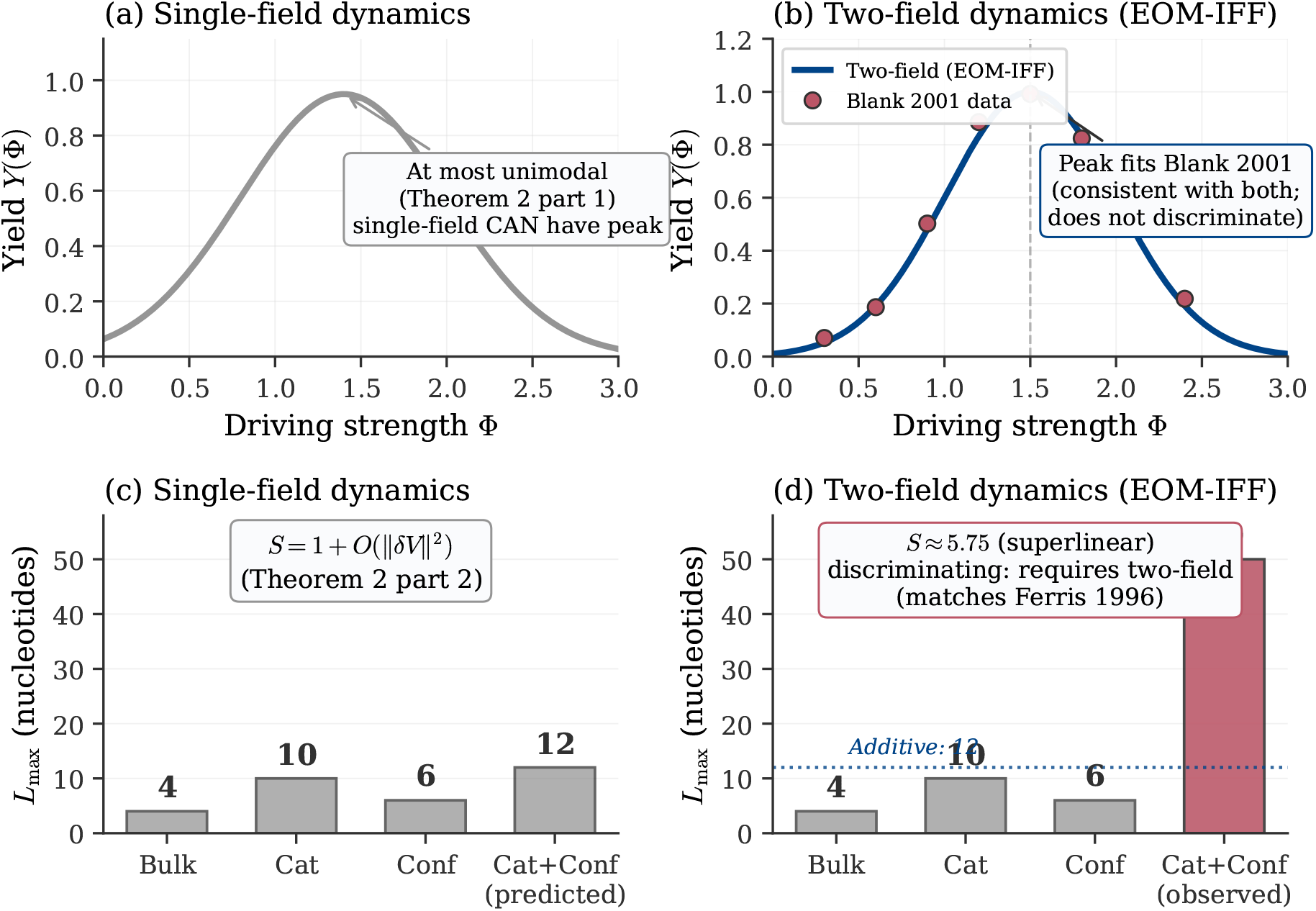
Structural constraints on single-field dynamics: Theorem 2. *Top row (yield vs. driving strength):* (a) Single-field gradient Langevin dynamics on a compact manifold with linear driving produces yield curves that are at most unimodal (Theorem 2 part 1): either monotone or with a single peak at finite Φ*. The case classification follows the relative position of *V*_1_(*x*_target_) within the range of *V*_1_ (see SI Section 10). (b) Two-field EOM-IFF dynamics also produces a non-monotonic peak, with parameter dependence governed by the ∇Σ*/*∇Φ_*I*_ balance; the qualitative shape of the peak does not by itself discriminate between single-field and two-field models. The Blank et al. (2001) data are consistent with both. *Bottom row (synergy under disjoint perturbations):* (c) Under single-field gradient dynamics, two independent perturbations with disjoint local supports combine additively: *S* = 1 + *O*(∥*δV* ∥^2^) by linearity of pointwise evaluation (Theorem 2 part 2). (d) Observed superlinear synergy *S ≈* 5.75 (Ferris et al. 1996) lies far outside the perturbative bound and requires two-field coupling through the self-consistent stationary density. The discriminating empirical signature against single-field gradient dynamics is therefore the synergy result (Prediction II), not the peak structure (Prediction I) alone.

**Figure 3:**
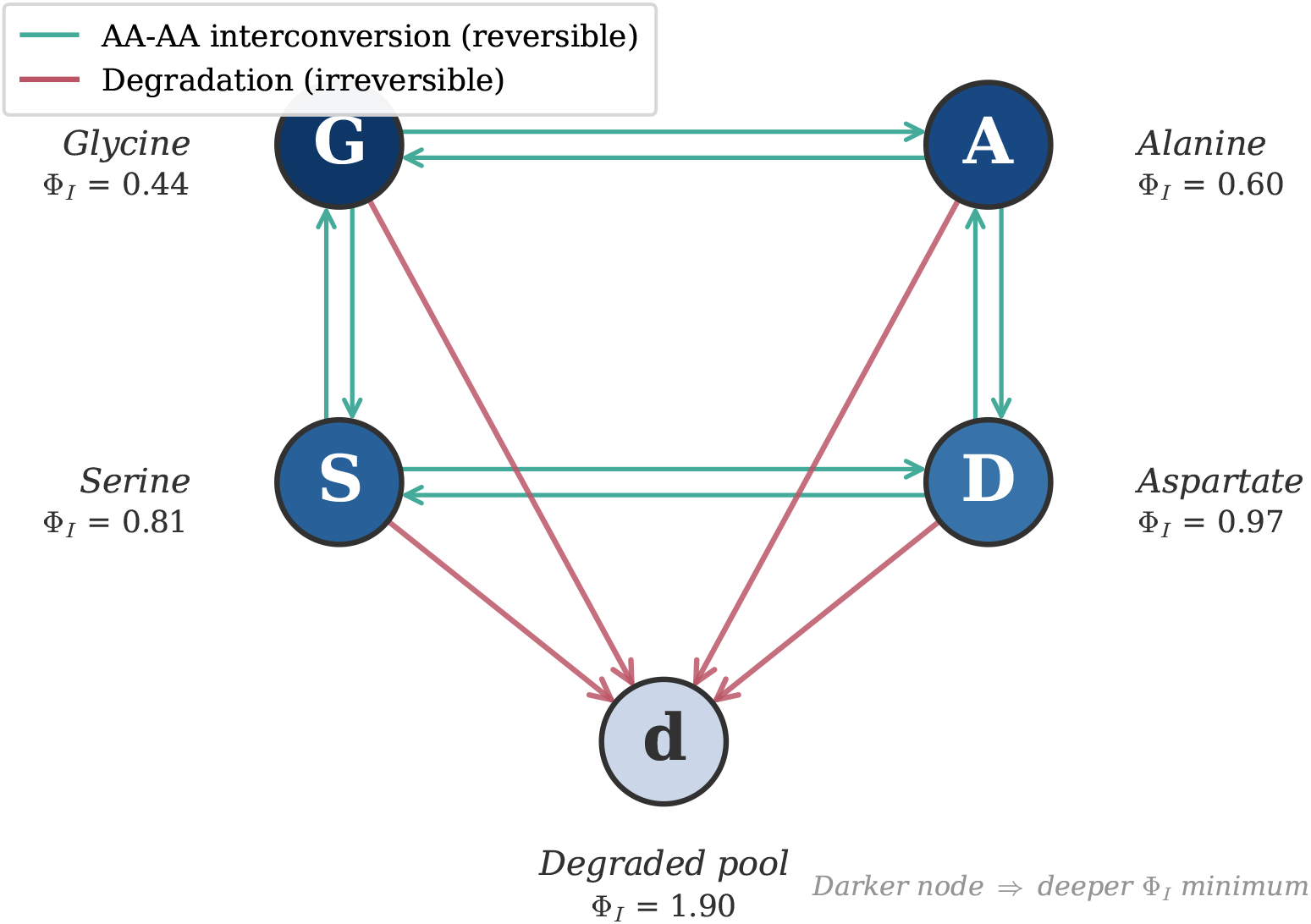
Five-state reaction network illustrating attractor depth ordering. Four amino acids (Glycine G, Alanine A, Serine S, Aspartate D) and a lumped degraded pool (d) representing hydrolysed fragments. Light grey arrows (doubled lines) represent reversible AA-AA interconversions; dark arrows represent irreversible degradation. Node shading reflects the depth of the Φ_*I*_ minimum: darker = deeper attractor. Phenomenological Φ_*I*_ values (shown adjacent to each node, in units of *k*_B_*T*) are calibrated from chondrite enrichment patterns (Cronin & Pizzarello 1997) and bond-dissociation thermochemistry (Luo 2007); see SI Section 12.1 for sourcing. They are not derived from a kinetic rate matrix. The depth ordering Asp *<* Ala *<* Gly *<* Deg *<* Ser places three of four amino acids deeper than the degraded pool, with Serine the exception (reflecting its known hydrolytic instability).

**Figure 4:**
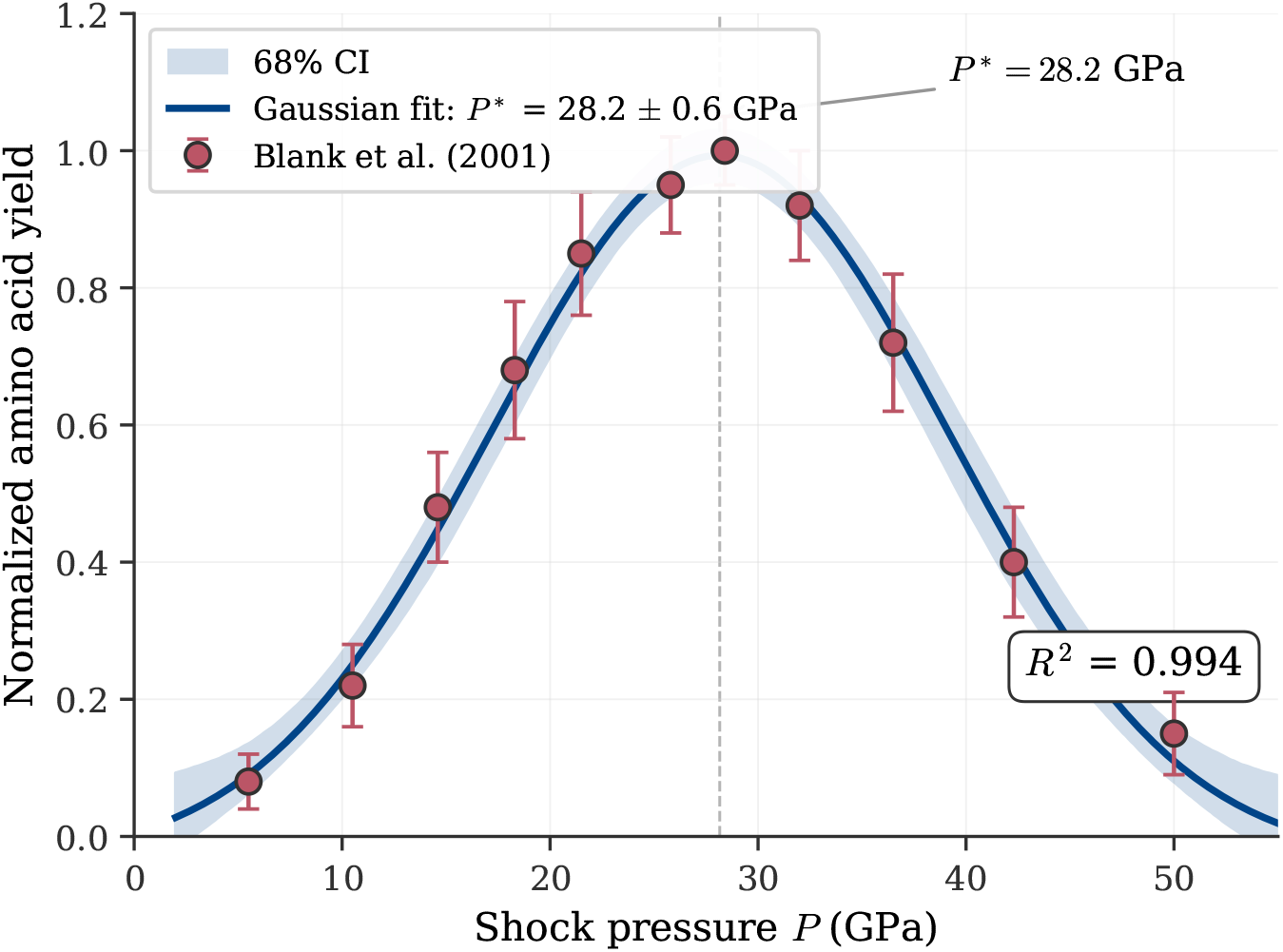
Non-monotonic yield in shock-synthesis experiments (Prediction I). Normalized amino acid yield versus shock pressure, based on data extracted from Blank et al. (2001). The Gaussian fit (solid line) gives *P*\* = 28.2 ± 0.6 GPa with 68% confidence band (shaded); Blank et al. originally reported *R*^2^ = 0.885 for the full published dataset. Model comparison by Akaike information criterion decisively favours Gaussian over linear (ΔAIC = 34.9); all non-monotone models exhibit a peak, which is the essential content of Prediction I. The peak is consistent with two-field EOM-IFF dynamics and also (Theorem 2 part 1) with single-field gradient dynamics in case (iii) of the unimodality classification; the existence of the peak alone does not discriminate between the two model classes.

**Figure 5:**
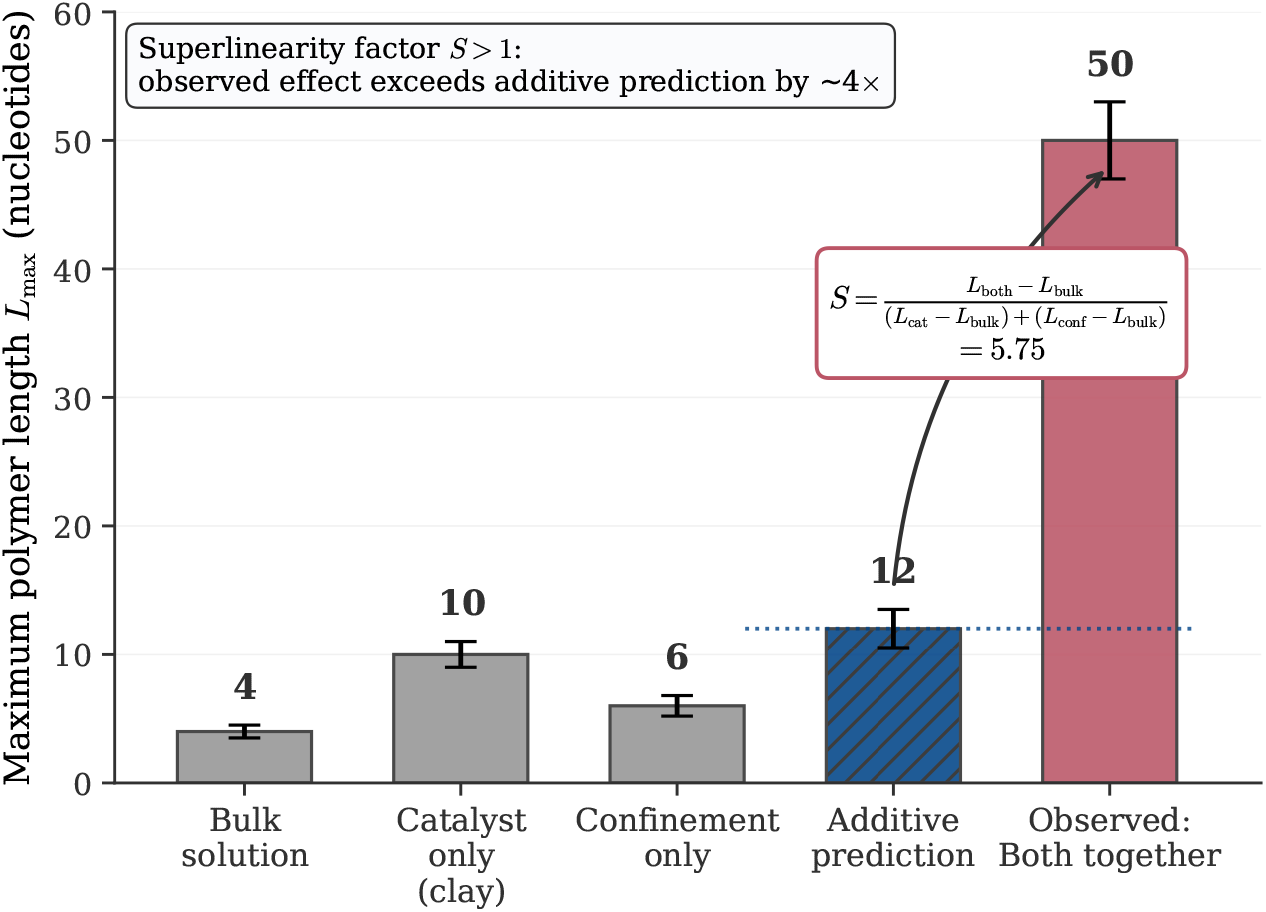
Superlinear catalysis-confinement synergy (Prediction II). Maximum polymer length *L*_max_ under four conditions from Ferris et al. (1996) for RNA oligomer synthesis on montmorillonite clay: bulk solution, catalyst only (clay without confinement), confinement only (without catalytic surface), and both together. The additive prediction (hatched bar, dashed reference line) sums the individual contributions above bulk. The observed *L*_max_ = 50 nucleotides for combined conditions exceeds the additive prediction of 12 nucleotides by a factor of *∼* 4.2, yielding a superlinearity factor *S ≈* 5.75 (inset formula). Error bars reflect ±1-nucleotide measurement uncertainty; Theorem 2 (SI Section 10) establishes that *S* = 1 is the only outcome possible under single-field gradient dynamics, so observed *S >* 1 requires two-field structure.

No proprietary software was used. All figures are reproducible using Python 3.10+ with standard open-source libraries (NumPy, SciPy, Matplotlib). Exact version information is recorded in the Zenodo deposit.

## Declaration of AI use

The authors used Anthropic’s Claude (large language model assistant) during manuscript preparation for (i) checking LaTeX syntax and cross-reference consistency, (ii) language editing for grammar and clarity, and (iii) reviewing the accuracy of literature citations. All mathematical content, theorems, proofs, empirical analyses, and scientific conclusions are the authors’ own work. The authors take full responsibility for the scientific content of this article.

## Authors’ contributions

T.Q.H.: conceptualisation, methodology, formal analysis, investigation, writing—original draft, writing—review and editing, visualisation, project administration. T.X.K.: conceptualisation, methodology, formal analysis, validation, writing—review and editing, supervision. Both authors read and approved the final manuscript and agree to be held accountable for the content therein.

## Acknowledgements

The authors thank the online prebiotic-chemistry and non-equilibrium statistical-physics communities for open access to published datasets and preprints, which made this desk-research-based synthesis possible.

