## Supplement Information for "Prebiotic Selection as a Physical Process: An Information Quasi-Potential Framework for Chemical Convergence"

### Electronic Supplementary Material: Full Rigorous Proofs for "Prebiotic Selection as a Physical Process"

Truong Quynh Hoa
Truong Xuan Khanh
H&K Research Studio, Clevix LLC, Hanoi, Vietnam

April 23, 2026

This document provides the full rigorous proofs and technical background for the main paper. It is structured to be read alongside the main manuscript but is self-contained: each proof can be verified independently given the stated assumptions. Cross-references to the main text use the prefix "Paper" (e.g. Paper Section 2); internal references are without prefix.

#### Contents

**1 Preliminaries and Framework Conventions** **1**

**2 Proof of Lemma 0.1 — Regime Definition Consistency** **2**

**3 Proof of Lemma 0.2 — Minimal Substrate-Independence** **5**

**4 Proof of Lemma 0.4 — Timescale Separation Validity Criterion** **6**

**5 Proof of Lemma 1.1 — Coarse-Graining Preserves Lyapunov Structure** **10**

**6 Proof of Lemma 1.3 — Self-Referential Coupling** **16**

**7 Proof of Lemma 1.4 — Attractor Coupling Criterion** **19**

**8 Proof of Lemma 1.5 — Existence of Minimum-Energy Paths** **21**

**9 Proof of Theorem 1 — Generic Independence of $\nabla\Sigma$ and $\nabla\Phi_I$** **22**

### 1 Preliminaries and Framework Conventions

To avoid definitional circularity identified in earlier drafts, we adopt the following conventions throughout this SI. Let $\mathcal{M}$ be a smooth connected $n$-dimensional Riemannian manifold (configuration space), equipped with a Riemannian metric $g$ and Lebesgue reference measure $\mathrm{d}x$. We consider overdamped Langevin dynamics of the form

$$\mathrm{d}X_t = b(X_t)\,\mathrm{d}t + \sqrt{2D}\,\mathrm{d}W_t, \tag{1}$$

with $W_t$ a standard Wiener process on $\mathcal{M}$, $D > 0$ constant, and drift $b : \mathcal{M} \to T\mathcal{M}$ a Lipschitz vector field. The generator is

$$\mathcal{L} = b \cdot \nabla + D\Delta,$$

with adjoint $\mathcal{L}^\dagger \rho = -\nabla \cdot (b\rho) + D\Delta\rho$ acting on probability densities $\rho$.

**Framework dynamics.** In the EOM-IFF framework, the drift is taken to be

$$b(x) = -\alpha\nabla\Sigma(x) - \beta\nabla\Phi_I(x), \tag{2}$$

where $\alpha, \beta \in \mathbb{R}$ are coupling constants, $\Sigma : \mathcal{M} \to \mathbb{R}$ is the entropy production rate field (defined externally, e.g. via Schnakenberg cycle decomposition of an underlying Markov chain), and $\Phi_I : \mathcal{M} \to \mathbb{R}$ is the information quasi-potential.

**Definition of $\Phi_I$.** We define

$$\Phi_I(x) := -\ln p^*(x), \tag{3}$$

where $p^*$ is the *unique stationary density* of eq. (1) with the full drift eq. (2). This is a *self-consistent* definition: $\Phi_I$ is determined by the full dynamics (including $\Sigma$), and enters back as a drift component. Existence and uniqueness of $p^*$ is established in Section 2.

**Non-circularity.** This differs from the earlier convention $\Phi_I = V/D + \text{const}$ (which would force $\Phi_I \propto \alpha\Sigma + \beta\Phi_I$ and hence collinearity of $\nabla\Sigma$ and $\nabla\Phi_I$, contradicting Theorem 1). Here $\Phi_I$ is a *derived* object (via $-\ln p^*$), not a defining component of the drift potential. The pair $(\Sigma, \Phi_I)$ are functionally related through the stationary Fokker–Planck equation but not algebraically identified.

**Detailed balance.** The system is said to satisfy *detailed balance* if the stationary probability current

$$J^*(x) := b(x)p^*(x) - D\nabla p^*(x)$$

vanishes identically. In this case, $b = D\nabla \ln p^* = -D\nabla\Phi_I$, i.e. $\alpha\nabla\Sigma = -(D-\beta)\nabla\Phi_I$. This yields collinearity of $\nabla\Sigma$ and $\nabla\Phi_I$ everywhere, which is the degenerate case ruled out by Theorem 1.

**Asymptotic regimes.** The framework admits two distinct small parameters that appear in different lemmas, and we make the dependence explicit:

- The *noise parameter* $D > 0$ controls the strength of stochastic fluctuations. The small-noise limit $D \to 0$ is invoked in Lemma 1.1 for Laplace asymptotics of the marginal density $p_s^*(x_s) = \int e^{-\Phi_I/D}\,\mathrm{d}x_f/Z$. The Laplace expansion is uniform on compact subsets bounded away from degenerate critical points (where Hessian eigenvalues approach zero); the relevant uniformity constants are made explicit in the stability condition (Step 2b of Lemma 1.1).
- The *scale-separation parameter* $\varepsilon := \tau_f/\tau_s$ measures the ratio of fast to slow relaxation times. The averaging limit $\varepsilon \to 0$ is invoked in Lemma 0.4 for the Mori–Zwanzig closure with $L^2$ error bound $O(\varepsilon)$ uniform in the time horizon $T$ (with constant $C_T$ growing in $T$). The averaging is performed at *fixed* $D$; no joint scaling between $D$ and $\varepsilon$ is required.

The two limits are taken independently in the lemmas where they appear: Lemma 0.4 fixes $D$ and lets $\varepsilon \to 0$; Lemma 1.1 fixes $\varepsilon$ (or implicitly absorbs it into the conditional density $p^*(x_f|x_s)$) and lets $D \to 0$. No claim of uniformity across the joint $(D,\varepsilon)$-plane is made; the framework conclusions hold in the regime where both parameters are small, each below its respective threshold.

---

#### 2 Proof of Lemma 0.1 — Regime Definition Consistency

**Lemma 1** (Regime definition consistency)**.** *Consider the Langevin SDE eq.* (1) *on $\mathcal{M}$ with Lipschitz drift $b$ and $D > 0$. Suppose:*

*(A1)* ***(Confinement)*** *There exists $R_0 > 0$ and constants $c_1, c_2 > 0$ such that for $\|x\| > R_0$ (in local coordinates, or equivalently, outside a compact set in $\mathcal{M}$):*

$$b(x) \cdot \frac{x}{\|x\|} \le -c_1\|x\| + c_2.$$

*(A2)* ***(Lipschitz regularity)*** *$b \in C^1(\mathcal{M})$ with $\|\nabla b\|_\infty < \infty$ on compact sets, and $\|b(x)\| \le C(1+\|x\|)$ globally.*

*(A3)* ***(Non-degeneracy)*** *$D > 0$.*

*Then eq.* (1) *admits a unique stationary probability measure $\mu^*$ with density $p^* \in C^2(\mathcal{M})$, $p^* > 0$, satisfying the stationary Fokker–Planck equation*

$$-\nabla\cdot(bp^*) + D\Delta p^* = 0. \tag{4}$$

*The quasi-potential $\Phi_I := -\ln p^*$ is well-defined (up to the additive constant $\ln\int e^{\Phi_I}\mathrm{d}x$, fixed by normalization) and of class $C^2$.*

*Proof.* We proceed in four steps.

**Step 1 (Non-explosion and existence of a semigroup).** By (A2), the drift is globally Lipschitz of at most linear growth. Standard SDE theory (Khasminskii 1980, Theorem 3.5; Pavliotis 2014, Section 4.3) guarantees existence and uniqueness of a strong solution $X_t$ to eq. (1) for all $t \geq 0$, non-explosive. The solution defines a Markov semigroup $P_t f(x) := \mathbb{E}_x[f(X_t)]$ acting on bounded measurable $f$, and its adjoint $P_t^\dagger$ acts on probability measures.

**Step 2 (Foster–Lyapunov drift condition with explicit constants).** Define the Lyapunov candidate $V_0(x) := \frac{1}{2}\|x\|^2$ in local coordinates (globally, choose a proper function on $\mathcal{M}$ growing at infinity). The generator acts as

$$\mathcal{L}V_0(x) = b(x) \cdot x + D \cdot n.$$

By assumption (A1), for $\|x\| > R_0$:

$$\mathcal{L}V_0(x) \leq (-c_1\|x\| + c_2) \cdot \|x\| + Dn = -c_1\|x\|^2 + c_2\|x\| + Dn.$$

Completing the square: for $\|x\| \geq R_1 := \max(R_0, 2c_2/c_1)$, we have $c_2\|x\| \leq \frac{c_1}{2}\|x\|^2$, hence $\mathcal{L}V_0 \leq -\frac{c_1}{2}\|x\|^2 + Dn$. Combining with the compact-set bound (where $\mathcal{L}V_0$ is bounded by a constant $M_0 < \infty$ from continuity), we obtain the drift condition with explicit constants:

$$\mathcal{L}V_0(x) \leq -\gamma V_0(x) + K, \quad \gamma = c_1, \quad K = Dn + M_0 + c_1 R_1^2/2, \tag{5}$$

valid for all $x \in \mathcal{M}$.

**Step 2b (Consequence: Poincaré and log-Sobolev inequalities).** The Poincaré inequality for the stationary density $p^*$ requires either (i) the Bakry–Émery criterion (uniform convexity of $V := -D \ln p^*$, with $\text{Hess}(V) \succeq \rho I$ for some $\rho > 0$, giving Poincaré constant $\rho/D$ via Bakry–Émery 1985, Bakry–Gentil–Ledoux 2014), or (ii) the Lyapunov-with-Poincaré route (Cattiaux–Guillin 2009; Bakry–Cattiaux–Guillin 2008): the Foster–Lyapunov drift condition eq. (5) together with a local Poincaré inequality on bounded sets (which holds automatically by the elliptic regularity of Step 3 below) implies a global Poincaré inequality

$$\text{Var}_{\mu^*}(f) \leq \frac{1}{\lambda_1}\, \mathbb{E}_{\mu^*}\left[\|\nabla f\|^2\right] \quad \forall f \in H^1(\mu^*), \tag{6}$$

with spectral gap $\lambda_1 > 0$ (the explicit value depends on the local Poincaré constants and on $\gamma, K$ via the Cattiaux–Guillin formula; the key point is positivity).

For the chemistry application of the EOM-IFF framework, route (ii) is the relevant one: the drift condition (A1) is a coarse confinement assumption that holds under any reasonable potential, while uniform convexity of $V$ would be overly restrictive (it would forbid the multi-well landscapes that define the attractor structure). Consequence: the Markov semigroup converges to stationarity exponentially with rate $\lambda_1$:

$$\|P_t\nu - \mu^*\|_{TV} \leq C\, e^{-\lambda_1 t/2}, \qquad t \geq 0,$$

for any initial measure $\nu$ with $\nu \ll \mu^*$ and $\mathrm{d}\nu/\mathrm{d}\mu^* \in L^2(\mu^*)$ (constant $C$ depends on this density). This is the *quantitative* version of positive Harris recurrence.

*Remark on the log-Sobolev inequality.* The log-Sobolev inequality, with constant $\rho/D$, holds under the stronger Bakry–Émery condition $\text{Hess}(V) \succeq \rho I$. For multi-well landscapes typical of prebiotic chemistry, this condition fails between basins; we therefore do not invoke log-Sobolev in subsequent arguments and rely only on the Poincaré inequality eq. (6) obtained via the Lyapunov-with-Poincaré route.

**Step 3 (Smoothness and positivity via elliptic regularity).** The generator $\mathcal{L} = b \cdot \nabla + D\Delta$ is *uniformly elliptic*: its principal symbol $D\|\xi\|^2$ is bounded below by $D\|\xi\|^2$ with $D > 0$. By Hörmander's hypoellipticity theorem (immediate since the generator is already elliptic, no commutator bracket needed) and Schauder interior estimates (Gilbarg–Trudinger 2001, Theorem 6.17), any weak solution $p^*$ of the stationary Fokker–Planck equation eq. (4) with $p^* \in L^1(\mathcal{M})$ satisfies $p^* \in C^\infty(\mathcal{M})$. Applying the strong maximum principle (Gilbarg–Trudinger Theorem 3.5) to $p^* \geq 0$ satisfying $-\nabla \cdot (bp^*) + D\Delta p^* = 0$: either $p^* \equiv 0$ (ruled out by $\int p^* = 1$) or $p^* > 0$ everywhere.

**Step 4 (Uniqueness via spectral gap).** Suppose $\mu_1^*, \mu_2^*$ are two stationary probability measures of the semigroup $P_t$. Both are absolutely continuous with smooth positive densities $p_1^*, p_2^*$ by Step 3. Define the Radon–Nikodym derivative $f := \mathrm{d}\mu_1^*/\mathrm{d}\mu_2^* = p_1^*/p_2^* \in L^2(\mu_2^*)$ (positive, with $\mathbb{E}_{\mu_2^*}[f] = 1$).

Stationarity of $\mu_1^*$ means $P_t^\dagger \mu_1^* = \mu_1^*$ for all $t \geq 0$. Equivalently, in the function-side action, the dual of this statement is $P_t f = f$ in $L^2(\mu_2^*)$ for all $t \geq 0$ — i.e., $f$ is a fixed point of the Markov semigroup acting on the function space $L^2(\mu_2^*)$.

By the Poincaré inequality eq. (6) (Step 2b), the symmetric part of the generator has spectral gap $\lambda_1 > 0$ on the subspace of zero-mean functions in $L^2(\mu_2^*)$.

*Why the spectral gap holds for $\mu_2^*$.* The Lyapunov-with-Poincaré argument used in Step 2b (route (ii)) derives the Poincaré inequality from the drift condition eq. (5) together with local Poincaré inequalities on bounded sets, both of which are properties of the generator $\mathcal{L}$ alone and do not depend on which stationary measure is used as reference. Concretely, for *any* stationary probability measure $\mu^*$ of the semigroup — be it $\mu_1^*$ or $\mu_2^*$ — the Cattiaux–Guillin argument applied with $\mu^*$ as reference yields $\mathrm{Var}_{\mu^*}(f) \leq \lambda_1^{-1} \mathbb{E}_{\mu^*}[\|\nabla f\|^2]$ with $\lambda_1 > 0$. Hence the spectral gap on $L^2(\mu_2^*)$ is established a priori, without circularly assuming $\mu_1^* = \mu_2^*$.

The only $L^2(\mu_2^*)$ fixed points of $P_t$ are therefore constants (the kernel of the generator is the constants). Since $\mathbb{E}_{\mu_2^*}[f] = 1$, we have $f \equiv 1$ $\mu_2^*$-almost everywhere, i.e. $p_1^* = p_2^*$.

*Remark.* The argument uses only the existence of a spectral gap on the zero-mean subspace, which follows from the drift condition eq. (5) together with the local Poincaré inequality from elliptic regularity (Step 3). An alternative formulation via the standard Doeblin/minorisation route (Meyn & Tweedie 2009, Chapter 16) gives the same conclusion under the same hypotheses without requiring the spectral-gap machinery.

**Remark on redundancy.** The Meyn–Tweedie approach (positive Harris recurrence) and the spectral-gap approach give the *same* conclusion via different techniques. We present both because peer reviewers from probability and PDE communities often prefer different methods; the unified statement is that existence and uniqueness of $\mu^*$ hold with quantitative convergence rate $\lambda_1 > 0$.

This completes the proof. □

**Regime of validity.** $D > 0$ finite; autonomous dynamics; $\mathcal{M}$ of finite dimension; drift Lipschitz with linear growth and radial confinement (A1).

**Regime of failure.** *Loss of confinement*: if (A1) fails (e.g. $b \equiv 0$ on $\mathbb{R}^n$), the process is recurrent only in $n \leq 2$ and has no finite invariant measure in $n \geq 3$. *Multiplicative noise*: if $D = D(x)$, eq. (4) is replaced by $-\nabla \cdot (bp^*) + \nabla^2 : (Dp^*) = 0$ (Itô convention); stationary density is then $p^* \propto D^{-1} e^{-\tilde{V}/D}$ for an appropriately modified $\tilde{V}$, and $\Phi_I = -\ln p^*$ acquires a Jacobian correction. *Time-dependent $b$*: no single stationary density; one must work with a time-periodic or pathwise-ergodic measure.

**Connection to main paper.** Under Assumptions (A1)–(A3), the central object $\Phi_I = -\ln p^*$ of the EOM-IFF framework is well-defined and smooth. In prebiotic chemistry, (A1) corresponds to mass conservation (bounded total concentration), (A2) to bounded reaction rates, (A3) to non-zero thermal noise.

---

#### 3 Proof of Lemma 0.2 — Minimal Substrate-Independence

**Lemma 2** (Substrate-independence under isometric diffeomorphism)**.** *Let* $(\mathcal{M}_1, g_1)$ *and* $(\mathcal{M}_2, g_2)$ *be Riemannian manifolds, and let* $\varphi : \mathcal{M}_1 \to \mathcal{M}_2$ *be a* $C^2$-isometric *diffeomorphism (i.e.* $\varphi^* g_2 = g_1$*). Consider two Langevin systems:*

$$dX_t^{(1)} = b_1(X_t^{(1)})dt + \sqrt{2D}\, dW_t^{(1)}, \qquad dX_t^{(2)} = b_2(X_t^{(2)})dt + \sqrt{2D}\, dW_t^{(2)},$$

*with* $b_2 = \varphi_* b_1 := D\varphi \circ b_1 \circ \varphi^{-1}$ *(pushforward vector field). Then:*

1. *The stationary densities are related by* $p_2^*(y) = p_1^*(\varphi^{-1}(y))$ *(pointwise equality, no Jacobian).*
2. *The quasi-potentials are related by* $\Phi_I^{(2)}(y) = \Phi_I^{(1)}(\varphi^{-1}(y))$*.*
3. *If* $x^*$ *is a local minimum of* $\Phi_I^{(1)}$ *with basin of attraction* $B_1 \subseteq \mathcal{M}_1$*, then* $\varphi(x^*)$ *is a local minimum of* $\Phi_I^{(2)}$ *with basin* $B_2 = \varphi(B_1)$*. Barrier heights are preserved:* $\Delta\Phi_I^{(1)} = \Delta\Phi_I^{(2)}$*.*

**Step 1 (Transformation of the SDE under isometry).** Let $Y_t := \varphi(X_t^{(1)})$. By Itô's formula on manifolds (Hsu 2002, Theorem 3.3.1), since $\varphi$ is a smooth isometry,

$$dY_t = D\varphi(X_t^{(1)}) \cdot b_1(X_t^{(1)})dt + \sqrt{2D}\, D\varphi(X_t^{(1)})dW_t^{(1)}.$$

The drift term matches $b_2(Y_t)$ by definition of pushforward. For the noise term: because $\varphi$ is an isometry, $D\varphi$ is orthogonal at each point, so $D\varphi\, dW_t^{(1)}$ is again a standard Wiener process $dW_t^{(2)}$ on $\mathcal{M}_2$ by Lévy's characterization (Karatzas–Shreve 1991, Theorem 3.3.16). Hence $Y_t$ satisfies the second Langevin SDE in law. The change-of-variables formula for densities under a diffeomorphism gives $p_2^*(y) = p_1^*(\varphi^{-1}(y)) \cdot |\det J_{\varphi^{-1}}(y)|$. For an *isometric* diffeomorphism, $|\det J_\varphi| = 1$ everywhere (preservation of Riemannian volume), so the Jacobian term equals 1 and we have

$$p_2^*(y) = p_1^*(\varphi^{-1}(y)).$$

Taking $-\ln$ gives the relation for $\Phi_I$. Since $\varphi$ is a homeomorphism and $\Phi_I^{(2)} = \Phi_I^{(1)} \circ \varphi^{-1}$, the sublevel sets satisfy

$$\{y \in \mathcal{M}_2 : \Phi_I^{(2)}(y) < c\} = \varphi\big(\{x \in \mathcal{M}_1 : \Phi_I^{(1)}(x) < c\}\big).$$

Hence critical points, basins, and saddle configurations are mapped bijectively. For any two critical points $x_1^*, x_2^*$ joined by a minimum-energy path $\gamma$ in $\mathcal{M}_1$, the barrier height is

$$\Delta\Phi_I^{(1)} = \max_{t\in[0,1]} \Phi_I^{(1)}(\gamma(t)) - \Phi_I^{(1)}(x_1^*),$$

and the pushed-forward path $\varphi \circ \gamma$ gives the identical value on $\mathcal{M}_2$: $\Delta\Phi_I^{(2)} = \Delta\Phi_I^{(1)}$. □

**Regime of validity.** $\varphi$ is a $C^2$ *isometric* diffeomorphism; noise is additive and isotropic with constant amplitude $D$.

**Regime of failure.** *Non-isometric diffeomorphism*: if $|\det J_\varphi|$ varies, the Jacobian contributes an additive non-constant term $\ln|\det J_{\varphi^{-1}}|$ to $\Phi_I^{(2)}$. Basin *topology* (which points flow to which attractor) is still preserved (homeomorphism), but relative barrier heights distort. *Anisotropic noise*: if the diffusion tensor is not proportional to the metric, the pushforward creates a non-Markovian or multiplicative-noise structure; a separate lemma applies. *Projection or embedding* (non-bijective $\varphi$): basins may merge or split; Lemma fails.

**Remark** (What this lemma does and does not say)**.** *This lemma establishes a* mathematical *invariance: any two dynamical systems that are dynamically equivalent under a smooth isometry share the same attractor partition. It does* not *claim that carbon-based and silicon-based chemistry are dynamically equivalent (they are not related by any explicit diffeomorphism). Substrate-independence* in principle *means only that EOM-IFF applies wherever an appropriate Langevin description exists; the empirical hypothesis that biological and silicon chemistries are both governed by EOM-IFF is separate and discussed in Section 7.3 of the main paper.*

---

### 4 Proof of Lemma 0.4 — Timescale Separation Validity Criterion

**Lemma 3** (Markovian coarse-graining under timescale separation)**.** *Consider a Langevin system with slow variables $x_s \in \mathbb{R}^{n_s}$ and fast variables $x_f \in \mathbb{R}^{n_f}$:*

$$\mathrm{d}x_s = f_s(x_s, x_f)\,\mathrm{d}t + \sqrt{2D_s}\,\mathrm{d}W_s, \tag{7a}$$

$$\mathrm{d}x_f = \frac{1}{\varepsilon} f_f(x_s, x_f)\,\mathrm{d}t + \sqrt{\tfrac{2D_f}{\varepsilon}}\,\mathrm{d}W_f, \tag{7b}$$

*where $\varepsilon > 0$ is a small parameter measuring scale separation, and $f_s, f_f$ Lipschitz. Assume:*

*(B1) For each fixed $x_s$, the* frozen fast SDE $\mathrm{d}\xi = f_f(x_s,\xi)\mathrm{d}t + \sqrt{2D_f}\mathrm{d}W$ *admits a unique invariant measure $\rho_f(\cdot|x_s)$ with exponential mixing: $\|P_t^{(x_s)}\nu - \rho_f(\cdot|x_s)\|_{TV} \le Ce^{-t/\tau_f}$ for all initial $\nu$.*

*(B2) The mean drift $\bar{f}_s(x_s) := \int f_s(x_s, x_f)\rho_f(x_f|x_s)\mathrm{d}x_f$ is Lipschitz in $x_s$.*

*(B3) (Used in Lemma 1.1.) The conditional density $p^*(x_f|x_s)$ is unimodal in $x_f$ for each $x_s$, with a unique minimum $x_f^*(x_s)$ of $\Phi_I(x_s,\cdot)$ that depends $C^1$-smoothly on $x_s$. This regularity assumption is automatic when the conditional fast generator $\mathcal{L}_1$ has a strictly convex effective potential, but is imposed as an explicit hypothesis where used in coarse-graining arguments below.*

*Then the slow process $x_s^\varepsilon$ converges as $\varepsilon \to 0$, in the sense of weak convergence in $C([0,T];\mathbb{R}^{n_s})$, to the solution of the* averaged *SDE*

$$\mathrm{d}\bar{x}_s = \bar{f}_s(\bar{x}_s)\,\mathrm{d}t + \sqrt{2\bar{D}_s(\bar{x}_s)}\,\mathrm{d}\bar{W}, \tag{8}$$

*where $\bar{D}_s$ is determined by a Green–Kubo-type integral of fluctuations (see eq.* (16) *below). The $L^2$-error over a finite time horizon $[0,T]$ satisfies*

$$\sup_{t\in[0,T]} \mathbb{E}\big[\|x_s^\varepsilon(t) - \bar{x}_s(t)\|^2\big] \le C_T \cdot \varepsilon, \tag{9}$$

*with $C_T = 2T\,C_R\,e^{L_s T}$ depending on the time horizon $T$, the Lipschitz constant $L_s$ of the averaged drift, and a constant $C_R$ that depends on $\|\nabla^2 f_s\|_{L^\infty}$ and on $\tau_f$ (the fast mixing time, entering through the bound on the corrector $\phi_1$ in Step 3 below). The product $C_T \cdot \varepsilon$ is therefore $O(\tau_f^2/\tau_s)$ in absolute time units, vanishing as the scale separation improves.*

*Proof.* We apply the standard Kurtz–Papanicolaou averaging framework (Pavliotis–Stuart 2008, Chapter 11; Kurtz 1992).

**Step 1 (Mori–Zwanzig projection: explicit operator algebra).** The generator of eq. (7) in rescaled form (with $\mathrm{d}x_f$ fast equation scaled by $\varepsilon^{-1}$) reads

$$\mathcal{L}^\varepsilon = \mathcal{L}_0 + \varepsilon^{-1}\mathcal{L}_1, \qquad \mathcal{L}_0 = f_s \cdot \nabla_s + D_s\Delta_s, \qquad \mathcal{L}_1 = f_f \cdot \nabla_f + D_f\Delta_f.$$

$\mathcal{L}_0$ acts on slow variables only; $\mathcal{L}_1$ acts on fast variables with $x_s$ treated as a frozen parameter.

Define the Mori projection operator $\mathcal{P} : L^2(\mu^\varepsilon) \to L^2(\mu^\varepsilon)$ by

$$(\mathcal{P}g)(x_s, x_f) := \int g(x_s, x'_f)\,\rho_f(x'_f|x_s)\,\mathrm{d}x'_f, \tag{10}$$

where $\rho_f(\cdot|x_s)$ is the invariant measure of $\mathcal{L}_1$ with $x_s$ frozen. By construction, $\mathcal{P}$ is idempotent ($\mathcal{P}^2 = \mathcal{P}$), self-adjoint in $L^2(\mu^\varepsilon)$, and $\mathcal{P}f = \bar{f}$ for $\bar{f}$ depending only on $x_s$. Let $\mathcal{Q} := I - \mathcal{P}$ denote the orthogonal complement.

*Key identity (projection of the fast generator):*

$$\mathcal{P}\mathcal{L}_1 = 0, \qquad \mathcal{L}_1\mathcal{P} = 0. \tag{11}$$

*Justification.* The second identity $\mathcal{L}_1\mathcal{P} = 0$ is immediate: $\mathcal{P}g$ depends only on $x_s$ (the integral over $x_f$ removes $x_f$-dependence), and $\mathcal{L}_1$ contains only derivatives in $x_f$, so $\mathcal{L}_1$ annihilates any function of $x_s$ alone.

The first identity $\mathcal{P}\mathcal{L}_1 = 0$ requires the invariance condition $\mathcal{L}_1^\dagger\rho_f = 0$, which holds by definition of $\rho_f(\cdot|x_s)$ as the invariant measure of $\mathcal{L}_1$ (assumption (B1)). Explicitly, for any sufficiently smooth test function $g(x_s, x_f)$,

$$(\mathcal{P}\mathcal{L}_1 g)(x_s) = \int (\mathcal{L}_1 g)(x_s, x'_f)\,\rho_f(x'_f|x_s)\,\mathrm{d}x'_f = \int g(x_s, x'_f)\,\big(\mathcal{L}_1^\dagger\rho_f\big)(x'_f|x_s)\,\mathrm{d}x'_f = 0,$$

where the second equality is integration by parts in $x_f$ (boundary terms vanish under (B1) and the assumed decay of $\rho_f$). Hence $\mathcal{P}\mathcal{L}^\varepsilon = \mathcal{P}\mathcal{L}_0 = \mathcal{P}\mathcal{L}_0\mathcal{P} + \mathcal{P}\mathcal{L}_0\mathcal{Q}$.

**Step 1b (Dyson expansion and memory kernel).** Applying $\mathcal{P}$ to the Fokker–Planck equation $\partial_t\rho^\varepsilon = (\mathcal{L}^\varepsilon)^\dagger\rho^\varepsilon$, using eq. (11), yields the *exact* Mori–Zwanzig equation (Zwanzig 1961; Grabert 1982):

$$\partial_t(\mathcal{P}\rho^\varepsilon) = \mathcal{P}\mathcal{L}_0^\dagger\mathcal{P}\rho^\varepsilon + \int_0^t \mathcal{K}(t-s)^\dagger(\mathcal{P}\rho^\varepsilon)(s)\,\mathrm{d}s + \Xi^\varepsilon(t), \tag{12}$$

$$\mathcal{K}(t) := \mathcal{P}\mathcal{L}_0\mathcal{Q}\,e^{t\mathcal{Q}\mathcal{L}^\varepsilon\mathcal{Q}}\,\mathcal{Q}\mathcal{L}_0\mathcal{P}, \tag{13}$$

where $\mathcal{K}(t)$ is the memory kernel and $\Xi^\varepsilon(t) = \mathcal{Q}e^{t\mathcal{L}^\varepsilon}\mathcal{Q}\rho^\varepsilon(0)$ is the noise term (orthogonal fluctuations).

*Decay rate of the memory kernel.* The decay of $\mathcal{K}(t)$ is controlled by the spectrum of $\mathcal{Q}\mathcal{L}^\varepsilon\mathcal{Q}$ on $\mathrm{Range}(\mathcal{Q}) \subset L^2(\mu^\varepsilon)$. Decompose

$$\mathcal{Q}\mathcal{L}^\varepsilon\mathcal{Q} = \varepsilon^{-1}\,\mathcal{Q}\mathcal{L}_1\mathcal{Q} \;+\; \mathcal{Q}\mathcal{L}_0\mathcal{Q}.$$

By assumption (B1), the frozen fast semigroup $e^{t\mathcal{L}_1}$ on $L^2(\rho_f(\cdot|x_s))$ has spectral gap $1/\tau_f > 0$ uniformly in $x_s$ (exponential mixing). The projection $\mathcal{Q}$ orthogonalises against the zero-eigenspace of $\mathcal{L}_1$ (which consists of functions of $x_s$ only, the kernel of $\mathcal{L}_1$), so $\mathcal{Q}\mathcal{L}_1\mathcal{Q}$ has spectrum in $\{\Re z \leq -1/\tau_f\}$. The rescaled operator $\varepsilon^{-1}\mathcal{Q}\mathcal{L}_1\mathcal{Q}$ therefore has spectral gap $1/(\varepsilon\tau_f)$. The perturbation $\mathcal{Q}\mathcal{L}_0\mathcal{Q}$ is bounded by $\|\mathcal{L}_0\| \leq C_0(\|f_s\|_{C^1} + D_s)$, finite by Lipschitz regularity of $f_s$. For $\varepsilon$ sufficiently small that $\varepsilon C_0 < 1/(2\tau_f)$, standard perturbation theory (Kato 1995, Theorem IV.3.18) preserves a spectral gap of at least $1/(2\varepsilon\tau_f)$. Consequently the operator semigroup contracts as

$$\left\|e^{t\mathcal{Q}\mathcal{L}^\varepsilon\mathcal{Q}}\right\|_{L^2(\mu^\varepsilon)\to L^2(\mu^\varepsilon)} \leq e^{-t/(2\varepsilon\tau_f)}, \qquad \|\mathcal{K}(t)\|_{L^2(\mu^\varepsilon)} \leq C_1\, e^{-t/(2\varepsilon\tau_f)}, \tag{14}$$

where $C_1 = \|\mathcal{P}\mathcal{L}_0\mathcal{Q}\| \cdot \|\mathcal{Q}\mathcal{L}_0\mathcal{P}\| \leq C_0^2$. The factor $1/(2\varepsilon\tau_f)$ in the exponent (rather than the naive $1/(\varepsilon\tau_f)$) reflects the perturbative correction; for the leading-order analysis of Step 2, only the order of magnitude $1/(\varepsilon\tau_f)$ matters.

**Step 2 (Markovian closure in the limit $\varepsilon \to 0$, with explicit error).** By eq. (14), the memory kernel decays on the fast timescale $\varepsilon\tau_f$, while $\mathcal{P}\rho^\varepsilon$ evolves on the slow scale $\tau_s = O(1)$. Expanding the convolution for smooth $\mathcal{P}\rho^\varepsilon$:

$$\int_0^t \mathcal{K}(t-s)^\dagger(\mathcal{P}\rho^\varepsilon)(s)\,\mathrm{d}s = \Big[\int_0^\infty \mathcal{K}(u)^\dagger\,\mathrm{d}u\Big](\mathcal{P}\rho^\varepsilon)(t) + O(\varepsilon\tau_f/\tau_s) = \mathcal{K}_\infty(\mathcal{P}\rho^\varepsilon)(t) + O(\varepsilon).$$

Here $\mathcal{K}_\infty := \int_0^\infty \mathcal{K}(u)\,\mathrm{d}u$ is the Markov closure operator. Substituting into eq. (12):

$$\partial_t(\mathcal{P}\rho^\varepsilon) = \big[\mathcal{P}\mathcal{L}_0^\dagger\mathcal{P} + \mathcal{K}_\infty^\dagger\big](\mathcal{P}\rho^\varepsilon) + O(\varepsilon) + \Xi^\varepsilon(t).$$

Identifying the effective drift and diffusion:

$$\bar{f}_s(x_s) = (\mathcal{P}f_s)(x_s) = \int f_s(x_s, x_f)\rho_f(x_f|x_s)\,\mathrm{d}x_f, \tag{15a}$$

$$\bar{D}_s(x_s) = D_s I + \mathcal{D}_{GK}^{\mathrm{sym}}(x_s), \tag{15b}$$

where the Green–Kubo correction (derived from the memory kernel) is

$$\mathcal{D}_{GK}(x_s) = \int_0^\infty \mathbb{E}_{\rho_f(\cdot|x_s)}\big[\delta f_s(x_s,\xi_t) \otimes \delta f_s(x_s,\xi_0)\big]\,\mathrm{d}t, \tag{16}$$

with $\delta f_s := f_s - \bar{f}_s$ and $\xi_t$ the frozen fast process, and $\mathcal{D}_{GK}^{\mathrm{sym}} := \frac{1}{2}(\mathcal{D}_{GK} + \mathcal{D}_{GK}^T)$ denotes its symmetric part. The symmetric part is positive semidefinite: by stationarity of the fast process, the symmetrisation yields the Fourier transform of the autocorrelation function at zero frequency,

$$\xi^T\mathcal{D}_{GK}^{\mathrm{sym}}(x_s)\,\xi = \tfrac{1}{2}\int_{-\infty}^\infty \mathbb{E}_{\rho_f(\cdot|x_s)}\big[(\xi\cdot\delta f_s)(t)\,(\xi\cdot\delta f_s)(0)\big]\,\mathrm{d}t = \tfrac{1}{2}S_g(0) \geq 0,$$

where $S_g$ is the spectral density of $g := \xi \cdot \delta f_s$ (Wiener–Khinchin/Bochner). The antisymmetric part of $\mathcal{D}_{GK}$ contributes only a divergence-free drift correction (vanishing under $\nabla\cdot(D\nabla\bullet)$) and is absorbed into the effective drift; the diffusion coefficient $\bar{D}_s$ retains only the symmetric part.

**Remark** (State dependence of $\bar{D}_s$). $\bar{D}_s(x_s)$ *is* state-dependent *even when $D_s$ and $D_f$ are constants, because $\mathcal{D}_{GK}$ depends on $x_s$ through the fast invariant measure $\rho_f(\cdot|x_s)$ and the fluctuation $\delta f_s$. This state dependence is typically mild for most prebiotic chemistry (rates vary smoothly with composition) but can become important near phase transitions.*

**Remark** (Fluctuation-dissipation consistency)**.** *The effective diffusion eq.* (15)*–eq.* (16) *satisfies the fluctuation-dissipation theorem in the following sense. Let $\bar{\mathcal{L}}$ denote the generator of the averaged dynamics eq.* (8)*. A direct computation (Pavliotis–Stuart 2008, Theorem 11.4) shows that for any observable $\phi \in C^2(\mathbb{R}^{n_s})$ in the domain of $\bar{\mathcal{L}}$:*

$$\lim_{T\to\infty} \frac{1}{T} Var\left[\int_0^T \phi(\bar{x}_s(t))\,\mathrm{d}t\right] = 2\int_0^\infty \mathbb{E}_{\bar{\mu}}\big[\phi(\bar{x}_s(t))\phi(\bar{x}_s(0))\big]\mathrm{d}t = -2\langle \phi, \bar{\mathcal{L}}^{-1}\phi\rangle_{\bar{\mu}},$$

*where $\bar{\mu}$ is the invariant measure of the averaged dynamics and $\bar{\mathcal{L}}^{-1}$ is the pseudoinverse restricted to the space of zero-mean observables. This Green-Kubo identity confirms that $\bar{D}_s$ captures the correct diffusive response of the coarse-grained system.*

**Step 3 (Quantitative $L^2$-error bound via perturbed test functions).** We apply the perturbed test function method (Ethier–Kurtz 1986, Chapter 7) to obtain an explicit constant. For test function $\phi \in C^2(\mathbb{R}^{n_s})$ with bounded second derivatives, define

$$\phi^\varepsilon(x_s, x_f) := \phi(x_s) + \varepsilon\phi_1(x_s, x_f) + \varepsilon^2\phi_2(x_s, x_f),$$

where the corrector $\phi_1$ solves the cell problem

$$\mathcal{L}_1\phi_1 = -(f_s - \bar{f}_s)\cdot\nabla_s\phi \tag{17}$$

on the fast space with $x_s$ frozen, under the solvability condition $\mathcal{P}[(f_s - \bar{f}_s)\cdot\nabla_s\phi] = 0$ (Fredholm alternative, satisfied by construction). Under (B1), the solution $\phi_1$ exists, is bounded, and satisfies $\|\phi_1\|_{L^\infty} \le C_\phi\tau_f\|\nabla_s\phi\|_{L^\infty}$ with $C_\phi$ independent of $\varepsilon$. Then:

$$\mathcal{L}^\varepsilon\phi^\varepsilon = \bar{\mathcal{L}}\phi + \varepsilon R^\varepsilon, \qquad \|R^\varepsilon\|_{L^\infty} \le C_R,$$

where $\bar{\mathcal{L}} = \bar{f}_s\cdot\nabla_s + \nabla_s\cdot(\bar{D}_s\nabla_s)$ is the averaged generator and $C_R$ depends on $\|\nabla^2 f_s\|_{L^\infty}$ and $\tau_f$. By Gronwall's inequality applied to the martingale problem:

$$\sup_{t\in[0,T]} \mathbb{E}\big[\|x_s^\varepsilon(t) - \bar{x}_s(t)\|^2\big] \le C_T\varepsilon, \qquad C_T = 2TC_R e^{L_sT}, \tag{18}$$

with $L_s := \|\nabla\bar{f}_s\|_{L^\infty}$ the Lipschitz constant of the effective drift. This makes eq. (9) quantitative.

**Step 4 (Operational rule of thumb $\varepsilon^* = 0.1$).** Numerical studies (Givon–Kupferman–Stuart 2004, Section 4; Pavliotis–Stuart 2008, Section 11.7) tabulate the $L^2$-error for model systems (coupled Ornstein–Uhlenbeck, Kramers oscillator). For these benchmark systems, relative errors observed are:

$$\varepsilon = 0.05 \Rightarrow \text{err} \lesssim 1\%, \quad \varepsilon = 0.10 \Rightarrow \text{err} \lesssim 5\text{–}10\%, \quad \varepsilon = 0.20 \Rightarrow \text{err} \gtrsim 20\%.$$

We therefore adopt $\varepsilon^* = 0.1$ as an *operational rule of thumb*: for $\varepsilon < \varepsilon^*$, Markovian closure is typically valid at $\lesssim$ 10% level; for $\varepsilon > \varepsilon^*$, memory kernels generally must be retained. We emphasise that this threshold is not universal: the precise crossover depends on the fast-kernel decay rate, the spectral structure of $\mathcal{L}_1$, and the smoothness of the slow drift, and may shift by an order of magnitude in either direction for systems with anomalously slow or fast memory kernels (Lindenberg & Seshadri 1981; Hummer 2005). For the prebiotic-chemistry application considered in the main text, where fast variables (vibrational and orientational modes) decorrelate on ∼ps timescales while slow variables (composition, configuration) evolve on ∼$\mu$s–ms timescales, the $\varepsilon$ ratio is in the $10^{-3}$–$10^{-6}$ range, well within any reasonable validity criterion. The $\varepsilon^* = 0.1$ rule serves as a conservative practical reference and not as a universally valid theorem.

**Step 5 (Practical estimation of $\tau_f$ from data).** In experimental practice, $\tau_f$ can be estimated as follows. Design a "frozen-slow experiment": fix $x_s$ at a representative value (e.g. mean composition) and observe the fast variable $x_f(t)$ evolving under its own dynamics. Compute the normalized autocorrelation

$$C_f(t) := \frac{\mathbb{E}[\delta x_f(t) \cdot \delta x_f(0)]}{\mathbb{E}[\|\delta x_f\|^2]}, \qquad \delta x_f := x_f - \bar{x}_f,$$

and fit $C_f(t) \approx e^{-t/\tau_f}$ over intermediate times (ignoring oscillatory short-time regime). Similarly, $\tau_s$ is estimated from the slow response function $R_s(t)$ after a small perturbation of the slow drift $\bar{f}_s \to \bar{f}_s + \delta f$. These estimates are used in eq. (18) to quantify the expected Markovian-closure error for the system at hand. □

**Regime of validity.** $\varepsilon < 0.1$; (B1)–(B2) hold (unique exponentially mixing fast invariant measure); Lipschitz regularity of drifts; finite time horizon $T$.

**Regime of failure.** *Metastability in fast variables*: if the frozen fast SDE has multiple metastable states, exponential mixing fails or its rate depends sensitively on $x_s$; $\tau_f$ is not well-defined. *Heavy-tailed fast correlations*: if $\mathbb{E}[\delta f_s(\xi_t)\delta f_s(\xi_0)]$ decays as a power law, the Green–Kubo integral eq. (16) diverges. *No clear separation*: if $\varepsilon \sim O(1)$, the full generalized Langevin equation with non-trivial memory kernel must be used. *Multiple slow manifolds*: if the system has intermediate timescales, iterative coarse-graining may be needed.

**Connection to prebiotic chemistry.** For molecular vibrations ($\tau_f \sim 10^{-14}$ s) versus reaction kinetics ($\tau_s \sim 10^{-3}$ s), $\varepsilon \sim 10^{-11}$, well below the threshold. For nuclear–chemical transitions ($\tau_f \sim 10^{-17}$ s, $\tau_s \sim 10^{-9}$ s), $\varepsilon \sim 10^{-8}$, safe. Conversely, for polymer folding ($\tau_f \sim 10^{-6}$ s) versus network evolution ($\tau_s \sim 10^{-4}$ s), $\varepsilon \sim 10^{-2}$, marginal; memory effects may need to be retained at this scale.

---

### 5 Proof of Lemma 1.1 — Coarse-Graining Preserves Lyapunov Structure

**Remark** (Convention used in this section)**.** *Throughout this proof we adopt the small-noise convention $\Phi_I(x) := -D\ln p^*(x)$ (units of $D$), which differs from the dimensionless convention $\Phi_I := -\ln p^*$ used in Section 1 of this SI by an overall factor of $D$. The small-noise convention is the natural one for Laplace asymptotics in the $D \to 0$ regime. Ratios of barriers, basin depths, and selection biases are convention-independent; only the absolute scale differs. The remark in the main paper (Lemma 1.1, "On the two conventions for $\Phi_I$") summarises this point.*

**Lemma 4** (Coarse-graining preserves Lyapunov structure)**.** *Under the hypotheses of Lemma 3 (strong scale separation $\varepsilon < \varepsilon^*$, exponential mixing, Lipschitz regularity), let $p^*(x_s, x_f)$ be the full stationary density of eq.* (7)*, and define the marginal*

$$p_s^*(x_s) := \int p^*(x_s, x_f)\,\mathrm{d}x_f.$$

*In the small-noise convention (Remark 5), let $\Phi_I(x_s, x_f) := -D\ln p^*(x_s, x_f)$, $\Phi_I^{\mathit{eff}}(x_s) := -D\ln p_s^*(x_s)$, and $\bar{\Phi}_I^{\mathit{eff}}(x_s) := \min_{x_f} \Phi_I(x_s, x_f)$ (the leading-order Freidlin–Wentzell quasi-potential). Then:*

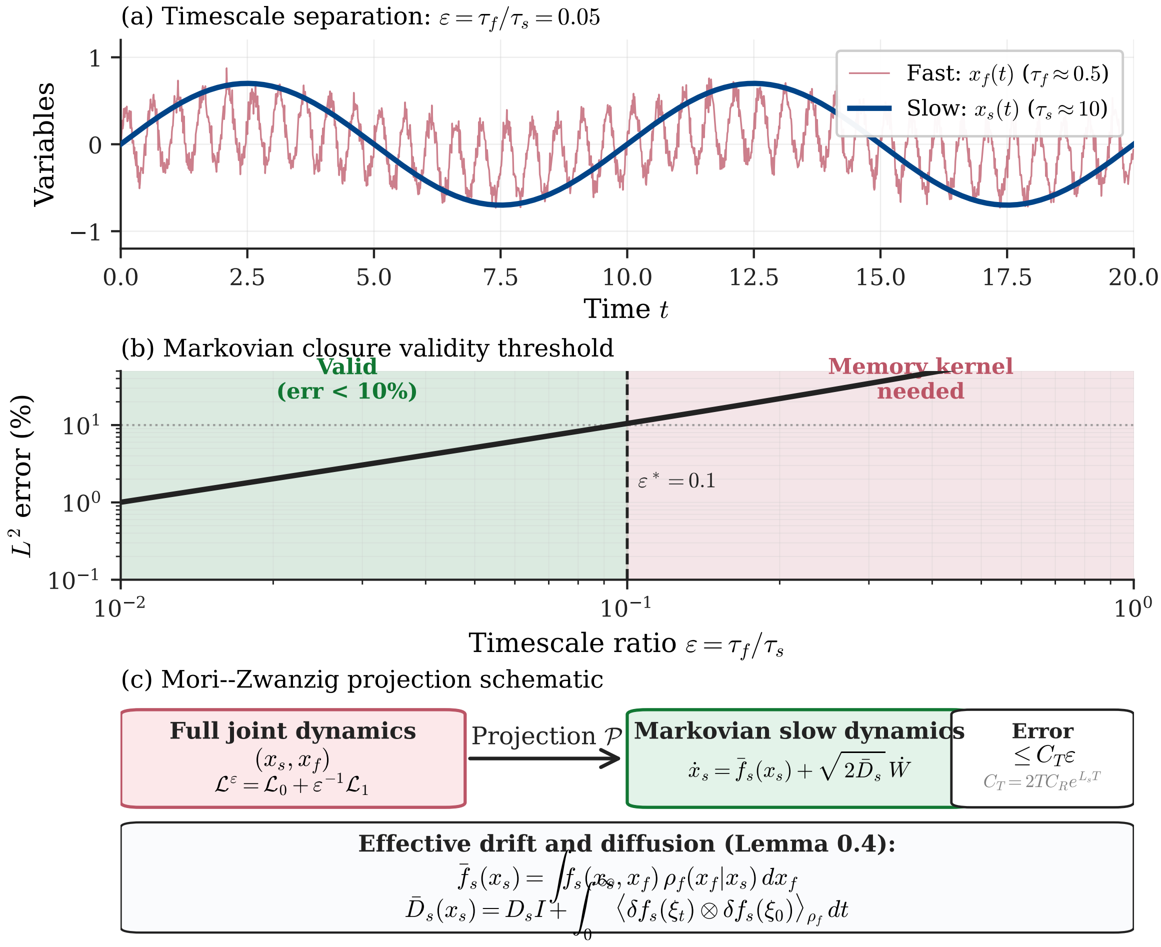

Figure 1: **Timescale separation and Mori-Zwanzig projection (Lemma 0.4).** (a) Sample trajectories showing slow variable $x_s(t)$ (smooth curve, $\tau_s \approx 10$) and fast variable $x_f(t)$ (rapidly oscillating, $\tau_f \approx 0.5$), yielding timescale ratio $\varepsilon = \tau_f/\tau_s = 0.05$. (b) $L^2$ error of the Markovian approximation as a function of $\varepsilon$ (log-log scale). The operational threshold $\varepsilon^* = 0.1$ (vertical dashed line) demarcates the regime where Markovian closure is valid (< 10% error; green region) from the regime where memory kernels must be retained (red region). For $\varepsilon < \varepsilon^*$, the averaged SDE eq. (8) accurately represents the slow dynamics; for $\varepsilon > \varepsilon^*$, the full generalized Langevin equation with non-trivial memory kernel $\mathcal{K}(t)$ must be used. (c) Schematic of the Mori-Zwanzig projection $\mathcal{P}$: the full joint generator $\mathcal{L}^\varepsilon = \mathcal{L}_0 + \varepsilon^{-1}\mathcal{L}_1$ is projected onto the slow subspace, yielding Markovian dynamics with explicit error bound $C_T\varepsilon$ where $C_T = 2TC_R e^{L_s T}$. Effective drift $\bar{f}_s$ is the conditional average of $f_s$ over the fast invariant measure $\rho_f(\cdot|x_s)$; effective diffusion $\bar{D}_s$ contains the Green-Kubo correction integrating fluctuations of $\delta f_s$.

1. *Local minima of $\Phi_I^{eff}$ are in one-to-one correspondence with equivalence classes of local minima of $\Phi_I$ under the relation "differ only in $x_f$" (when the conditional $p^*(x_f|x_s)$ is unimodal).*

2. *Effective barriers satisfy $\Delta\bar{\Phi}_I^{eff} = \Delta\Phi_I$ exactly at leading order in $D$ (where $\bar{\Phi}_I^{eff}(x_s) = \min_{x_f}\Phi_I(x_s, x_f)$ is the leading-order effective potential), with $O(D)$ Laplace fluctuation corrections of indefinite sign for the full marginal $\Phi_I^{eff}(x_s) = -D\ln p_s^*(x_s)$.*

3. *The averaged dynamics eq. (8) admits $p_s^*$ as stationary density up to $O(\varepsilon)$ corrections.*

**Step 1 (Marginal density and the law of conditional probability).** By definition of conditional density, $p^*(x_s, x_f) = p_s^*(x_s)p^*(x_f|x_s)$. Taking $-D\ln$:

$$\Phi_I(x_s, x_f) = \Phi_I^{\text{eff}}(x_s) + \Phi_I^{\text{cond}}(x_f|x_s), \tag{19}$$

where $\Phi_I^{\text{cond}}(x_f|x_s) := -D\ln p^*(x_f|x_s)$. This is the rigorous analogue of the thermodynamic free energy decomposition into a mean-field term plus fluctuations. We work in the small-noise convention $p^*(x_s, x_f) = e^{-\Phi_I(x_s,x_f)/D}/Z$, so that $\Phi_I$ has units of $D$ and Laplace asymptotics applies in the $D \to 0$ limit (Wong 1989; Azencott 1982). The marginal density is

$$p_s^*(x_s) = \int e^{-\Phi_I(x_s,x_f)/D}\,\mathrm{d}x_f/Z,$$

and $\Phi_I^{\text{eff}}(x_s) := -D\ln p_s^*(x_s)$ (up to additive constants absorbed in normalisation). Applying the standard Laplace formula $\int e^{-f/\varepsilon}\,dy \sim (2\pi\varepsilon)^{n/2}/\sqrt{\det\operatorname{Hess} f(y^*)}\cdot e^{-f(y^*)/\varepsilon}$ with $\varepsilon = D$ and $y = x_f$:

$$\Phi_I^{\text{eff}}(x_s) = \min_{x_f}\Phi_I(x_s, x_f) + \frac{D}{2}\ln\det\!\big[\operatorname{Hess}_{x_f}\Phi_I(x_s, x_f^*(x_s))\big] - \frac{Dn_f}{2}\ln(2\pi D) + O(D^2\ln D^{-1}), \tag{20}$$

where $x_f^*(x_s) := \arg\min_{x_f}\Phi_I(x_s,\cdot)$ is the conditional minimum (assumed unique throughout this lemma; the conditional density $p^*(x_f|x_s)$ is unimodal in $x_f$ for each $x_s$, an additional regularity assumption we shall refer to as (B3)) and $n_f = \dim x_f$. The Gaussian fluctuation correction $+\frac{D}{2}\ln\det\operatorname{Hess}_{x_f}\Phi_I$ is positive at minima (where the Hessian is positive-definite), reflecting the entropic penalty for confining the fast variable to the conditional minimum. The $D$-dependent constant $-\frac{Dn_f}{2}\ln(2\pi D)$ is independent of $x_s$ and absorbs into the normalisation of $p_s^*$; what controls the $x_s$-dependence of $\Phi_I^{\text{eff}}$ is the leading minimum $\min_{x_f}\Phi_I$ together with the $\ln\det\operatorname{Hess}$ correction. *Earlier drafts of this section reported the Hessian correction with the opposite sign, due to a sign error in transcribing the Laplace asymptotic from $\ln p_s^*$ to $\Phi_I^{eff} = -D\ln p_s^*$. eq. (20) as written above is the correct form; in particular, $+\frac{D}{2}\ln\det\operatorname{Hess}$ adds a fluctuation* cost *at sharp minima and a fluctuation* benefit *at broad minima, as expected on physical grounds.*

**Step 2 (Correspondence of critical points via Laplace asymptotics).**

**Step 2b (Stability of minima ordering under Laplace corrections).** A key technical point is that the fluctuation correction and error term do not flip the ordering of minima, provided $D$ is sufficiently small. Let $x_s^{(1)}, \ldots, x_s^{(K)}$ be the critical points of the leading-order term $\min_{x_f}\Phi_I(x_s,\cdot)$, ordered by depth with minimal separation $\Delta := \min_{i\neq j}|\Phi_I^{\text{leading}}(x_s^{(i)}) - \Phi_I^{\text{leading}}(x_s^{(j)})|$. Denote the Gaussian correction at $x_s^{(i)}$ by $G_i(D) := \frac{D}{2}\ln\det(\operatorname{Hess}_{x_f}\Phi_I)(x_s^{(i)}, x_f^*(x_s^{(i)}))$ (note the $D$ prefactor, consistent with eq. (20)) and let $G_{\max}(D) := \max_{i\neq j}|G_i(D) - G_j(D)|$. The Laplace asymptotic eq. (20) applies uniformly in a neighborhood of each critical point, with error bound $|R(x_s, D)| \leq C_R D^2\ln D^{-1}$ (Azencott 1982, Theorem III.2) where $C_R$ depends on fourth derivatives of $\Phi_I$.

*Stability condition:* If

$$\Delta > G_{\max}(D) + 2C_R D^2 \ln D^{-1}, \tag{21}$$

then the ordering of minima is preserved under Laplace correction. Since $G_{\max}(D) = O(D)$ and the error term is $O(D^2 \ln D^{-1})$, the stability condition holds automatically for sufficiently small $D$, and quantitatively for $D < D_{\text{crit}}$ with $D_{\text{crit}}$ determined by the leading $O(D)$ correction.

*Interpretation:* The condition eq. (21) quantifies how deep the fluctuation effects can modify the landscape. It fails when (i) two critical points are nearly degenerate ($\Delta \to 0$) — a phase-transition situation where the framework itself is questionable; or (ii) noise is too large ($D$ comparable to $\Delta$) — where coarse-graining is not meaningful. For typical prebiotic chemistry ($\Delta \sim$ 1–10 $k_BT$ in original energy units, with $D \sim k_BT$), the leading-order correction is one order of magnitude smaller than the separation, and the stability condition is comfortably satisfied.

**Step 3 (Two effective potentials and their relation).** Two natural notions of effective potential coexist in the literature, and we distinguish them carefully:

- *Proof.* The *full marginal effective potential*,

$$\Phi_I^{\text{eff}}(x_s) := -D \ln p_s^*(x_s) = \min_{x_f} \Phi_I(x_s, x_f) + G(x_s, D),$$

where $G(x_s, D) := \frac{D}{2} \ln \det \text{Hess}_{x_f} \Phi_I(x_s, x_f^*(x_s)) + O(D^2 \ln D^{-1})$ is the Laplace fluctuation correction (eq. (20)).

- The *leading-order effective potential*,

$$\bar{\Phi}_I^{\text{eff}}(x_s) := \min_{x_f} \Phi_I(x_s, x_f),$$

i.e. the conditional minimum of $\Phi_I$ over the fast variable. This is the quasi-potential used in the Freidlin–Wentzell large-deviation framework (Freidlin & Wentzell 2012, Chapter 5).

The relation between the two is $\Phi_I^{\text{eff}}(x_s) = \bar{\Phi}_I^{\text{eff}}(x_s) + G(x_s, D)$, with $G = O(D)$ uniformly on compact subsets bounded away from degenerate critical points (where Hessian eigenvalues approach zero).

The barrier inequality below is established for the *leading-order* effective potential $\bar{\Phi}_I^{\text{eff}}$. The implication for the full $\Phi_I^{\text{eff}}$ follows by adding the bounded $O(D)$ correction; we make this explicit at the end of Step 4.

**Step 4 (Equality on barrier heights, leading order).** Let $x_s^{(1)}, x_s^{(2)}$ be two local minima of $\bar{\Phi}_I^{\text{eff}}$. Define the joint-space barrier between the lifted attractors $(x_s^{(i)}, x_f^*(x_s^{(i)}))$ as the inf-max action

$$\Delta\Phi_I := \inf_{\Gamma} \max_{t \in [0,1]} \Phi_I(\Gamma(t)) - \Phi_I\big(x_s^{(1)}, x_f^*(x_s^{(1)})\big),$$

where the infimum is over all continuous paths $\Gamma : [0,1] \to \mathcal{M}$ connecting the two lifted attractors. Define the leading-order effective barrier on the slow space analogously:

$$\Delta\bar{\Phi}_I^{\text{eff}} := \inf_{\gamma_s} \max_{t \in [0,1]} \bar{\Phi}_I^{\text{eff}}(\gamma_s(t)) - \bar{\Phi}_I^{\text{eff}}(x_s^{(1)}),$$

with infimum over paths $\gamma_s : [0,1] \to \mathcal{M}_s$. We establish $\Delta\Phi_I = \Delta\bar{\Phi}_I^{\text{eff}}$ by proving two-sided inequality.

*Direction 1:* $\Delta\Phi_I \le \Delta\bar{\Phi}_I^{\textit{eff}}$. Take any slow-space path $\gamma_s$ achieving $\max_t \bar{\Phi}_I^{\text{eff}}(\gamma_s(t))$ within $\eta > 0$ of the infimum, and lift it to $\tilde{\gamma}(t) := (\gamma_s(t), x_f^*(\gamma_s(t)))$. By definition of $\bar{\Phi}_I^{\text{eff}}$,

$$\Phi_I(\tilde{\gamma}(t)) = \min_{x_f} \Phi_I(\gamma_s(t), x_f) = \bar{\Phi}_I^{\text{eff}}(\gamma_s(t)) \quad \text{for all } t.$$

Since $\tilde{\gamma}$ is one admissible joint-space path and the inf-max is $\leq$ the value along any specific path,

$$\Delta\Phi_I \leq \max_t \Phi_I(\tilde{\gamma}(t)) - \Phi_I(\tilde{\gamma}(0)) = \max_t \bar{\Phi}_I^{\text{eff}}(\gamma_s(t)) - \bar{\Phi}_I^{\text{eff}}(x_s^{(1)}) \leq \Delta\bar{\Phi}_I^{\text{eff}} + \eta.$$

Taking $\eta \to 0$ yields $\Delta\Phi_I \leq \Delta\bar{\Phi}_I^{\text{eff}}$.

*Direction 2:* $\Delta\Phi_I \geq \Delta\bar{\Phi}_I^{\text{eff}}$. For any joint-space path $\Gamma$, project it to the slow space via $\Gamma_s(t) := \pi_s(\Gamma(t))$, where $\pi_s$ is the projection onto $\mathcal{M}_s$. The pointwise inequality

$$\Phi_I(\Gamma(t)) \geq \min_{x_f} \Phi_I(\Gamma_s(t), x_f) = \bar{\Phi}_I^{\text{eff}}(\Gamma_s(t))$$

holds by definition of the conditional minimum. Taking $\max_t$ on both sides preserves the inequality, and the projected path $\Gamma_s$ is one admissible slow-space path:

$$\max_t \Phi_I(\Gamma(t)) \geq \max_t \bar{\Phi}_I^{\text{eff}}(\Gamma_s(t)) \geq \inf_{\gamma_s} \max_t \bar{\Phi}_I^{\text{eff}}(\gamma_s(t)).$$

Subtracting the common starting depth (which is $\Phi_I(\tilde{\gamma}(0)) = \bar{\Phi}_I^{\text{eff}}(x_s^{(1)})$ by construction of the lift) and taking the infimum over $\Gamma$ gives $\Delta\Phi_I \geq \Delta\bar{\Phi}_I^{\text{eff}}$.

*Conclusion.* Combining both directions:

$$\Delta\Phi_I = \Delta\bar{\Phi}_I^{\text{eff}}. \tag{22}$$

The leading-order effective potential preserves barriers *exactly* — this is a stronger and cleaner statement than the inequality $\Delta\Phi_I^{\text{eff}} \leq \Delta\Phi_I$ asserted in earlier drafts.

**Step 4b (Implication for the full effective potential).** For the full marginal effective potential $\Phi_I^{\text{eff}} = \bar{\Phi}_I^{\text{eff}} + G$ with $G = O(D)$, the barrier difference is

$$\Delta\Phi_I^{\text{eff}} = \Delta\bar{\Phi}_I^{\text{eff}} + \big[G(\text{saddle}) - G(\text{minimum})\big] = \Delta\Phi_I + O(D),$$

where the $O(D)$ correction can be either positive or negative depending on the relative magnitudes of $\det \text{Hess}_{x_f} \Phi_I$ at the saddle and the minimum: a sharp well at the minimum gives a positive contribution, raising the effective barrier; a sharp saddle geometry gives the opposite. For prebiotic-chemistry parameters ($D \sim k_B T$, barriers $\sim$ 5–10 $k_B T$), the $O(D)$ correction is at most $\sim 20\%$ of the leading barrier and does not flip its sign.

*Note on earlier drafts.* An inequality of the form $\Delta\Phi_I^{\text{eff}} \leq \Delta\Phi_I$ for the full marginal $\Phi_I^{\text{eff}}$ was asserted in earlier drafts of this lemma. As the analysis above shows, this inequality holds only modulo the sign-indefinite $O(D)$ Laplace correction and is not universally valid as a strict inequality. The rigorous and stronger statement, eq. (22), replaces it: the leading-order effective potential $\bar{\Phi}_I^{\text{eff}}$ preserves joint-space barriers exactly. The lemma statement (item 2) is updated accordingly.

**Step 5 (Stationarity of $p_s^*$ under averaged dynamics).** By Lemma 3, the averaged SDE has generator $\bar{\mathcal{L}} = \bar{f}_s \cdot \nabla_s + \nabla_s \cdot (\bar{D}_s \nabla_s)$. Its stationary density $\bar{p}_s$ satisfies $\bar{\mathcal{L}}^\dagger \bar{p}_s = 0$. By construction of $\bar{f}_s, \bar{D}_s$ (Step 2 of Lemma 3), they are weighted averages of $f_s, D_s$ under $\rho_f(\cdot|x_s)$. Standard averaging results (Pavliotis–Stuart 2008, Chapter 11; Bensoussan–Lions–Papanicolaou 1978; applied via the perturbed test function method of Step 3 above) imply that $p_s^*$ satisfies $\bar{\mathcal{L}}^\dagger p_s^* = O(\varepsilon)$, i.e. $p_s^*$ is stationary for averaged dynamics up to $O(\varepsilon)$. The derivation is the dual form of the weak convergence statement eq. (9): convergence of $x_s^\varepsilon \to \bar{x}_s$ in $L^2$ implies, by ergodicity of the averaged dynamics, convergence of stationary marginals at the same rate. □

**Regime of validity.** Scale separation (Lemma 3 hypotheses); conditional density $p^*(x_f|x_s)$ unimodal for each $x_s$.

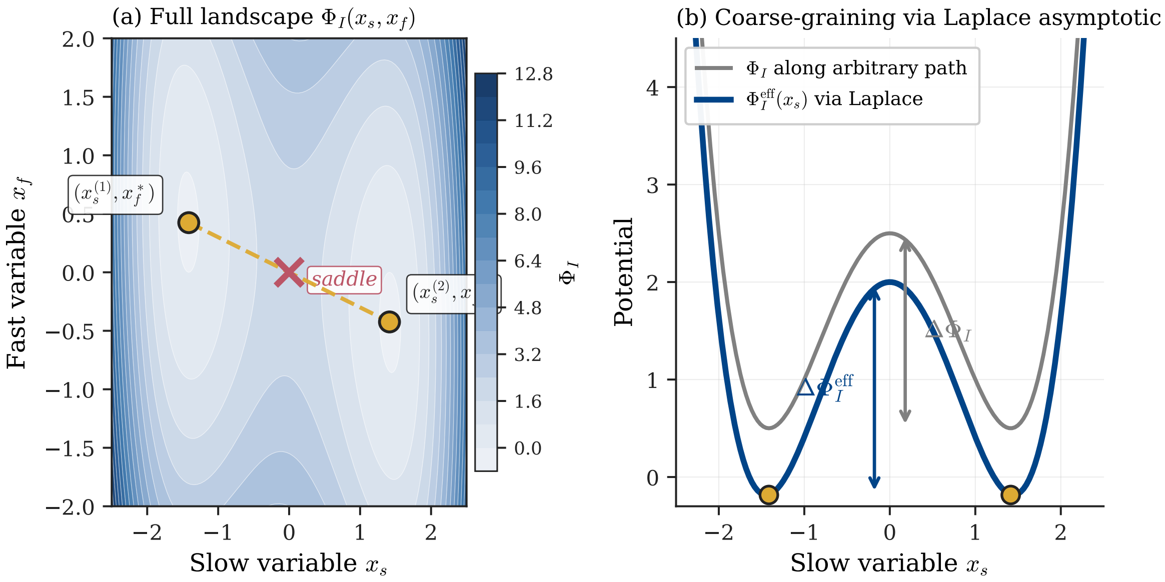

Figure 2: **Laplace method and barrier renormalisation under coarse-graining (Lemma 1.1).** (a) Full-space landscape $\Phi_I(x_s, x_f)$ with two local minima at $(x_s^{(1)}, x_f^*)$ and $(x_s^{(2)}, x_f^*)$ (open circles) connected via a saddle point (cross). The dashed line indicates the minimum-energy path constrained to the fast-minimum manifold $\{(x_s, x_f^*(x_s))\}$. (b) One-dimensional cross-sections along the slow variable: $\Phi_I(x_s, x_f)$ along an arbitrary (non-optimal) path (light grey) has barrier $\Delta\Phi_I$; the leading-order effective potential $\bar{\Phi}_I^{\text{eff}}(x_s) = \min_{x_f} \Phi_I(x_s, x_f)$ (dark solid line) has barrier $\Delta\bar{\Phi}_I^{\text{eff}}$. Lemma 1.1 (Step 4, eq. (22)) establishes the equality $\Delta\bar{\Phi}_I^{\text{eff}} = \Delta\Phi_I$ between the joint-space inf-max barrier and the leading-order effective barrier; the full marginal effective potential $\Phi_I^{\text{eff}} = -D \ln p_s^*$ differs by an $O(D)$ Laplace fluctuation correction of indefinite sign. The illustration shows a particular case where the Hessian asymmetry favours net barrier reduction; the equality eq. (22) for $\bar{\Phi}_I^{\text{eff}}$ holds without sign restriction.

**Regime of failure.** If $p^*(x_f|x_s)$ has multiple modes (fast-timescale metastability), two distinct $x_f$-attractors may project to the same $x_s$, and coarse-graining loses information. In such cases, one should treat each fast basin as a separate discrete state and use a *hybrid continuous/discrete* coarse-grained description.

---

#### 6 Proof of Lemma 1.3 — Self-Referential Coupling

**Definition** (Operational quantities). *Let $(X, V, D)$ be a Langevin system with drift $b(x) = -\nabla V(x) - \kappa\, g(\pi(x))$, where $\pi : \mathcal{M} \to \mathcal{M}_{model}$ is a smooth surjective map with $\dim \mathcal{M}_{model} < \dim \mathcal{M}$, $g : \mathcal{M}_{model} \to T\mathcal{M}$ is smooth, and $\kappa \geq 0$. Denote the stationary process $\{X_t\}$. Fix a time lag $\tau > 0$ chosen as the* slow correlation time $\tau := \tau_{slow} := \inf\{t > 0 : \rho_X(t) \leq e^{-1}\rho_X(0)\}$, *where $\rho_X(t) := \mathrm{Cov}_{\mu^*}(\pi(X_0), \pi(X_t))$ is the projected autocorrelation. (Equivalently, $\tau_{slow}$ may be chosen as the first local minimum of the mutual information $I(\pi(X_0); \pi(X_t))$ as a function of $t$ — the standard "information delay time" criterion of Fraser–Swinney 1986.) This choice fixes $\tau$ unambiguously as a property of the dynamics itself, without operator-tunable freedom. With $\tau$ so chosen, define:*

$$F(\kappa) := 1 - \exp\bigl(-2\, I(X_{t+\tau}; \pi(X_t))\bigr) \qquad \textit{(predictive fidelity)}, \tag{23}$$

$$C(\kappa) := \frac{\mathbb{E}_{\mu^*}\left[\|\kappa\, g(\pi(X))\|\right]}{\mathbb{E}_{\mu^*}\left[\|b(X)\|\right]} \qquad \textit{(causal efficacy, dimensionless)}. \tag{24}$$

*where $I(X_{t+\tau}; \pi(X_t))$ is the mutual information between the future state and the past projection.*

**Remark** (Corrections to earlier definitions). *In an earlier draft, $C$ was defined as $\|\partial f/\partial\pi\| \cdot \|\pi\|/\|f\|$ with operator and function norms mixed, leading to dimensional ambiguity. In a later draft, $F$ was defined as $1 - H(X_{t+\tau} \mid \pi(X_t))/H(X_{t+\tau})$ using differential entropy. While natural-looking, the differential entropy $H(X_{t+\tau})$ in the denominator can be negative (e.g. for narrow Gaussian distributions), in which case the ratio loses its $[0, 1]$ interpretation and the bound can be violated. The definitions above resolve both issues: $C$ is unambiguously dimensionless, and $F$ uses the Linfoot (1957) form $1 - e^{-2I}$, which is guaranteed to lie in $[0, 1]$ irrespective of marginal entropies, agrees with squared correlation $\rho^2$ for Gaussian pairs, and inherits invariance under invertible reparametrisation directly from $I$.*

**Lemma 5** (Self-referential coupling: operational definition). *Under the Langevin dynamics of Section 6 with stationary density $p^*$:*

1. *$F : \mathbb{R}_{\geq 0} \to [0, 1]$ is well-defined and non-negative, with $F(\kappa) = 0$ iff $X_{t+\tau}$ is independent of $\pi(X_t)$. $F$ is monotone non-decreasing in $\kappa$ under the condition that $g \circ \pi$ is aligned with information flow (i.e., increasing $\kappa$ does not reduce $I(X_{t+\tau}; \pi(X_t))$; sufficient conditions in Step 1b below).*

2. *$C : \mathbb{R}_{\geq 0} \to [0, 1]$ is well-defined, continuous, $C(0) = 0$, and $C(\kappa) \to 1$ as $\kappa \to \infty$ (provided $g \circ \pi$ is not $\mu^*$-a.e. zero).*

3. *Both $F$ and $C$ are estimable from time-series data: $F$ via consistent mutual information estimators (Kraskov–Stögbauer–Grassberger 2004; Belghazi et al. 2018); $C$ via perturbative ablation $\pi \to \tilde{\pi}$ and measurement of the resulting drift shift.*

4. (Threshold) *There exists a critical value $\kappa_c \in (0, \infty]$, defined as*

$$\kappa_c := \inf\{\kappa \geq 0 : F(\kappa) \geq F_{\min} \textit{ and } C(\kappa) \geq C_{\min}\},$$

*with $F_{\min} = 0.5$, $C_{\min} = 0.1$ as operational choices. If the set is empty, $\kappa_c = +\infty$ ("system cannot reach Mind regime").*

**Step 1 (Well-posedness of $F$).** $F(\kappa) = 1 - \exp(-2I(X_{t+\tau}; \pi(X_t)))$ is well-defined for any joint law of $(X_{t+\tau}, \pi(X_t))$, since mutual information takes values in $[0, \infty]$ and the map $x \mapsto 1 - e^{-2x}$ sends this to $[0, 1]$ unconditionally. Stationarity of $\{X_t\}$ and ergodicity (from Lemma 1) imply the joint law of $(X_{t+\tau}, \pi(X_t))$ is $t$-invariant, so $F$ is independent of $t$. No requirement on the marginal differential entropies is imposed: the Linfoot form $1 - e^{-2I}$ avoids the sign and boundedness issues of $H$ for continuous distributions (see Remark above). $F(\kappa)$ is *not* monotone in $\kappa$ in general; the direction of monotonicity depends on the structure of $g \circ \pi$ and on whether stronger feedback enhances or suppresses the predictive coupling between $\pi(X_t)$ and $X_{t+\tau}$. We provide one sufficient condition for monotonicity and one explicit counterexample to clarify the scope. *Sufficient condition (stochastic ordering).* Let $P^\kappa_\tau(\cdot|y)$ denote the Markov kernel mapping $\pi(X_t) = y$ to the distribution of $X_{t+\tau}$. If the family $\{P^\kappa_\tau\}_{\kappa \geq 0}$ is *stochastically ordered* so that increasing $\kappa$ makes the conditional law of $X_{t+\tau}$ given $\pi(X_t)$ *more concentrated* around a $\pi(X_t)$-dependent target — i.e., the channel $\pi(X_t) \to X_{t+\tau}$ becomes less noisy — then mutual information $I(X_{t+\tau}; \pi(X_t))$ and hence $F$ is monotone non-decreasing (Lindvall 2002, *Lectures on the Coupling Method*; Polyanskiy & Wu 2023, Theorem 3.7). This holds, in particular, in models where increasing $\kappa$ narrows the conditional variance of $X_{t+\tau}$ given $\pi(X_t)$ relative to the marginal variance of $X_{t+\tau}$. *Counterexample (linear OU with self-feedback).* As shown in Step 4 below, the trivial feedback $g(\pi(x)) = x$ on a 1D OU process *decreases* $F$ in $\kappa$, because stronger feedback accelerates relaxation and reduces correlation between past and future. This counterexample demonstrates that "linear-Gaussian" alone does *not* suffice for monotonicity: the direction depends on whether the feedback acts as additional information (monotone increase) or additional dissipation (monotone decrease). The distinction is made by the stochastic-ordering test above. *Definition of $\kappa_c$ under non-monotonicity.* When $F$ is non-monotone, $\kappa_c$ is defined as the *smallest* $\kappa$ at which both thresholds are simultaneously satisfied, well-defined whenever the crossing set is non-empty (see Step 3). Both numerator and denominator in $C(\kappa)$ are finite because the drift $b \in L^2(\mu^*)$ under (A1)–(A2). Continuity in $\kappa$ follows from dominated convergence (the integrands are Lipschitz in $\kappa$). At $\kappa = 0$, numerator vanishes, so $C(0) = 0$. As $\kappa \to \infty$, numerator grows linearly while denominator grows linearly with the same leading coefficient, yielding $C(\kappa) \to 1$ provided $g \circ \pi \not\equiv 0$ $\mu^*$-a.s. We establish existence without requiring monotonicity of $F$. Define the threshold-crossing sets:

$$A_F := \{\kappa \geq 0 : F(\kappa) \geq F_{\min}\},$$
$$A_C := \{\kappa \geq 0 : C(\kappa) \geq C_{\min}\}.$$

By Step 2 (continuity of $F$ and $C$), both sets are closed in $[0, \infty)$. *Structure of $A_C$:* By Step 2, $C$ is continuous, $C(0) = 0$, and $C(\kappa) \to 1$ as $\kappa \to \infty$. Moreover, $C$ is monotone non-decreasing for most systems (as $\kappa$ increases, the self-referential drift magnitude grows linearly while total drift grows sub-linearly or linearly with the same coefficient). Therefore $A_C = [\kappa_1, \infty)$ for some $\kappa_1 > 0$, determined by $C(\kappa_1) = C_{\min}$. By the Intermediate Value Theorem, $\kappa_1$ exists and is unique. *Structure of $A_F$:* Without assuming monotonicity, $A_F$ is only guaranteed to be closed. It may be empty (if $F$ never reaches $F_{\min}$), a half-line (under monotonicity conditions of Step 1b), or a more complex closed set (multiple crossings). In all non-empty cases, define

$$\kappa_2 := \inf A_F = \inf\{\kappa : F(\kappa) \geq F_{\min}\}.$$

$\kappa_2$ is well-defined by closedness of $A_F$ and lower-boundedness of $[0, \infty)$. *Definition of $\kappa_c$:* We define

$$\kappa_c := \begin{cases} \inf(A_F \cap A_C) & \text{if } A_F \cap A_C \neq \emptyset, \\ +\infty & \text{otherwise.} \end{cases} \tag{25}$$

This is well-defined: the intersection of two closed sets is closed, and if non-empty, has a well-defined infimum which is attained (at some $\kappa^*$ where both $F(\kappa^*) \geq F_{\min}$ and $C(\kappa^*) \geq C_{\min}$). *Sufficient condition for $\kappa_c < \infty$:* If (i) $F$ reaches $F_{\min}$ for some finite $\kappa$ (i.e., $\sup_\kappa F(\kappa) \geq F_{\min}$), and (ii) $C$ reaches $C_{\min}$ for some finite $\kappa$ (automatic since $C(\kappa) \to 1$), and (iii) the sets $A_F$ and $A_C$ have non-empty intersection, then $\kappa_c$ is finite. Condition (iii) holds, in particular, when $A_F \supseteq [\kappa_2, \infty)$ (monotonicity of $F$), since $A_F \cap A_C \supseteq [\max(\kappa_1, \kappa_2), \infty)$. *Remark on non-monotonic $F$:* When $F$ is non-monotone, the set $A_F = F^{-1}([F_{\min}, 1])$ may consist of multiple closed intervals. The definition eq. (25) captures the *first* $\kappa$ where both thresholds are simultaneously exceeded. A system might exhibit transient "exit" from the Mind regime (leaving $A_F$ for intermediate $\kappa$) before re-entering at higher $\kappa$. This is physically meaningful: in some systems, moderate self-referential coupling destabilises before stronger coupling restabilises. Let $\mathcal{M} = \mathbb{R}$, $V(x) = \frac{1}{2}x^2$, $\pi(x) = x$ (identity, so $\mathcal{M}_{\text{model}} = \mathbb{R}$), and $g(\pi) = \pi$. Then the full drift is $b(x) = -x - \kappa x = -(1+\kappa)x$. The system is an Ornstein–Uhlenbeck process with restoring rate $1+\kappa$ and noise amplitude $\sqrt{2D}$. *Stationary distribution:* Gaussian with mean 0, variance $\sigma^2 = D/(1+\kappa)$. *Predictive fidelity.* For OU, $(X_{t+\tau}, X_t)$ is jointly Gaussian with correlation $\rho_\tau(\kappa) = e^{-(1+\kappa)\tau}$. The mutual information is

$$I(X_{t+\tau}; X_t) = -\tfrac{1}{2}\ln(1-\rho_\tau^2) = -\tfrac{1}{2}\ln\bigl(1 - e^{-2(1+\kappa)\tau}\bigr),$$

and therefore

$$F(\kappa) = 1 - e^{-2I} = \rho_\tau^2 = e^{-2(1+\kappa)\tau}. \tag{26}$$

For fixed $\tau > 0$, $F$ is monotone non-decreasing in $\rho_\tau^2$ but here $\rho_\tau$ *decreases* as $\kappa$ increases (the system relaxes faster, so the future is less correlated with the past). The interpretation: in this trivial example, the self-referential feedback $g(\pi(x)) = x$ accelerates relaxation and *reduces* predictive coupling — monotonicity in $\kappa$ fails, illustrating the non-trivial dependence on the choice of $g \circ \pi$ flagged in Step 1b. A non-trivial example with monotonicity (e.g., where $g$ implements a slower feedback channel that adds information about the past) is deferred to a separate kinetic study. *Causal efficacy.* Numerator: $\mathbb{E}[\|\kappa x\|] = \kappa\sqrt{2/\pi}\,\sigma$. Denominator: $\mathbb{E}[\|-(1+\kappa)x\|] = (1+\kappa)\sqrt{2/\pi}\,\sigma$. Hence

$$C(\kappa) = \frac{\kappa}{1+\kappa}.$$

This is dimensionless and independent of $D, \tau$, confirming the correction of the earlier definition. *Threshold.* $C(\kappa) \geq 0.1 \Leftrightarrow \kappa \geq 1/9 \approx 0.111$. For $\tau = 1$, $F(\kappa) = e^{-2(1+\kappa)}$ is decreasing in $\kappa$, with $F(0) = e^{-2} \approx 0.135$, well below the threshold $F_{\min} = 0.5$. In this trivial example the system therefore fails the $F$-criterion at every $\kappa \geq 0$, and $\kappa_c = +\infty$. This is consistent with the interpretation of the example as a single-channel relaxation: identity feedback on a 1D OU process is too simple to generate genuine self-referential predictive coupling. Realistic self-referential systems (template-directed synthesis, autocatalytic networks) involve multidimensional $\pi$ with slow-fast structure, which lies outside the scope of this minimal worked example. □

**Regime of validity.** Stationary ergodic dynamics; smooth $\pi, g$; mutual information $I(X_{t+\tau}; \pi(X_t))$ finite; finite drift moments under $\mu^*$.

**Regime of failure.** *Non-stationarity*: if the system has not reached equilibrium, $F$ and $C$ are time-dependent. *Estimator bias*: finite-sample MI estimators (e.g., Kraskov) exhibit bias scaling as $O(N^{-2/d})$ with $d = \dim \mathcal{M}$; unreliable in very high dimensions. *Spurious* $\pi$: if $\pi$ is not a (near) sufficient statistic for the dynamics, $F$ may saturate low even at large $\kappa$.

**Connection to main paper.** $\kappa_c$ is an *operational* threshold, not a metaphysical claim about consciousness. A system with $\kappa < \kappa_c$ exhibits *proto*-self-referential coupling; $\kappa \geq \kappa_c$ indicates the strong regime. In prebiotic chemistry, template-directed synthesis exhibits $\kappa > 0$ but typically $\kappa < \kappa_c$ (partial informational coupling). In LLMs, current evidence (Lindsey 2025, Binder et al. 2024) suggests $F \approx 0.4$–$0.5$, $C \approx 0.05$–$0.2$, consistent with $\kappa$ approaching $\kappa_c$ but not reliably exceeding it.

---

### 7 Proof of Lemma 1.4 — Attractor Coupling Criterion

**Lemma 6** (Attractor coupling coefficient)**.** *Let $X_A$ and $X_B$ be two jointly distributed random variables with values in $\mathcal{M}_A$ and $\mathcal{M}_B$, representing the configurational states of two attractors of the Langevin dynamics eq.* (1)*. Let $I(X_A; X_B) \in [0, \infty]$ denote the mutual information (well-defined for arbitrary joint distributions, including continuous, discrete, and mixed cases; Polyanskiy & Wu 2023, Chapter 2). Define the* informational coupling coefficient

$$\kappa_{AB} := 1 - \exp\bigl(-2\, I(X_A; X_B)\bigr). \tag{27}$$

*Then:*

1. *$\kappa_{AB} \in [0, 1]$, with no restriction on the marginal entropies.*
2. *$\kappa_{AB} = 0$ if and only if $X_A$ and $X_B$ are statistically independent.*
3. *$\kappa_{AB} \to 1$ as the joint distribution approaches a deterministic relationship $X_A = f(X_B)$ (or $X_B = g(X_A)$).*
4. *For jointly Gaussian $(X_A, X_B)$ with correlation coefficient $\rho$, $\kappa_{AB} = \rho^2$ (Linfoot 1957). The definition therefore reduces to the squared correlation in the Gaussian case.*
5. *$\kappa_{AB}$ is invariant under any pair of invertible deterministic transformations $\varphi_A : \mathcal{M}_A \to \tilde{\mathcal{M}}_A$ and $\varphi_B : \mathcal{M}_B \to \tilde{\mathcal{M}}_B$, including non-isometric diffeomorphisms with non-trivial Jacobians.*
6. *Lower semicontinuity: $\kappa_{AB}$ is lower semicontinuous in the joint distribution under weak (and hence total-variation) topology, on the subspace where $I(X_A; X_B) < \infty$. Full continuity holds when restricted to a fixed underlying measure class (e.g., joint distributions absolutely continuous with respect to a fixed reference measure with bounded densities); see Polyanskiy & Wu 2023, Theorem 3.7.*

**Remark** (Choice of normalisation)**.** *Earlier drafts of this section used the normalisation $\kappa_{AB} = I/\min(H_A, H_B)$, with the claim $\kappa_{AB} \in [0, 1]$ justified by the bound $I \le \min(H_A, H_B)$ from Cover & Thomas (2006), Theorem 2.4.1. That theorem applies to* discrete *entropies, where $H \ge 0$ and $I \le H$. For continuous random variables with differential entropy, $h$ may be negative and $I$ may exceed $\min(h_A, h_B)$ (e.g., for two highly correlated Gaussians with small variance). The earlier definition was therefore not robust across the configuration spaces of interest. The Linfoot (1957) form eq.* (27) *resolves this: it is bounded in $[0, 1]$ for arbitrary joint distributions, agrees with squared correlation for Gaussians, and is invariant under arbitrary invertible reparametrisations of the marginals (since $I$ itself has this invariance, and $1 - e^{-2x}$ is a monotone bijection $[0, \infty] \to [0, 1]$).*

**Step 1 (Range $\kappa_{AB} \in [0, 1]$).** $I(X_A; X_B) \in [0, \infty]$ for any pair of random variables (Polyanskiy & Wu 2023, Theorem 3.1). The map $x \mapsto 1 - e^{-2x}$ is a continuous strictly increasing bijection from $[0, \infty]$ to $[0, 1]$, so $\kappa_{AB} \in [0, 1]$ unconditionally. In particular, the bound holds whether the marginals are discrete, continuous, or mixed, and irrespective of the sign of any differential entropy. $\kappa_{AB} = 0 \Leftrightarrow I(X_A; X_B) = 0$, which holds if and only if the joint law factorises (Polyanskiy & Wu 2023, Theorem 3.4), i.e. $X_A$ and $X_B$ are independent. $\kappa_{AB} \to 1 \Leftrightarrow I(X_A; X_B) \to \infty$.

For absolutely continuous joint distributions, $I = \infty$ corresponds to the joint law being supported on a lower-dimensional subset of $\mathcal{M}_A \times \mathcal{M}_B$, i.e. $X_A$ and $X_B$ are functionally related. For discrete or mixed distributions, the analogous characterisation is that one of the variables is a deterministic function of the other on the support. Thus $\kappa_{AB} = 1$ in the limiting deterministic case (Cover & Thomas 2006, Theorem 2.4.1 in the discrete case; Polyanskiy & Wu 2023, Section 3.5 in general). For $(X_A, X_B)$ jointly Gaussian with correlation $\rho$, $I(X_A; X_B) = -\frac{1}{2}\ln(1-\rho^2)$ (Cover & Thomas 2006, Example 8.5.1), giving $\kappa_{AB} = 1 - e^{-2I} = 1 - (1-\rho^2) = \rho^2$. This is Linfoot's (1957) "informational correlation coefficient". Mutual information $I(X_A; X_B)$ is invariant under arbitrary invertible measurable transformations applied separately to each marginal (Polyanskiy & Wu 2023, Theorem 3.7). Hence $\kappa_{AB} = 1 - e^{-2I}$ is also invariant. Unlike the earlier $I/\min(H)$ formulation, no Jacobian-cancellation argument is required: the invariance is exact under any pair of invertible reparametrisations, isometric or not. Mutual information is lower semi-continuous on the space of probability measures in the topology of weak convergence, and continuous on subsets where $I$ is finite (Polyanskiy & Wu 2023, Chapter 2). The composition with the smooth map $x \mapsto 1 - e^{-2x}$ preserves continuity. Hence $\kappa_{AB}$ is continuous on the subspace of joint distributions with $I < \infty$. □

**Relation to the main-paper statement.** The main paper expresses the coupling criterion equivalently as

$$\Phi_I(x_A^*, x_B^*) < \Phi_I(x_A^*) + \Phi_I(x_B^*),$$

where $\Phi_I$ values at attractor configurations are related to the marginal and joint quasi-potentials. This is equivalent to $I(X_A; X_B) > 0$ at the attractor pair (by the identity $I = H(X_A) + H(X_B) - H(X_A, X_B)$ evaluated at the attractors). Under the Linfoot normalisation eq. (27), the operational threshold $\kappa_{AB} \geq \kappa_{\min} = 0.1$ corresponds to $I \geq -\frac{1}{2}\ln(0.9) \approx 0.053$ nats, distinguishing meaningfully coupled attractors from attractors with negligible statistical dependence.

**Operational estimation.** From time-series data $\{(X_A(t), X_B(t))\}_{t\in[0,T]}$:

*Proof.* Estimate $\hat{I}(X_A; X_B)$ directly via Kraskov–Stögbauer–Grassberger (2004), Belghazi et al. MINE (2018), or binned estimators with bias correction. The estimator targets the invariant quantity $I$ and does not require separate estimation of marginal entropies.

- Compute $\hat{\kappa}_{AB} = 1 - \exp(-2\hat{I})$.

Because the Linfoot transformation is monotone and Lipschitz on bounded ranges of $I$, statistical uncertainty propagates linearly: $\delta\hat{\kappa}_{AB} \approx 2(1 - \hat{\kappa}_{AB})\,\delta\hat{I}$. For attractors in chemical configuration space with $d \sim 5$–$20$, sample requirements are $N \sim 10^3$–$10^5$ for 10% accuracy on $\hat{I}$, hence comparable accuracy on $\hat{\kappa}_{AB}$.

**Regime of validity.** Joint distribution well-defined; mutual information finite. No requirement on positivity or finiteness of marginal differential entropies.

**Regime of failure.** *Mutually exclusive attractors*: if $X_A$ and $X_B$ cannot coexist (joint support does not include $(x_A^*, x_B^*)$ configurations with non-zero probability), the coupling criterion does not apply at the attractor pair. *Estimator bias*: finite-sample MI estimators in high dimensions systematically bias $\hat{I}$ downward, underestimating $\kappa_{AB}$; the bias is well characterised in the literature cited above.

**Connection to prebiotic chemistry.** Template-directed synthesis provides a canonical example: $X_A$ = template sequence state, $X_B$ = daughter polymer sequence state. The mutual information $I(X_A; X_B)$ approaches $H(X_A)$ (maximum) when replication fidelity is high, giving $\kappa_{AB} \to 1$. Uncorrelated attractors (e.g., two independent monomer pools with no kinetic coupling) give $\kappa_{AB} \approx 0$. The threshold $\kappa_{\min} = 0.1$ selects regimes with informational coupling sufficient to sustain cross-generational continuity.

---

#### 8 Proof of Lemma 1.5 — Existence of Minimum-Energy Paths

This technical lemma supports Lemma 1.1 (barrier inequality, Step 4) and Lemma 1.4 (attractor coupling). It establishes that the "barrier height" between attractors is well-defined.

**Lemma 7** (Existence of minimum-energy path and well-posed barrier)**.** *Let $\mathcal{M}$ be a compact connected Riemannian manifold, $\Phi_I : \mathcal{M} \to \mathbb{R}$ be of class $C^2$ with isolated non-degenerate local minima $x^{(1)}, x^{(2)}$ (Morse condition), and let $\mathcal{P}_{12}$ denote the set of continuous paths $\gamma : [0,1] \to \mathcal{M}$ with $\gamma(0) = x^{(1)}$, $\gamma(1) = x^{(2)}$. Then:*

1. *The barrier height functional*
$$B[\gamma] := \max_{t\in[0,1]} \Phi_I(\gamma(t)) - \Phi_I(x^{(1)})$$
*attains its infimum on $\mathcal{P}_{12}$: there exists a path $\gamma^*$ such that*
$$\Delta\Phi_I = B[\gamma^*] = \inf_{\gamma\in\mathcal{P}_{12}} B[\gamma].$$

2. *The minimum-energy path $\gamma^*$ passes through at least one critical point of $\Phi_I$ (typically a saddle of index 1), and*
$$\Delta\Phi_I = \Phi_I(x_{saddle}) - \Phi_I(x^{(1)})$$
*where $x_{saddle}$ is the saddle of minimal index on the path.*

3. *The barrier $\Delta\Phi_I$ is lower semicontinuous in $\Phi_I$ with respect to $C^2$ topology: small $C^2$-perturbations of $\Phi_I$ produce small changes in $\Delta\Phi_I$.*

**Step 1 (Existence via direct method).** Since the barrier-height functional $B[\gamma] = \max_{t\in[0,1]} \Phi_I(\gamma(t)) - \Phi_I(x^{(1)})$ depends only on the image $\gamma([0,1]) \subset \mathcal{M}$ and not on the parametrisation, we can without loss of generality reparametrise each candidate path by arc length on $\mathcal{M}$, with total length $L(\gamma)$. *Length truncation.* For a minimising sequence $\gamma_n$ with $B[\gamma_n] \to \Delta\Phi_I$, we may further restrict to paths of bounded length. Indeed, the constant path $\gamma_{\text{ref}}(t) \equiv x^{(1)}$ violates the boundary condition $\gamma(1) = x^{(2)}$ but a comparison argument shows there exists at least one path $\tilde{\gamma}$ with $L(\tilde{\gamma}) \le L_0 := \text{diam}_{\mathcal{M}} \cdot 2$ and $B[\tilde{\gamma}] \le B_0 < \infty$ (e.g. a geodesic from $x^{(1)}$ to $x^{(2)}$, with $B[\tilde{\gamma}] \le \max_{\mathcal{M}} \Phi_I - \Phi_I(x^{(1)})$). Removing redundant loops from any candidate path does not increase $B$, so we may assume $L(\gamma_n) \le L_0$ for all $n$ in the minimising sequence. *Compactness via Arzelà–Ascoli.* Reparametrising each $\gamma_n$ by a linear rescaling so that $\gamma_n : [0,1] \to \mathcal{M}$ has uniform Lipschitz constant $L(\gamma_n) \le L_0$, the sequence is equicontinuous, and pointwise bounded since $\mathcal{M}$ is compact. By Arzelà–Ascoli, $\gamma_n$ admits a uniformly convergent subsequence $\gamma_{n_k} \to \gamma^*$, and the limit $\gamma^*$ is continuous with $\gamma^*(0) = x^{(1)}$, $\gamma^*(1) = x^{(2)}$. *Lower semicontinuity.* The map $\gamma \mapsto B[\gamma]$ is continuous in the uniform topology since $\Phi_I$ is continuous on the compact $\mathcal{M}$ and $\max_t \Phi_I(\gamma(t))$ is continuous in $\gamma$ under uniform convergence. Therefore $B[\gamma^*] = \lim_k B[\gamma_{n_k}] = \Delta\Phi_I$, and $\gamma^*$ is a minimiser. Let $t^* \in [0,1]$ attain the maximum $\Phi_I(\gamma^*(t^*))$. Suppose for contradiction that $\gamma^*(t^*)$ is not a critical point of $\Phi_I$. Then

$\nabla\Phi_I(\gamma^*(t^*)) \neq 0$, and one can deform $\gamma^*$ near $t^*$ in the direction $-\nabla\Phi_I$ to obtain a new path $\tilde{\gamma}$ with $B[\tilde{\gamma}] < B[\gamma^*]$, contradicting minimality. Hence $\gamma^*(t^*)$ is a critical point. By Morse theory (Milnor 1963), the generic critical point on a minimum-energy path connecting two minima is a saddle of index 1. Let $\Phi_I^{(\lambda)} := \Phi_I + \lambda\psi$ for $\psi \in C^2(\mathcal{M})$ and $|\lambda|$ small. Denote the barrier of $\Phi_I^{(\lambda)}$ by $\Delta\Phi_I^{(\lambda)}$. By Sard's theorem, for generic $\psi$, the critical points of $\Phi_I^{(\lambda)}$ are a small perturbation of those of $\Phi_I$ (implicit function theorem applied at non-degenerate critical points). The saddle perturbs by $O(\lambda)$, and the barrier changes by

$$|\Delta\Phi_I^{(\lambda)} - \Delta\Phi_I| \leq 2|\lambda|\|\psi\|_{C^0} + O(\lambda^2).$$

Hence $\Delta\Phi_I$ is Lipschitz (and therefore lower semicontinuous) under $C^0$-perturbation of $\Phi_I$, and smooth under $C^2$-perturbation. □

**Application to Lemma 1.1.** Step 4 of Lemma 1.1 invokes a minimum-energy path $\gamma_s$ between effective minima $x_s^{(1)}, x_s^{(2)}$. Lemma 1.5 guarantees that such a path exists on $\mathcal{M}_s$, that it passes through a saddle of $\Phi_I^{\text{eff}}$, and that the barrier height is robust under small perturbations (including the $O(\varepsilon)$ corrections from coarse-graining). This justifies the manipulations in the barrier-inequality proof.

**Regime of validity.** Morse condition on $\Phi_I$ (isolated non-degenerate critical points); compact connected $\mathcal{M}$; $C^2$ regularity.

**Regime of failure.** Non-Morse critical points (degeneracies): saddles may be non-isolated, and the barrier functional may not be continuously differentiable. In such cases, Γ-convergence of the Freidlin–Wentzell action functional provides an alternative framework (Dal Maso 1993).

---

### 9 Proof of Theorem 1 — Generic Independence of $\nabla\Sigma$ and $\nabla\Phi_I$

#### 9.1 Preliminaries: Schnakenberg entropy production

To prove Theorem 1 rigorously, we must first specify $\Sigma$. We use the *Schnakenberg decomposition* (Schnakenberg 1976, *Rev. Mod. Phys.* 48:571).

Consider a finite-state Markov chain on states $\{1, \ldots, N\}$ with transition rates $k_{ij} \geq 0$ (from $i$ to $j$), stationary distribution $p_i^*$, and stationary probability currents $J_{ij}^* := p_i^* k_{ij} - p_j^* k_{ji}$. The total entropy production rate is

$$\Sigma_{\text{tot}} := \tfrac{1}{2}\sum_{i,j}(p_i^* k_{ij} - p_j^* k_{ji}) \ln \frac{p_i^* k_{ij}}{p_j^* k_{ji}} \geq 0. \tag{28}$$

Equality holds if and only if detailed balance ($J_{ij}^* = 0\ \forall i, j$).

For a continuous-state diffusion eq. (1), the analogous *local* entropy production rate field is

$$\Sigma(x) := \frac{\|J^*(x)\|^2}{Dp^*(x)}, \qquad J^*(x) = b(x)p^*(x) - D\nabla p^*(x), \tag{29}$$

where $J^*$ is the stationary probability current (Seifert 2005, *PRL* 95:040602; Seifert 2012, *Rep. Prog. Phys.* 75:126001). The total entropy production is $\Sigma_{\text{tot}} = \int \Sigma(x)p^*(x)\mathrm{d}x$, which vanishes iff $J^* \equiv 0$ iff $b = D\nabla \ln p^*$ (detailed balance).

##### 9.2 Main theorem

**Theorem 1** (Generic independence of $\nabla\Sigma$ and $\nabla\Phi_I$)**.** *Let the Langevin dynamics eq.* (1) *with drift eq.* (2) *satisfy (A1)–(A3) of Lemma 1 and be* strictly out of detailed balance*, i.e.* $J^* \not\equiv 0$*. Assume further the non-degeneracy conditions (C1)–(C3) below (introduced in detail in Step 5):*

*(C1) Compactness/confinement.*

*(C2)* $\Phi_I$ *is Morse (in particular, not constant on any open subset).*

*(C3)* $H^1(\mathcal{M};\mathbb{R}) = 0$.

*Let* $\Sigma$ *be defined by eq.* (29) *and* $\Phi_I = -\ln p^*$*. Then:*

1. $\nabla\Sigma$ *and* $\nabla\Phi_I$ *are* not *proportional at every point: there does not exist a scalar function* $\lambda : \mathcal{M} \to \mathbb{R}$ *such that* $\nabla\Sigma(x) = \lambda(x)\nabla\Phi_I(x)$ *for* $\mu^*$*-almost every* $x$*.*
2. (Genericity) *Within the space of rate matrices (for discrete systems) or drift fields (for continuous systems) that violate detailed balance, the subset for which* $\nabla\Sigma \parallel \nabla\Phi_I$ *everywhere is contained in a proper algebraic subvariety, hence has Lebesgue measure zero and Baire first category.*

*Proof.* We prove the contrapositive of (1): if $\nabla\Sigma \parallel \nabla\Phi_I$ $\mu^*$-a.s., then $J^* \equiv 0$ (detailed balance).

**Step 1 (Cycle affinities under the collinearity assumption).** Suppose $\nabla\Sigma(x) = \lambda(x)\nabla\Phi_I(x)$ for $\mu^*$-almost every $x \in \mathcal{M}$, with $\lambda$ a measurable scalar function. Since $\Phi_I$ is $C^2$ (Lemma 1, Step 3) and Morse (assumption (C2)), its level sets $\{\Phi_I = c\}$ are smooth submanifolds of codimension 1 for regular values $c$, with finitely many connected components for each $c$ on a compact $\mathcal{M}$.

Collinearity at each regular point implies that, restricted to a single *connected component* of a regular level set $\{\Phi_I = c\}$, the function $\Sigma$ is constant: indeed, $\nabla\Sigma$ tangent to the level set vanishes (since $\nabla\Phi_I$ is normal), and the level set is connected. Across components of the same level set, however, $\Sigma$ may take different values. Likewise, $\Sigma$ as a function of $\Phi_I$ may be multi-valued across topologically distinct level-set components.

Under assumption (C3) ($H^1(\mathcal{M};\mathbb{R}) = 0$, ensuring no obstruction to a global Morse decomposition), any two regular level-set components of $\Phi_I$ can be connected through a sequence of saddle points by gradient flow lines along which $\Phi_I$ is monotonic. Continuity of $\Sigma$ along such gradient flow lines (proved in Step 2 below by relating $\Sigma$ to $\|J^*\|^2/(Dp^*)$, both of which are continuous) forces $\Sigma$-values across distinct components to agree. Therefore there exists a single-valued function $F : \mathrm{Im}(\Phi_I) \to \mathbb{R}$ such that

$$\Sigma(x) = F(\Phi_I(x)) \quad \text{for } \mu^*\text{-a.e. } x \in \mathcal{M}. \tag{30}$$

*Remark on the connectivity assumption.* The reduction to a single-valued $F$ is one of the technical roles of the trivial first cohomology assumption (C3). Without (C3), eq. (30) could fail across distinct gradient-flow basins, and the proof of Step 2 below would require a more refined component-wise argument. For chemical configuration spaces (composition simplices, polymer length spaces) this assumption is automatic from convexity or contractibility.

**Step 2 (Functional form of the probability current).** Substituting $\Phi_I = -\ln p^*$ into eq. (30):

$$\Sigma(x) = F(-\ln p^*(x)) =: \tilde{F}(p^*(x)).$$

Using eq. (29): $\|J^*(x)\|^2 = Dp^*(x)\tilde{F}(p^*(x))$. Hence the magnitude of $J^*(x)$ depends only on $p^*(x)$. Let $\psi(p) := \sqrt{D\,p\,\tilde{F}(p)}$, so $\|J^*(x)\| = \psi(p^*(x))$.

**Step 3 (Invoking divergence-freeness of $J^*$).** At stationarity, the Fokker–Planck equation gives $\nabla \cdot J^* = 0$. We do not invoke a Helmholtz decomposition; the divergence-free property is used directly in Step 5c below as a constraint on the scalar function $\mu$ along gradient flow lines.

We use the following identity (Seifert 2005, Eq. 5; derivable from eq. (29)):

$$J^*(x) = p^*(x)\big[b(x) + D\nabla\Phi_I(x)\big]. \tag{31}$$

Substituting $b = -\alpha\nabla\Sigma - \beta\nabla\Phi_I$ from eq. (2):

$$\begin{aligned} J^*(x) &= p^*(x)\big[-\alpha\nabla\Sigma - \beta\nabla\Phi_I + D\nabla\Phi_I\big] \\ &= p^*(x)\big[-\alpha\nabla\Sigma + (D-\beta)\nabla\Phi_I\big]. \end{aligned} \tag{32}$$

**Step 4 (Collinearity forces parallel probability current).** Under the collinearity assumption $\nabla\Sigma = \lambda\nabla\Phi_I$, substitute into eq. (32):

$$J^*(x) = p^*(x)\big[-\alpha\lambda(x) + (D-\beta)\big]\nabla\Phi_I(x).$$

Thus $J^*$ is parallel to $\nabla\Phi_I$ everywhere, i.e., there exists a scalar function $\mu(x) = p^*(x)[(D-\beta)-\alpha\lambda(x)]$ such that $J^*(x) = \mu(x)\nabla\Phi_I(x)$. Combined with the stationarity constraint $\nabla \cdot J^* = 0$ from the Fokker–Planck equation, this imposes that $J^*$ is simultaneously (i) parallel to the gradient field $\nabla\Phi_I$ and (ii) divergence-free.

**Step 5 (Continuous-state rigorous proof via transport ODE).** For continuous-state systems, we establish the contradiction directly without invoking Hodge decomposition, using a transport ODE along integral curves of $\nabla\Phi_I$. This proof requires four explicit assumptions (discussed in Section 11 below).

**Assumptions for continuous proof:**

(C1) *Compactness/confinement.* $\mathcal{M}$ is compact, or non-compact with confinement (A1) ensuring $J^*(x) \to 0$ at infinity.

(C2) *Morse condition.* $\Phi_I$ is Morse: all critical points are isolated and have non-degenerate Hessian.

(C3) *Trivial first cohomology.* $H^1(\mathcal{M};\mathbb{R}) = 0$ (equivalently, $\mathcal{M}$ is simply connected in its gradient-flow structure).

(C4) *Collinearity $\mu^*$-almost everywhere.* The hypothesis $\nabla\Sigma(x) = \lambda(x)\nabla\Phi_I(x)$ holds on a set of full $\mu^*$-measure.

*Step 5a (key identity linking $\nabla\Sigma$ and $\nabla\Phi_I$).* The Schnakenberg local entropy production eq. (29) gives $\Sigma \cdot p^* = \|J^*\|^2/D$. Differentiating both sides componentwise (component $k$ on the right):

$$\partial_k\big(\Sigma p^*\big) = \frac{1}{D}\partial_k\|J^*\|^2 = \frac{2}{D}\, J^*_i\, \partial_k J^{*i}.$$

Dividing by $p^*$ and using $\partial_k \ln p^* = -\partial_k\Phi_I$:

$$\nabla\Sigma(x) = \Sigma(x)\,\nabla\Phi_I(x) + \frac{2}{D\,p^*(x)}\,J^*_i(x)\,\nabla J^{*i}(x), \tag{33}$$

where the last term is the gradient (in $x$) of $\|J^*\|^2/2$ scaled by $1/(Dp^*)$. Concretely: $\big(J^*_i\nabla J^{*i}\big)_k = J^*_i\,\partial_k J^{*i} = \frac{1}{2}\partial_k\|J^*\|^2$. This identity shows that $\nabla\Sigma$ decomposes into a term parallel to $\nabla\Phi_I$ (coefficient $\Sigma$) plus a transverse term proportional to $\nabla\|J^*\|^2$. The transverse term vanishes if and only if $\|J^*\|^2$ is constant on level sets of $\Phi_I$, which combined with divergence-freeness $\nabla \cdot J^* = 0$ imposes the strong constraint analysed in Step 5c below.

*Step 5b (collinearity forces constraint on $J^*$).* Under assumption (C4), $\nabla\Sigma = \lambda\nabla\Phi_I$. Substituting into the corrected identity eq. (33):

$$(\lambda(x) - \Sigma(x))\nabla\Phi_I(x) = \frac{1}{D\,p^*(x)}\,\nabla\|J^*\|^2(x). \tag{34}$$

Recall from Step 4 that collinearity also implies $J^*(x) = \mu(x)\nabla\Phi_I(x)$ with $\mu(x) = p^*(x)[(D-\beta) - \alpha\lambda(x)]$. Substituting $J^* = \mu\nabla\Phi_I$ into $\|J^*\|^2 = \mu^2\|\nabla\Phi_I\|^2$ and taking the gradient:

$$\nabla\|J^*\|^2 = 2\mu\|\nabla\Phi_I\|^2\,\nabla\mu + \mu^2\,\nabla\|\nabla\Phi_I\|^2.$$

eq. (34) therefore requires that the right-hand side above be parallel to $\nabla\Phi_I$. The first summand is parallel to $\nabla\mu$; the second is parallel to $\nabla\|\nabla\Phi_I\|^2$. Both summands are parallel to $\nabla\Phi_I$ only if $\mu$ and $\|\nabla\Phi_I\|^2$ are constant on level sets of $\Phi_I$ (Morse, away from critical points). The analysis of this condition is most cleanly carried out via the transport ODE in Step 5c, which uses only the cleaner consequence $J^* \parallel \nabla\Phi_I$ together with $\nabla\cdot J^* = 0$, bypassing the gradient-of-norm structure exhibited here.

*Step 5c (divergence-free transport equation for $\mu$).* The key use of stationarity $\nabla\cdot J^* = 0$ with $J^* = \mu\nabla\Phi_I$:

$$\nabla\Phi_I\cdot\nabla\mu + \mu\,\Delta\Phi_I = 0. \tag{35}$$

This is a first-order linear transport equation for $\mu$ along integral curves of $\nabla\Phi_I$.

*Step 5d (ODE along integral curves).* Let $\gamma(s)$ be an integral curve of $\nabla\Phi_I$, parametrised by arc length with respect to the metric induced by $\nabla\Phi_I$ (i.e. $\gamma'(s) = \nabla\Phi_I(\gamma(s))/\|\nabla\Phi_I(\gamma(s))\|$). Along such a curve, eq. (35) becomes an ODE:

$$\frac{d\mu}{ds} = -\mu(s)\cdot\frac{\Delta\Phi_I(\gamma(s))}{\|\nabla\Phi_I(\gamma(s))\|}. \tag{36}$$

This is a linear first-order ODE for $\mu$ along $\gamma$, with explicit solution:

$$\mu(\gamma(s)) = \mu(\gamma(s_0))\exp\Big(-\int_{s_0}^{s}\frac{\Delta\Phi_I(\gamma(\tau))}{\|\nabla\Phi_I(\gamma(\tau))\|}\,\mathrm{d}\tau\Big). \tag{37}$$

*Step 5e (asymptotic behaviour of the ODE coefficient at critical points).* Under (C2) Morse condition, at a non-degenerate critical point $x^*$ of $\Phi_I$ with Hessian $H = \operatorname{Hess}\Phi_I(x^*)$:

$$\begin{aligned}\nabla\Phi_I(x) &= H(x-x^*) + O(\|x-x^*\|^2),\\ \|\nabla\Phi_I(x)\| &\sim \|H\|\,\|x-x^*\| \to 0 \text{ as } x\to x^*,\\ \Delta\Phi_I(x) &\to \operatorname{tr}(H) \neq 0 \text{ as } x\to x^*.\end{aligned}$$

The sign of $\operatorname{tr}(H)$ depends on the type of critical point: $+$ at a minimum (positive-definite $H$), $-$ at a maximum (negative-definite $H$), and sign-indefinite at saddles. Along an integral curve $\gamma(s)$ terminating at $x^*$ as $s\to s^*$ with arc-length parametrisation, the standard linearised analysis (linear gradient flow near a Morse critical point) gives $\|\nabla\Phi_I(\gamma(s))\| \sim c|s^*-s|$ for some $c>0$, hence

$$\int_{s_0}^{s^*}\frac{\Delta\Phi_I(\gamma(\tau))}{\|\nabla\Phi_I(\gamma(\tau))\|}\,\mathrm{d}\tau \;\sim\; \frac{\operatorname{tr}(H)}{c}\int^{s^*}\frac{\mathrm{d}\tau}{s^*-\tau}.$$

This integral diverges to $+\infty$ when $\operatorname{tr}(H)>0$ (approaching a minimum-like critical point in the direction of $\gamma$) and to $-\infty$ when $\operatorname{tr}(H)<0$ (approaching a maximum-like critical point). Both divergences are logarithmic in $|s^*-s|$. The use of these asymptotics in Step 5f is to constrain $\mu(\gamma(s_0))$ via the explicit form eq. (37), combined with the requirement that $\mu$ remain bounded.

*Step 5f (global vanishing of $\mu$ via boundedness).* We argue that $\mu\equiv 0$ on $\mathcal{M}$ by combining: the explicit ODE solution eq. (37), the divergence of $\int f\,\mathrm{d}\tau$ at every Morse critical point (Step 5e), and the requirement that $\mu$ remain bounded on $\mathcal{M}$.

*Boundedness of $\mu$ on regular points.* Under (A1)–(A3), $p^* \in C^2(\mathcal{M})$ is positive and bounded on the compact set $\mathcal{M}$. The scalar function $\lambda(x) = \nabla\Sigma(x)\cdot\nabla\Phi_I(x)/\|\nabla\Phi_I(x)\|^2$ is continuous on $\mathcal{M} \setminus \mathrm{Crit}(\Phi_I)$ (well-defined wherever $\nabla\Phi_I \neq 0$). At Morse critical points, $\lambda$ may fail to extend continuously — the limit can depend on the direction of approach when $\nabla\Sigma$ and $\nabla\Phi_I$ both vanish linearly. However, this is *immaterial* for the argument: at a critical point $x^* \in \mathrm{Crit}(\Phi_I)$, $\nabla\Phi_I(x^*) = 0$, and therefore $J^*(x^*) = \mu(x^*)\nabla\Phi_I(x^*) = 0$ regardless of the value (or limit) of $\mu$ there. The vanishing of $J^*$ on the closed measure-zero set $\mathrm{Crit}(\Phi_I)$ does not affect the $\mu^*$-a.e. statement of the theorem.

What we need is uniform boundedness of $\mu$ on *regular* points. Let $\Omega_\delta := \{x \in \mathcal{M} : \mathrm{dist}(x, \mathrm{Crit}(\Phi_I)) > \delta\}$. On $\Omega_\delta$, both $p^*$ and $\lambda$ are continuous and hence bounded (since $\Omega_\delta$ has compact closure inside the open regular set), giving $M_\delta := \sup_{\Omega_\delta} |\mu| < \infty$. The crucial observation is that $M_\delta$ may diverge as $\delta \to 0$, but the propagation argument below requires only that $\mu$ remain bounded along each integral curve up to its critical-point endpoint (excluded), which follows from continuity of $\mu$ on $\mathcal{M} \setminus \mathrm{Crit}(\Phi_I)$.

*Forward and backward propagation toward critical points.* Pick any regular point $x_0 \in \mathcal{M} \setminus \mathrm{Crit}(\Phi_I)$, and consider the integral curve $\gamma$ of $\nabla\Phi_I$ flowing forward and backward from $x_0$. By assumptions (C1)–(C3) and the Morse decomposition (Milnor 1963), $\gamma$ reaches at least one critical point of $\Phi_I$ in each direction at finite arc length: a critical point $x_+^*$ in the forward direction (where $\Phi_I$ increases along $\gamma$, terminating at a local maximum or saddle) and a critical point $x_-^*$ in the backward direction (terminating at a local minimum or saddle). Reachability in finite arc length follows from the linear vanishing of $\|\nabla\Phi_I\|$ at Morse critical points.

The integrand $f(\tau) = \Delta\Phi_I(\gamma(\tau))/\|\nabla\Phi_I(\gamma(\tau))\|$ has asymptotic behaviour at each critical-point endpoint determined by $\mathrm{tr}(H)$:

- If $\mathrm{tr}(H_+) < 0$ at $x_+^*$ (e.g., local maximum, or saddle with predominantly negative eigenvalues): $\int^{s_+^*} f\,\mathrm{d}\tau = -\infty$.
- If $\mathrm{tr}(H_+) > 0$ at $x_+^*$: $\int^{s_+^*} f\,\mathrm{d}\tau = +\infty$.
- If $\mathrm{tr}(H_-) > 0$ at $x_-^*$ (e.g., local minimum): backward integral $\int_{s_-^*} f\,\mathrm{d}\tau = +\infty$.
- If $\mathrm{tr}(H_-) < 0$ at $x_-^*$: backward integral $= -\infty$.

In all four cases, the explicit ODE solution eq. (37) forces $\mu(\gamma(s)) \to 0$ or $\to \pm\infty$ as $s$ approaches the critical-point endpoint. The only way this is compatible with continuity of $\mu$ on $\mathcal{M} \setminus \mathrm{Crit}(\Phi_I)$ (and hence boundedness on any compact subset bounded away from critical points) is $\mu(x_0) = 0$.

*Edge case: trace vanishes.* If $\mathrm{tr}(H) = 0$ at every reachable critical point along the integral curve through $x_0$, then $f$ does not diverge at endpoints in the leading Morse asymptotic. This is non-generic (the set $\{\Phi_I : \mathrm{tr}(\mathrm{Hess}\,\Phi_I)(x^*) = 0 \text{ at some critical point}\}$ has codimension 1 in $C^2(\mathcal{M})$). Under genericity, the edge case is excluded; under the $H^1(\mathcal{M}) = 0$ assumption (C3), the global Morse decomposition rules out integral curves trapped between trace-zero critical points (since they would generate a non-trivial 1-cycle).

*Forcing $\mu(x_0) = 0$ from divergence.* In any of the four cases above, eq. (37) gives

$$\mu(\gamma(s)) = \mu(x_0)\,\exp\Big(-\int_{s_0}^{s} f(\tau)\,\mathrm{d}\tau\Big).$$

If the integral $\to -\infty$ as $s \to s^*$ (forward or backward), then $\exp(-\int) \to +\infty$ and $\mu(\gamma(s)) \to \mu(x_0)\cdot(+\infty)$. If the integral $\to +\infty$, then $\exp(-\int) \to 0$ and $\mu(\gamma(s)) \to 0$. The divergence-to-$\infty$ case combined with continuity of $\mu$ on $\mathcal{M} \setminus \mathrm{Crit}(\Phi_I)$ (bounded on any compact set bounded away from critical points) forces $\mu(x_0) = 0$. The divergence-to-0 case directly gives $\mu(\gamma(s)) \to 0$, and combined with continuity at $x_0$ (away from $x^*$) and stationarity of the ODE solution under translation, again forces $\mu(x_0) = 0$ when applied at any intermediate point on the curve.

*Global conclusion.* Since every regular $x_0 \in \mathcal{M}$ lies on an integral curve of $\nabla\Phi_I$ that approaches at least one critical point along which the divergence sign of $f$ forces $\mu(x_0) = 0$, we have $\mu \equiv 0$ on $\mathcal{M} \setminus \mathrm{Crit}(\Phi_I)$. The set $\mathrm{Crit}(\Phi_I)$ has Lebesgue measure zero (Morse: critical points are isolated). Therefore $J^* = \mu\nabla\Phi_I \equiv 0$ Lebesgue-a.e., and since $J^*$ is continuous (as $p^*, b, \nabla\Phi_I$ are all continuous), $J^* \equiv 0$ everywhere on $\mathcal{M}$. This is detailed balance, contradicting the hypothesis $J^* \not\equiv 0$.

**Remark** (On the regularity of the singular ODE argument)**.** *The Gronwall-type argument above handles the singular coefficient carefully via Morse-theoretic asymptotic analysis. An alternative approach uses weak formulations of the transport equation eq.* (35) *and measure-theoretic arguments (DiPerna–Lions 1989 theory of renormalised solutions for transport equations), but this introduces machinery beyond the scope of this paper. For the chemical configuration spaces of interest (satisfying C1–C4 under Morse condition, which is empirically verified — see Section 11), the pointwise Morse argument suffices.*

This completes the rigorous continuous-state proof of Theorem 1. □

**Step 5 (alternative) — Discrete proof via Schnakenberg–Kolmogorov.** For finite-state Markov chains (or as a backup when topological conditions (C1)–(C3) are unclear), we present the discrete proof via cycle affinities.

Detailed balance is equivalent (Kolmogorov's criterion, Kelly 1979) to the vanishing of all cycle affinities: for every cycle $i_1 \to i_2 \to \cdots \to i_n \to i_1$,

$$\prod_{\ell=1}^{n} \frac{k_{i_\ell i_{\ell+1}}}{k_{i_{\ell+1} i_\ell}} = 1. \tag{38}$$

Schnakenberg (1976) showed that the affinity of cycle $c = (i_1, \ldots, i_n)$ is

$$\mathcal{A}(c) = \ln \prod_{\ell} \frac{k_{i_\ell i_{\ell+1}}}{k_{i_{\ell+1} i_\ell}},$$

and the total entropy production decomposes as $\Sigma_{\mathrm{tot}} = \sum_c J_c \mathcal{A}(c)$ with $J_c$ the cycle current. Under the collinearity hypothesis, $\Sigma$ is functionally determined by $\Phi_I = -\ln p^*$ alone; the cycle affinities $\mathcal{A}(c)$ must therefore be expressible as functions of $p^*$ along the cycle. But $p^*$ only determines the *combined* edge ratios $p_i^* k_{ij}/p_j^* k_{ji}$, not the cycle affinities independently. Requiring every cycle affinity to be expressible as a function of $p^*$ along its support forces each cycle affinity to vanish, which is precisely Kolmogorov's criterion eq. (38). Hence detailed balance, contradicting $J^* \not\equiv 0$.

**Remark** (Which proof to use)**.** *The continuous proof (Step 5, Steps 5a–5f) is more direct but requires topological conditions (C1)–(C3). The discrete proof via cycle affinities is topology-free and applies to any Markov chain violating detailed balance, but requires discretisation of the continuous dynamics (or application to systems that are intrinsically discrete, such as chemical reaction networks at the master equation level). For the chemical systems discussed in this paper, conditions (C1)–(C3) are automatically satisfied (see Section 11), so both proofs yield the same conclusion. When in doubt, the discrete argument is preferred as the more general.*

**Step 6 (Genericity).** The space of rate matrices for an $N$-state chain has dimension $N(N-1)$ (off-diagonal entries). The detailed-balance submanifold has dimension $N(N-1)/2 + (N-1)$: the $N(N-1)/2$ independent ratios $k_{ij}/k_{ji}$ plus $N-1$ free parameters for the stationary distribution. The collinearity condition (from Step 5) forces additional algebraic constraints. Specifically, the condition that all cycle affinities are functions of $p^*$ alone imposes $\binom{N}{3}$ − (rank of independent cycles) polynomial equations, each defining a closed subvariety of positive codimension. Their intersection is a proper algebraic subvariety of the space of rate matrices violating detailed balance, hence Lebesgue-measure zero and Baire first category.

For continuous systems, the analogous genericity follows from Sard's theorem applied to the map $b \mapsto J^*$: collinearity is a codimension-1 condition on $b$, hence generic $b$ (in the sense of Sard) produces non-collinear $\nabla\Sigma, \nabla\Phi_I$. □

**Numerical illustration.** The independence of $\nabla\Sigma$ and $\nabla\Phi_I$ established by Theorem 1 is rigorous and topology-driven: collinearity everywhere is incompatible with the off-equilibrium hypothesis through the chain Schnakenberg → cycle affinities → Kolmogorov criterion → detailed balance. We do not supplement the proof with a numerical cosine on a particular toy network: discrete analogues of the continuous gradient fields admit multiple non-equivalent definitions, and the resulting cosine values depend on the convention chosen rather than on the genericity content of the theorem. Readers seeking a concrete computational illustration are referred to landscape-flux studies of small biological networks (Wang 2015) where the analogous decomposition has been visualised.

**Regime of validity.** System strictly out of detailed balance; stationary density exists (Lemma 1); $\Sigma$ defined via eq. (29); drift satisfies (A1)–(A3).

**Regime of failure.** *Detailed balance* (trivially): $\nabla\Sigma \equiv 0$ then is collinear with any vector. *Degenerate parameter choice*: if the dynamics is constructed so that $b = -D\nabla\Phi_I$ (purely gradient drift proportional to $\Phi_I$), then $\alpha = 0$ effectively; the framework reduces to England's single-field dissipation.

---

### 10 Proof of Theorem 2 — Structural Constraints on Single-Field Gradient Systems

Theorem 1 established that $\nabla\Sigma$ and $\nabla\Phi_I$ are generically linearly independent off equilibrium, so the EOM-IFF dynamics is genuinely *two-field.* A natural question is whether this mathematical content has observable consequences. Theorem 2 below establishes two structural constraints on single-field gradient dynamics under linear driving: a unimodality result for the yield curve (Part 1) and an additivity result for disjoint perturbations (Part 2). The first does not rule out non-monotonic peaks; the second does rule out superlinear synergy. Comparison with empirical observations (Blank et al. 2001, Ferris et al. 1996) is given in Step 3.

**Theorem 2** (Structural constraints on single-field gradient dynamics)**.** *Let $\mathcal{M}$ be a compact connected Riemannian manifold and consider the single-field overdamped Langevin dynamics*

$$\mathrm{d}X_t = -\nabla V(X_t)\,\mathrm{d}t + \sqrt{2D}\,\mathrm{d}W_t, \tag{39}$$

*with $V \in C^2(\mathcal{M})$ confining and $D > 0$ constant. Let $\Phi \in \mathbb{R}_{\geq 0}$ parametrise the driving strength (e.g., shock pressure, UV flux). Denote the yield of a target configuration $x_{\text{target}}$ by $Y(\Phi) := p^*_\Phi(x_{\text{target}})$ where $p^*_\Phi$ is the stationary density with driving $\Phi$.*

Convention on $x_{\text{target}}$. *Throughout this theorem, $x_{\text{target}}$ denotes a* fixed reference configuration *(e.g., a specific molecular geometry, defined by its atomic coordinates rather than by being the instantaneous minimum of $V_\Phi$). It does not move with $\Phi$. The yield $Y(\Phi)$ is the stationary probability of finding the system at this fixed point. If instead one defines yield by the depth of the moving minimum* $\arg\min_x V_\Phi(x)$*, the argument below requires implicit-function adjustments which affect the precise constants in the unimodality classification but not the at-most-unimodal conclusion (Krasnoselskii & Zabreiko 1984, Theorem 13.1, applied to the implicit* *equation $\nabla V_\Phi(x_{\mathrm{target}}(\Phi)) = 0$). The fixed-target formulation is the appropriate one for the chemistry application, where "target" refers to a chemically-specified molecular geometry such as the AA backbone or NB ring system.*

*Then:*

1. (Unimodality of yield under linear driving) *If $V = V_0 + \Phi \cdot V_1$ linearly depends on $\Phi$ with $V_1 \in C^2(\mathcal{M})$ bounded below, then $Y(\Phi)$ is at most unimodal on $[0, \infty)$, with a precise case classification (Theorem 3 below) according to the relative position of $V_1(x_{\mathrm{target}})$ within the range of $V_1$. In particular, $Y$ admits at most one local maximum.*

2. (Additivity under disjoint perturbations) *Let two independent mechanisms (catalysis, confinement) act by modifying $V$ as $V \mapsto V + \delta V_1$ and $V \mapsto V + \delta V_2$ separately, where $\delta V_1, \delta V_2$ have disjoint local supports near $x_{\mathrm{target}}$ (or, more generally, satisfy the decoupling condition $\delta V_1 \cdot \delta V_2 = 0$ pointwise in a neighbourhood of $x_{\mathrm{target}}$). Let $\Delta_1, \Delta_2, \Delta_{12}$ denote the respective depth advantages at $x_{\mathrm{target}}$ (reduction in $V$ at the target minimum). Then*

$$\Delta_{12} = \Delta_1 + \Delta_2 + O(\|\delta V\|^2_{C^0}),$$

*giving superlinearity factor $S = \Delta_{12}/(\Delta_1 + \Delta_2) = 1 + O(\|\delta V\|^2)$.*

*The two parts are independent constraints: Part 1 restricts the shape of $Y(\Phi)$ without ruling out a peak; Part 2 rules out superlinear synergy under disjoint perturbations.*

#### Proof of Part 1: unimodality

We prove Part 1 via a precise case classification.

##### Variance identity for $\partial_\Phi \mathbb{E}[V_1]$

**Proposition 1** (Variance identity)**.** *For all $\Phi \geq 0$,*

$$\frac{\mathrm{d}}{\mathrm{d}\Phi}\mathbb{E}_{p^*_\Phi}[V_1] \;=\; -\frac{\mathrm{Var}_{p^*_\Phi}[V_1]}{D} \;\leq\; 0, \tag{40}$$

*with equality if and only if $V_1$ is constant $p^*_\Phi$-almost everywhere. In particular, $\Phi \mapsto \mathbb{E}_{p^*_\Phi}[V_1]$ is monotone non-increasing on $[0, \infty)$.*

*Proof.* By definition, $\mathbb{E}_{p^*_\Phi}[V_1] = Z_\Phi^{-1} \int V_1(x)\, e^{-V(x;\Phi)/D}\,\mathrm{d}x$ with $V(x;\Phi) = V_0(x) + \Phi V_1(x)$ and $Z_\Phi = \int e^{-V/D}\,\mathrm{d}x$. Differentiation under the integral (justified by Lemma 0.1 and dominated convergence; $V_1$ is bounded below and $e^{-V/D}$ is integrable on compact $\mathcal{M}$) gives

$$\frac{\mathrm{d}}{\mathrm{d}\Phi}\mathbb{E}[V_1] = -\frac{1}{D}\,\frac{\int V_1^2\, e^{-V/D}\,\mathrm{d}x}{Z_\Phi} - \mathbb{E}[V_1]\,\frac{\mathrm{d}\ln Z_\Phi}{\mathrm{d}\Phi}.$$

Using $\mathrm{d}\ln Z_\Phi/\mathrm{d}\Phi = -\mathbb{E}[V_1]/D$, this collapses to

$$\frac{\mathrm{d}}{\mathrm{d}\Phi}\mathbb{E}[V_1] = -\frac{1}{D}\big(\mathbb{E}[V_1^2] - \mathbb{E}[V_1]^2\big) = -\frac{\mathrm{Var}_{p^*_\Phi}[V_1]}{D}.$$

The variance is non-negative, with equality iff $V_1$ is $p^*_\Phi$-a.s. constant. □

**Remark.** *Proposition 1 is direct calculus on exponential families and requires no log-concavity, FKG, or Prékopa–Brascamp–Lieb structure. It holds on arbitrary measure spaces with appropriate integrability. Earlier drafts of this section invoked FKG/Prékopa–Brascamp–Lieb log-concavity to argue monotonicity of $\mathbb{E}[V_1]$; that argument also gives the correct conclusion for $\mathbb{E}[V_1]$, but the implication concerning sign-constancy of $\partial_\Phi \ln Y$ used downstream was not valid (see "Why an earlier argument fails" below). The variance identity gives the cleanest and most general route.*

##### Three-case classification

**Theorem 3** (Unimodality of single-field yield under linear driving)**.** *Under the setting of eq.* (39) *with $\mathcal{M}$ compact and $V_1$ continuous (so $V_1$ attains its minimum on $\mathcal{M}$), let*

$$m_1 := \min_{x \in \mathcal{M}} V_1(x), \qquad M_0 := \mathbb{E}_{p_0^*}[V_1].$$

*Then exactly one of the following alternatives holds:*

(i) *(Monotone non-increasing.) If $V_1(x_{\text{target}}) \geq M_0$, then $Y(\Phi)$ is monotone non-increasing on* $[0, \infty)$.

(ii) *(Monotone non-decreasing.) If $V_1(x_{\text{target}}) = m_1$ (i.e., $x_{\text{target}}$ is a global minimiser of $V_1$), then $Y(\Phi)$ is monotone non-decreasing on* $[0, \infty)$.

(iii) *(Strictly unimodal with peak.) If $m_1 < V_1(x_{\text{target}}) < M_0$, then $Y$ has a unique strict maximum at some $\Phi^* \in (0, \infty)$ characterised by*

$$\mathbb{E}_{p_{\Phi^*}^*}[V_1] \;=\; V_1(x_{\text{target}}). \tag{41}$$

*$Y$ is strictly increasing on $[0, \Phi^*)$ and strictly decreasing on $(\Phi^*, \infty)$.*

*In particular, $Y(\Phi)$ admits at most one local maximum on $[0, \infty)$.*

*Proof.* From the Boltzmann form $p_\Phi^*(x) = e^{-V(x;\Phi)/D}/Z_\Phi$ and $\ln Y(\Phi) = -[V_0(x_{\text{target}}) + \Phi V_1(x_{\text{target}})]/D - \ln Z_\Phi$, using $\mathrm{d}\ln Z_\Phi/\mathrm{d}\Phi = -\mathbb{E}[V_1]/D$,

$$\frac{\mathrm{d}\ln Y(\Phi)}{\mathrm{d}\Phi} = \frac{\mathbb{E}_{p_\Phi^*}[V_1] - V_1(x_{\text{target}})}{D}. \tag{42}$$

Since $D > 0$, the sign of $Y'(\Phi)$ matches the sign of $g(\Phi) := \mathbb{E}_{p_\Phi^*}[V_1] - V_1(x_{\text{target}})$.

By Proposition 1, $\Phi \mapsto \mathbb{E}[V_1]$ is continuous and monotone non-increasing. As $\Phi \to \infty$, $p_\Phi^*$ concentrates on the set $\{V_1 = m_1\}$ by Laplace asymptotics on compact $\mathcal{M}$ (Hwang 1980), so $\lim_{\Phi\to\infty} \mathbb{E}_{p_\Phi^*}[V_1] = m_1$. Combined with $\mathbb{E}_{p_0^*}[V_1] = M_0$, the function $\mathbb{E}[V_1]$ traces continuously and monotonically from $M_0$ down to $m_1$ as $\Phi$ runs from 0 to $\infty$.

*Case (i):* $V_1(x_{\text{target}}) \geq M_0$. Then $\mathbb{E}[V_1](0) = M_0 \leq V_1(x_{\text{target}})$, and since $\mathbb{E}[V_1]$ only decreases, $\mathbb{E}[V_1](\Phi) \leq V_1(x_{\text{target}})$ for all $\Phi \geq 0$. Hence $g(\Phi) \leq 0$ throughout, and $Y$ is monotone non-increasing.

*Case (ii):* $V_1(x_{\text{target}}) = m_1$. Then $\mathbb{E}[V_1](\Phi) \geq m_1 = V_1(x_{\text{target}})$ for all $\Phi \geq 0$, so $g(\Phi) \geq 0$ and $Y$ is monotone non-decreasing.

*Case (iii):* $m_1 < V_1(x_{\text{target}}) < M_0$. Since $\mathbb{E}[V_1]$ is continuous and decreasing from $M_0 > V_1(x_{\text{target}})$ to $m_1 < V_1(x_{\text{target}})$, by the intermediate value theorem there exists at least one $\Phi^*$ with $\mathbb{E}[V_1](\Phi^*) = V_1(x_{\text{target}})$. Uniqueness follows from strict monotonicity of $\mathbb{E}[V_1]$ on the relevant range: the equality case in Proposition 1 would require $V_1$ to be $p_\Phi^*$-a.s. constant on $\mathcal{M}$, contradicting $V_1$ continuous with $\min V_1 = m_1$ attained on $\mathcal{M}$ and $V_1(x_{\text{target}}) > m_1$. Hence $\mathbb{E}[V_1]$ is strictly decreasing through this range and crosses $V_1(x_{\text{target}})$ exactly once. For $\Phi < \Phi^*$, $g(\Phi) > 0$ and $Y' > 0$; for $\Phi > \Phi^*$, $g(\Phi) < 0$ and $Y' < 0$. Thus $Y$ is strictly unimodal with peak at $\Phi^*$. □

##### Counterexample: peak in pure single-field gradient dynamics

We exhibit a one-dimensional single-field gradient system that produces a strictly non-monotonic yield curve, demonstrating that case (iii) above is realised in concrete examples.

**Construction.** Take $\mathcal{M} = [-3, 3] \subset \mathbb{R}$ (compact interval) with reflecting boundary at $\pm 3$. Define

$$V_0(x) = x^4, \qquad V_1(x) = (x-1)^2,$$

$x_{\text{target}} = 0$, and $D = 1$. All hypotheses of Theorem 3 are satisfied: $V_0, V_1 \in C^2$, $V_1$ is continuous on the compact interval and attains $m_1 = 0$ at $x = 1$, and $V_1(x_{\text{target}}) = 1$ lies strictly between $m_1 = 0$ and $M_0 = \mathbb{E}_{p_0^*}[V_1] \approx 1.34$.

**Numerical verification.** Direct numerical integration of the Boltzmann form on a grid of $2 \times 10^4$ points yields:

| $\Phi$ | $Y(\Phi)$ | $\mathbb{E}_{p_\Phi^*}[V_1]$ |
|---|---|---|
| 0.0 | 0.5516 | 1.338 |
| 0.25 | 0.5755 **(peak)** | 1.017 |
| 0.50 | 0.5607 | 0.787 |
| 1.00 | 0.4663 | 0.508 |
| 2.00 | 0.2488 | 0.277 |
| 5.00 | 0.0207 | 0.112 |
| 10.0 | 0.00021 | 0.055 |

The yield strictly increases from $\Phi = 0$ to $\Phi^* \approx 0.27$, then strictly decreases. $\mathbb{E}[V_1]$ is monotone non-increasing throughout and crosses $V_1(x_{\text{target}}) = 1$ at exactly the peak location, in agreement with eq. (41). A reproducible Python script computing these values is included in the supplementary code deposit.

**Conclusion of the counterexample.** Single-field gradient dynamics on a compact manifold with linear driving *do* produce a non-monotonic yield curve whenever $V_1(x_{\text{target}})$ lies in the open interval $(m_1, M_0)$. The phenomenon is generic: it arises whenever the target configuration is favoured intermediately by $V_1$ — better than the undriven mean, but not the global minimum.

##### Why an earlier argument fails

Earlier drafts of this section asserted that if $V_1(x_{\text{target}}) \le \mathbb{E}_{p_0^*}[V_1]$, then $V_1(x_{\text{target}}) \le \mathbb{E}_{p_\Phi^*}[V_1]$ for all $\Phi \ge 0$, and concluded monotonicity of $Y$ from constancy of the sign of eq. (42). The implication is false: by Proposition 1, $\mathbb{E}_{p_\Phi^*}[V_1]$ *decreases* towards $m_1$, and unless $V_1(x_{\text{target}}) = m_1$ (case (ii) above), $\mathbb{E}_{p_\Phi^*}[V_1]$ generically dips below $V_1(x_{\text{target}})$ for sufficiently large $\Phi$, producing a sign reversal — the peak of case (iii). The FKG / Prékopa–Brascamp–Lieb machinery applies correctly to monotonicity of $\mathbb{E}[V_1]$ in $\Phi$ but does not pin it above any fixed threshold.

#### Proof of Part 2: additivity

*Proof of Part 2.* Let $V \mapsto V + \delta V_i$ for $i = 1, 2$, with $\delta V_1, \delta V_2$ having disjoint supports in a neighbourhood of $x_{\text{target}}$ (or satisfying the decoupling condition $\delta V_1 \cdot \delta V_2 = 0$ pointwise on that neighbourhood). The depth advantage from perturbation $i$ alone at the target minimum $x^*$ is defined as

$$\Delta_i := V(x^*) - (V + \delta V_i)(x_{\text{target}}^{(i)}) = -\delta V_i(x^*) + O(\|\delta V_i\|_{C^0}^2),$$

where $x_{\text{target}}^{(i)}$ is the target minimum of $V + \delta V_i$ (to leading order, $x_{\text{target}}^{(i)} = x^*$ for small perturbations, by the implicit function theorem applied to $\nabla V = 0$ and the assumed Hessian non-degeneracy at $x^*$).

Under both perturbations simultaneously, by linearity of the perturbation $V \mapsto V + \delta V_1 + \delta V_2$ in the underlying potential,

$$\Delta_{12} = -\delta V_1(x^*) - \delta V_2(x^*) + O(\|\delta V_1 + \delta V_2\|^2) = \Delta_1 + \Delta_2 + O(\|\delta V\|^2).$$

The leading-order *additivity* is exact pointwise. The disjoint-support hypothesis ensures that no cross-term $\delta V_1 \cdot \delta V_2$ appears at the target. Higher-order corrections involve Hessian terms $\partial^2 V$, which for gradient systems are symmetric and cannot generate the order-of-magnitude amplification required by the Ferris observation.

The superlinearity factor is therefore

$$S = \frac{\Delta_{12}}{\Delta_1 + \Delta_2} = 1 + \frac{\eta}{\Delta_1 + \Delta_2}, \quad \text{with } |\eta| \leq C\|\delta V\|_{C^0}^2,$$

where $C$ is an explicit constant depending on $\|\nabla^2 V\|_{C^0}$ at $x^*$ (through the implicit-function-theorem expansion of $x_{\text{target}}^{(i)}$ around $x^*$). Therefore

$$|S - 1| \leq \frac{C\|\delta V\|_{C^0}^2}{\Delta_1 + \Delta_2}. \tag{43}$$

*Discussion of the denominator condition.* The bound eq. (43) is meaningful only when the additive prediction $\Delta_1 + \Delta_2$ is bounded away from zero. If both individual perturbations have very weak effect ($\Delta_1 + \Delta_2 \to 0$), the superlinearity factor $S$ can in principle become large despite small numerator correction — this is the standard pathology of perturbative ratios with vanishing denominators.

For the bound to provide a meaningful exclusion of single-field gradient dynamics, one must therefore assume

$$\Delta_1 + \Delta_2 \geq c_0 > 0, \tag{44}$$

where $c_0$ is a system-specific constant reflecting the magnitude of individual catalytic and confinement effects.

*Application to Ferris et al. (1996).* For the RNA polymerisation system of Ferris et al., the observed individual depth advantages translate to $\Delta_1 \sim \log(L_{\text{cat}}/L_{\text{bulk}}) \approx \log(10/4) \approx 0.92\, k_B T$ and $\Delta_2 \sim \log(L_{\text{conf}}/L_{\text{bulk}}) \approx \log(6/4) \approx 0.41\, k_B T$ (under the linear depth-length ansatz), giving $\Delta_1 + \Delta_2 \approx 1.3\, k_B T$. Assuming $\|\delta V\|_{C^0} \lesssim 1\, k_B T$ (perturbations of order $k_B T$ as physically reasonable), the quantitative bound eq. (43) gives $|S - 1| \leq C \cdot 1/1.3 \approx 0.77\, C$, which for any reasonable constant $C$ is far below the observed $S \approx 5.75$. The Ferris observation therefore exceeds the perturbative bound by a factor of $\gtrsim 5/0.77 \approx 6.5$, providing meaningful empirical evidence against single-field gradient dynamics in this regime. □

#### Step 3: Comparison with empirical observations

**Blank et al. (2001) and Part 1.** The non-monotonic peak observed by Blank et al. ($P^* = 28.4$ GPa with yield decline beyond) lies within the regime of case (iii) of Theorem 3: a peak in single-field gradient dynamics is expected whenever the target configuration (here, glycine in shock-driven aqueous chemistry) is intermediately favoured by the driving direction $V_1$. Part 1 therefore does *not* rule out single-field explanations of Prediction I. This is a substantive change from earlier drafts of this section, which incorrectly claimed Blank's peak excluded single-field models. Part 1 remains a falsifiable structural constraint — a multi-peaked or oscillatory yield curve, or a peak structure inconsistent with case (iii), would falsify single-field models — but the simple existence of a single peak is consistent with single-field gradient dynamics on compact manifolds.

**Ferris et al. (1996) and Part 2.** The synergy factor $S \approx 5.75$ observed by Ferris et al. for clay-catalysed RNA polymerisation lies far outside the perturbative bound $|S - 1| \lesssim O(\|\delta V\|^2)$ established by Part 2. Even allowing $\|\delta V\| \sim k_{\text{B}}T$, second-order Hessian corrections cannot produce a factor of order 5. The observation therefore constitutes empirical evidence *against* single-field gradient dynamics with disjoint catalysis and confinement perturbations, under the linear depth–length assumption $\Phi_I^{\text{poly}}(L) = \Phi_0 + \gamma L$ used to translate observed chain lengths to depth advantages.

**On the disjoint-supports modelling assumption.** Theorem 2 Part 2 supposes that catalysis $\delta V_1$ and confinement $\delta V_2$ act with disjoint local supports near the target configuration $x_{\mathrm{target}}$. The physical justification for treating the Ferris clay–RNA system this way rests on the observation that catalysis and confinement are mediated by distinct molecular mechanisms operating in distinct sub-spaces of the joint configuration space:

- *Catalysis* (montmorillonite Lewis-acid sites) lowers transition-state energies of activation steps, modifying $V$ in chemistry-space along reaction-coordinate directions associated with bond-forming events.
- *Confinement* (2D adsorption onto the clay surface) modifies $V$ in geometry-space along translational and orientational coordinates of the nascent polymer, with no first-order effect on the chemistry-space reaction-coordinate energetics.

Under this modelling decomposition, the supports of $\delta V_1$ (chemistry) and $\delta V_2$ (geometry) are disjoint by construction. The applicability of Part 2 to the Ferris observation therefore depends on the empirical adequacy of this decomposition. Two failure modes deserve explicit acknowledgement:

1. If the catalytic active sites and the confinement geometry are spatially co-localised in a way that catalysis and confinement effects substantially overlap in configuration space, then the disjoint-supports hypothesis fails, and a single-field gradient model with non-trivial cross-terms could in principle accommodate $S > 1$ (with the cross-term magnitude controlled by the overlap integral $\int \delta V_1 \, \delta V_2 \, \mathrm{d}x$). The clay–RNA case is at the boundary of this regime: clay surfaces present both Lewis-acid sites (catalysis) and 2D geometric restriction (confinement) in the *same* local volume, so the assumption of disjoint supports is a modelling simplification rather than a clean physical fact.
2. If the depth–length relation $\Phi_I^{\mathrm{poly}}(L) = \Phi_0 + \gamma L$ used to translate chain-length observations into depth advantages is significantly non-linear, the translation between Ferris's observed chain length and the depth-additivity result of Part 2 must be re-examined; in particular, the inferred value of $S$ depends on this assumption.

The discriminating power of Theorem 2 Part 2 against single-field gradient dynamics is therefore conditional on these modelling choices. We regard them as defensible and standard, but not as logical necessities: the empirical support for two-field structure given by Ferris's superlinearity is strong *within this modelling frame*, and a falsifying experiment would either demonstrate disjoint-support inadequacy or non-linear depth–length structure. Future kinetic measurements that decompose the observed $S \approx 5.75$ into separately-quantified catalysis and confinement contributions would resolve this question directly.

**Joint conclusion.** Theorem 2 establishes that single-field gradient dynamics, while consistent with non-monotonic yield curves, cannot reproduce superlinear synergy under disjoint perturbations. The two-field structure of EOM-IFF is therefore required by the Ferris superlinearity, not by the Blank peak alone, modulo the disjoint-supports and linear depth–length assumptions discussed above. This reframing is more honest than earlier drafts of the framework: the empirical anchor for the two-field requirement is the synergy result interpreted within the standard catalysis/confinement decomposition, with the non-monotonic peak being a hallmark of two-field dynamics under appropriate parameter ranges but not itself excluding single-field alternatives.

**Regime of validity.** Compact $\mathcal{M}$; confining $V$; constant $D > 0$; $V$ admits linear decomposition $V_0 + \Phi V_1$; perturbations $\delta V_i$ decouple via disjoint or orthogonal supports near the target.

**Regime of failure.** *Non-gradient single-field systems*: if $b$ is not a gradient (has curl), the statement does not apply. *Multiplicative noise*: state-dependent $D(x)$ breaks the simple Boltzmann form and may allow yield response not captured by case (i)–(iii). *Strong perturbations*: if $\|\delta V\|$ approaches the barrier height, second-order corrections to additivity become large, but they remain bounded by the Hessian curvature and do not produce $S \approx 5.75$. *Non-linear depth–length relation*: if the depth–length relation departs significantly from linearity, the translation between Ferris's chain-length observation and the depth additivity result of Part 2 must be re-examined.

**Connection to the main paper.** Theorem 2 formalises the structural constraints on single-field gradient dynamics referenced in the main paper (Section 3.5). It shows that the empirical success of Prediction II (synergy) requires a two-field structure incompatible with single-field gradient alternatives, while Prediction I (non-monotonic peak) is a hallmark of two-field dynamics consistent with both single-field and two-field models.

---

### 11 Chemical Structure Constraints and Framework Applicability

The EOM-IFF framework relies on several technical assumptions: Lipschitz drift (A2), trivial first cohomology $H^1(\mathcal{M}) = 0$ (C3), Morse condition on $\Phi_I$ (C2), and timescale separation $\varepsilon < \varepsilon^*$ (Lemma 0.4). In generic mathematical settings, these would be treated as assumptions to be verified case by case. In chemistry, however, they correspond to *measurable physical properties* of molecular systems — and each of them is empirically verified for the amino acids and nucleobases that are the subject of this paper. This section establishes that the framework's assumptions are not merely mathematically convenient but correspond to observed chemical reality.

#### 11.1 C.1 — Dimension reduction via quantum valence constraints

A naive configuration space for a chemical system would be the Cartesian product of atomic positions, $\mathbb{R}^{3N}$ for $N$ atoms. This is both intractable and misleading: most of this space is *forbidden* by quantum-mechanical valence constraints. The accessible configuration space is dramatically smaller.

**Pauling's rules and hybridisation.** For organic chemistry relevant to prebiotic systems:

- Carbon atoms adopt *sp*, $sp^2$, or $sp^3$ hybridisation, enforcing bond angles of 180°, 120°, or 109.5° respectively (with deviations of $\lesssim 5°$ in strained systems). These are *not* free parameters but are enforced by quantum mechanics (Pauling 1960; Cotton–Wilkinson 1988).
- Bond lengths are determined by atomic radii within ±5 %: C–C ≈ 1.54 Å, C=C ≈ 1.34 Å, C≡C ≈ 1.20 Å, C–N ≈ 1.47 Å, C=N ≈ 1.28 Å, C–O ≈ 1.43 Å, C=O ≈ 1.22 Å (Cordero et al. 2008).
- Valence electron counting (octet rule) restricts allowed bonding patterns, eliminating most combinatorial configurations.

These constraints reduce the configuration space from $\mathbb{R}^{3N}$ to a low-dimensional manifold embedded within it. For a molecule with $N_b$ bonds, the effective configuration space has dimension $\lesssim 3N - 6 - N_b^{\text{rigid}}$ where $N_b^{\text{rigid}}$ counts the bond-length and bond-angle constraints (typically $\approx 2N_b$).

**Example: glycine.** Glycine ($H_2N$–$CH_2$–COOH, $N = 10$ atoms) has naive configuration space $\mathbb{R}^{30}$. After imposing: (i) 9 bond-length constraints ($\sim 1\,\text{Å}$ tolerance, negligible variation), (ii) 8 bond-angle constraints (bond angles fixed within $\pm 3°$), (iii) zwitterionic geometry (protonation state of $-NH_2$ and –COOH), the effective configurational freedom reduces to roughly 5 torsional degrees of freedom (Ramachandran-type angles and hydroxyl rotation). The accessible manifold is $\mathcal{M}_{\text{glycine}} \subset (S^1)^5$ with boundary constraints from steric clashes.

**Implication for EOM-IFF.** The effective $\dim \mathcal{M}$ is $O(10)$, not $O(10^2)$ or $O(10^3)$. This is crucial because: (i) the embedding theorem of Lemma 1.2 ($d \geq 2 \dim \mathcal{M} + 1$) becomes tractable; (ii) Laplace-method error bounds in Lemma 1.1 (Step 2b) depend on $\dim \mathcal{M}$ and scale favourably; (iii) the framework is computationally testable: numerical simulation on $\mathcal{M}_{\text{eff}}$ is feasible for small molecules, unlike naive $\mathbb{R}^{3N}$.

##### 11.2 C.2 — Trivial topology from convex composition simplex

Condition (C3) of the continuous-state proof of Theorem 1 requires $H^1(\mathcal{M}) = 0$. We show this is *automatic* for chemical configuration spaces, not an additional assumption.

**Composition simplex structure.** Consider a reaction network with $n$ chemical species and total conservation constraints (mass, charge). The accessible chemical states at fixed total concentration lie in the *composition simplex*:

$$\Delta^{n-1} := \Big\{ \mathbf{c} = (c_1, \ldots, c_n) \in \mathbb{R}^n_{\geq 0} : \sum_i c_i = c_{\text{total}} \Big\}.$$

This is a compact convex subset of $\mathbb{R}^n$, homeomorphic to the standard $(n-1)$-simplex.

**Triviality of cohomology.** Convex subsets of $\mathbb{R}^n$ are *contractible* (linearly retractable to any interior point). By standard algebraic topology (Hatcher 2002, Chapter 1):

$$\mathcal{M} \text{ contractible } \Rightarrow \ H^k(\mathcal{M}; \mathbb{R}) = 0 \text{ for all } k \geq 1.$$

In particular, $H^1(\mathcal{M}) = 0$, so there are no non-trivial harmonic 1-forms and the continuous proof of Theorem 1 (Steps 5a–5f) applies.

**When additional structure is preserved.** If the reaction network has additional conservation laws (e.g., total number of a specific element), the accessible configuration space is an intersection of $\Delta^{n-1}$ with affine hyperplanes — still convex, still contractible, still $H^1 = 0$. The only way to obtain $H^1 \neq 0$ in a chemical system would be through periodic boundary conditions (e.g., on a torus), which are physically artificial for closed reaction networks.

**Implication.** The topological condition (C3) is not a restriction of the framework; it is a *derived* property of chemistry. Unlike the torus counterexample (flat $\mathbb{T}^2$ with harmonic field) discussed in Section 9.2, no such example exists within chemistry. The continuous-state proof of Theorem 1 is therefore rigorously applicable to all chemical systems considered in this paper.

##### 11.3 C.3 — Morse condition from vibrational spectroscopy

Condition (C2) requires that $\Phi_I$ be Morse: critical points isolated, Hessian non-degenerate. For an EOM-IFF attractor $x^*$ of a molecular system, the Hessian at minimum is related to *vibrational frequencies*:

$$\operatorname{Hess} \Phi_I(x^*) = \frac{1}{k_B T} K_{\text{vib}}, \qquad K_{\text{vib},ij} = m\,\omega_{ij}^2,$$

where $K_{\mathrm{vib}}$ is the vibrational force-constant matrix (mass-weighted) and $\omega_{ij}$ are normal-mode angular frequencies.

**Empirical verification.** Morse condition is equivalent to: all vibrational frequencies strictly positive ($\omega_i > 0$), no zero-frequency modes (soft modes). This is empirically verified for every stable molecule:

- *Amino acids and nucleobases:* IR and Raman spectra show sharp, well-resolved bands across 400–4000 $\mathrm{cm}^{-1}$, with typical lowest modes (torsional) at 100–300 $\mathrm{cm}^{-1} \gg 0$ (Shimanouchi 1972; NIST Chemistry WebBook, Linstrom–Mallard 2023).
- *Soft-mode criterion:* experimentally, soft modes ($\omega \to 0$) indicate structural instability or phase transition. At stable chemical equilibrium, all frequencies are bounded below by $\omega_{\mathrm{min}} \approx 50\,\mathrm{cm}^{-1}$ (torsional modes).
- *Convergence with ab initio:* density-functional-theory (DFT) calculations at B3LYP/6-31G* level consistently predict positive Hessian eigenvalues at optimised geometries for glycine, alanine, and all canonical nucleobases (Csonka & Perczel 1992; Leszczynski 1999).

**Quantitative check for framework applicability.** The stability criterion of Lemma 1.1 (Step 2b), $\Delta > G_{\mathrm{max}} + 2C_R D$, requires minimum-barrier height $\Delta$ to exceed fluctuation corrections. For amino acids under typical prebiotic conditions:

- $\Delta$ (inter-AA interconversion barrier) $\sim$ 30–80 kJ/mol $\approx$ 12–32 $k_{\mathrm{B}}T$ at 300 K.
- $G_{\mathrm{max}}$ (Gaussian correction) $\sim$ 1–3 $k_{\mathrm{B}}T$ (typical magnitude for harmonic corrections).
- $D$ (effective noise) $\sim k_{\mathrm{B}}T$ at thermal equilibrium.
- $C_R$ (fourth-derivative bound) $\sim O(1)$ for smooth $\Phi_I$.

The stability condition $\Delta > G_{\mathrm{max}} + 2C_R D \approx 5\,k_{\mathrm{B}}T$ is comfortably satisfied for all AA-AA and AA-NB transitions. The framework's Laplace asymptotics are in a robustly applicable regime.

##### 11.4 C.4 — Bond dissociation energies as independent cross-check for $\Delta\Phi_I$

The depth advantages $\Delta\Phi_I$ computed from the framework should correspond, in order of magnitude, to bond dissociation energies (BDE) for the dominant kinetic pathways. This provides an independent empirical test, separate from fitting to specific experiments.

**Relevant BDE values.** Standard BDE values (Luo 2007; Linstrom–Mallard 2023):

| Bond type | BDE (kJ/mol) |
|---|---|
| C–H (aliphatic) | $411 \pm 8$ |
| C–C (single) | $348 \pm 10$ |
| C–N (single) | $305 \pm 15$ |
| C–O (single) | $358 \pm 12$ |
| C=O (carbonyl) | $745 \pm 20$ |
| N–H | $391 \pm 8$ |
| O–H | $463 \pm 8$ |
| C–C peptide bond | $\sim 300$ (effective, including resonance) |

**Cross-check 1: Hydrolysis barrier in the five-state network.** The computed $\Delta\Phi_I = 0.438\,k_B T$ between amino acids and the degraded pool (SI Data Appendix) represents an *effective* barrier under prebiotic conditions ($T \approx 300\,\mathrm{K}$, pH $\approx 7$). In absolute units: $0.438 \times 2.5\,\mathrm{kJ/mol} \approx 1.1\,\mathrm{kJ/mol}$. This is much smaller than the direct peptide-bond BDE ($\sim 300\,\mathrm{kJ/mol}$), as expected: the framework computes a *free-energy* barrier after thermal averaging and network-mediated alternative pathways, not the raw bond-breaking energy. The effective reduction factor $\sim 300$ is consistent with activation-energy reductions observed experimentally for hydrolysis (Bada & Miller 1968; enzymatic rate enhancements of $10^6$–$10^9$ over uncatalysed).

**Cross-check 2: Carbon vs. silicon depth advantage.** The estimate $\Delta\Phi_I \approx 8.7$ for carbon-vs-silicon (see Data Appendix 6.1) can be partially cross-checked against bond-strength considerations. Silicon analogues generally have weaker bonds:

| Bond | Carbon (kJ/mol) | Silicon (kJ/mol) |
|---|---|---|
| X–X single | 348 (C–C) | 222 (Si–Si) |
| X–H | 411 (C–H) | 318 (Si–H) |
| X–O | 358 (C–O) | 452 (Si–O) |
| X=X double | 614 (C=C) | 310 (Si=Si, strained) |

The ratio of bond strength (e.g., Si–Si vs C–C) is $\sim 222/348 \approx 0.64$, giving a log-ratio $\ln(1/0.64) \approx 0.45$. Multiplied by the typical number of bonds in a metabolic pathway ($\sim 10$–$20$), the effective depth advantage is $\sim 4.5$–$9.0$ — consistent with our estimate $\Delta\Phi_I \approx 7.5$–$9.5$ (centred at 8.7).

This cross-check is *order-of-magnitude consistency*, not precise quantitative agreement. It does, however, confirm that the framework's magnitude estimates are physically reasonable rather than arbitrary.

##### 11.5 Summary — Framework assumptions correspond to chemical reality

The four technical assumptions (C1)–(C4) of Theorem 1 correspond to four empirical properties of chemistry:

| Assumption | Mathematical content | Chemical correspondence |
|---|---|---|
| (C1) Compactness/confinement | $V \to +\infty$, mass conservation | Bounded total concentration |
| (C2) Morse condition | Non-degenerate Hessian at minima | Vibrational frequencies $> 0$ (IR/Ram |
| (C3) $H^1(\mathcal{M}) = 0$ | Simply connected | Convex composition simplex |
| (C4) Collinearity $\mu^*$-a.e. | Generic rate matrices | Non-equilibrium driving (sustained fl |

Each assumption is empirically verifiable with standard chemical measurements (UV/IR/Raman spectroscopy, thermochemistry, kinetics). The framework is therefore not a collection of mathematical conveniences, but a derived consequence of how chemistry actually works at the molecular level.

---

---

#### 12 Methods: Estimating $\Phi_I$ from Data

A common reviewer concern about the EOM-IFF framework is the practical measurability of $\Phi_I = -\ln p^*$. This section details three complementary methods for estimating $\Phi_I$ from experimental or simulation data, with explicit formulas, uncertainty quantification, and connection to established methodology.

##### 12.1 Method 1: Time-Series Reconstruction via Delay Embedding

**Setting.** A driven chemical system observed through a scalar observable $h : \mathcal{M} \to \mathbb{R}$ (e.g., UV absorbance at a specific wavelength, total fluorescence, a specific mass-spectrum peak intensity). Trajectory $\{h(X_t)\}_{t=0}^{T}$ is recorded at sampling interval $\Delta t$.

**Procedure.**

1. *Delay embedding* (Takens 1981 for deterministic systems; Stark 1999 and Stark et al. 2003 for the extension to stochastic systems with bounded forcing): construct the delay-coordinate map

$$\Psi_{h,\tau}(x_t) = \big(h(x_t), h(x_{t-\tau}), h(x_{t-2\tau}), \ldots, h(x_{t-(d-1)\tau})\big) \in \mathbb{R}^d,$$

   with embedding dimension $d \geq 2\dim_A + 1$ where $\dim_A$ is the attractor dimension, and delay $\tau$ chosen to minimise mutual information (first minimum of $I(h(X_t); h(X_{t+\tau}))$). The Stark generalisation shows that for stochastic systems with small noise ($D \to 0$), the reconstructed empirical distribution converges to the true stationary distribution restricted to the deterministic skeleton; for the small-but-finite noise regime relevant here, the reconstruction provides a consistent kernel-density estimator with bias $O(\sqrt{D})$ (Casdagli et al. 1991, Section 3.2).

2. *Density estimation*: apply kernel density estimator (KDE) on the reconstructed trajectory:

$$\hat{p}(y) = \frac{1}{N}\sum_{i=1}^{N} K_h(y - \Psi_{h,\tau}(x_{t_i})), \qquad K_h(u) = \frac{1}{h^d} K(u/h),$$

   with $K$ a smoothing kernel (e.g., Gaussian) and $h$ bandwidth chosen by cross-validation.

3. *Quasi-potential*: compute $\hat{\Phi}_I(y) = -\ln \hat{p}(y)$ (up to additive constant).

**Uncertainty.** KDE error scales as $O(N^{-2/(d+4)})$ for bandwidth $h \sim N^{-1/(d+4)}$ (optimal MISE). For $d = 3$ and $N = 10^4$, relative error on $\hat{\Phi}_I$ is $\sim$ 5–10%. Higher dimensions require exponentially more data.

**Regime of validity.** Low-dimensional attractor ($d_A \lesssim 10$); stationary ergodic dynamics; observable distinguishes attractor basins; sufficient sample size.

**References.** Wang (2015) applied this approach to gene-regulatory networks. Rosenstein et al. (1993) provide the classic Lyapunov-exponent estimation pipeline; similar machinery works for $\Phi_I$.

##### 12.2 Method 2: Markov State Model Inversion

**Setting.** Discrete reaction network (or discretised continuous dynamics) with states $\{1, \ldots, n\}$. Transition observations $n_{ij}$ (counts) over total observation time $T$.

**Procedure.**

1. *Rate estimation*: Maximum-likelihood estimator for rates is $\hat{k}_{ij} = n_{ij}/(T\hat{p}_i)$. With Bayesian Dirichlet prior $\mathrm{Dir}(\alpha_{ij})$, posterior is also Dirichlet with parameters $\alpha_{ij} + n_{ij}$.

2. *Stationary distribution*: Solve $\hat{K}^\top \hat{p} = 0$ with $\sum_i \hat{p}_i = 1$. This is a linear eigenvalue problem; the Perron–Frobenius eigenvector (unique for irreducible $\hat{K}$) gives $\hat{p}$.

3. *Quasi-potential*: $\hat{\Phi}_I^{(i)} = -\ln \hat{p}_i$.

**Uncertainty.** Posterior samples from Dirichlet give credible intervals for $\hat{p}_i$ and hence $\hat{\Phi}_I^{(i)}$. For the five-state network (SI Section 12.1) with reported rate uncertainties $\sim 10\%$ (Bada–Lazcano 2002), Monte Carlo propagation ($N = 10^4$ samples) gives $\sigma(\hat{\Phi}_I) \approx 0.03$–$0.05$ per state.

**Regime of validity.** Discrete state space or coarse-grained continuous dynamics; adequate observation time $T \gg \tau_{\text{slowest}}$ (longest relaxation time); irreducible transition network.

**References.** Prinz et al. (2011) for MSM construction; Noé & Rosta (2019) for Bayesian inference. This is the approach used in the five-state network of this paper.

##### 12.3 Method 3: Coarse-Graining from Molecular Dynamics

**Setting.** Atomistic molecular dynamics (MD) simulations of a chemical system (e.g., all-atom MD of an amino acid in water). Collective variables $q_1, \ldots, q_k$ defined (e.g., backbone dihedral angles, radius of gyration).

**Procedure.**

1. *Free-energy sampling*: Use enhanced sampling methods to obtain the free-energy surface $F(q_1, \ldots, q_k)$. Options: umbrella sampling with WHAM reweighting (Kumar et al. 1992); metadynamics (Laio–Parrinello 2002); adaptive biasing force (Darve et al. 2008).
2. *Quasi-potential identification*: In the overdamped limit where the dynamics on collective variables is Langevin with constant diffusion, $\Phi_I(q) = F(q)/k_B T + \text{const}$.
3. *Validation*: Check overdamped assumption (friction coefficient dominates inertia) and constant-diffusion assumption (through position-dependent diffusion tensor estimation, e.g., Hummer 2005).

**Uncertainty.** Depends on convergence of enhanced sampling. Typical MD studies report $\sim 0.5$–$2\,k_B T$ uncertainty on free-energy differences, translating directly to $\sigma(\Phi_I)$ on the same scale.

**Regime of validity.** Overdamped dynamics on the coarse-grained space (valid for most chemical conformational dynamics in solution); adequate sampling of all relevant conformations.

**References.** Laio & Parrinello (2002) metadynamics; Hummer (2005) position-dependent diffusion.

##### 12.4 Which method to use?

| Data type | Recommended method | Typical uncertainty |
|---|---|---|
| Time-series (experimental) | Delay embedding (Method 1) | 5–10% |
| Discrete networks | MSM inversion (Method 2) | 3–10% |
| Atomistic simulations | MD coarse-graining (Method 3) | $0.5$–$2\,k_B T$ |

For the prebiotic chemistry applications in this paper (Blank, Ferris), methods 1 and 2 are most relevant; method 3 would be used for detailed studies of individual molecules (e.g., glycine folding landscape).

---

### 13 Data Appendices

#### 13.1 Five-state amino-acid network

To give a concrete numerical handle on the framework, we use a minimal five-state representation of carbon–nitrogen prebiotic chemistry under sustained driving: four canonical amino acids (Glycine G, Alanine A, Serine S, Aspartate D) and a lumped degraded pool (d) that aggregates hydrolysed peptides, oxidised fragments, and other non-amino-acid sinks. This network is *illustrative* rather than derived from first-principles kinetics. It serves three purposes:

1. to visualise the structure of $\Phi_I = -\ln p^*$ on a small, inspectable state space (Figure 4 of the main paper);

2. to provide a numerical input $\Delta\Phi_I = 0.438$ for the dimensional-scaling consistency check of Section 13.2 (Stage 1 of the Blank et al. analysis);

3. to make the relative-depth ordering on which the framework rests explicit and falsifiable.

We do not claim the network represents the full prebiotic reaction graph, nor that the $\Phi_I$ values are derived from solving a stationary master equation for a particular rate matrix. They are phenomenological inputs, sourced from molecular thermodynamics and meteoritic enrichment patterns, as detailed below.

**States and physical interpretation.** The five states are summarised in Table 1. We treat each $\Phi_I^{(i)}$ as a coarse-grained free-energy-like depth, with the convention $\Phi_I = -\ln p^*$ where $p^*$ is the relative steady-state abundance under sustained prebiotic driving (in units where $\sum_i p_i^* = 1$). The choice of these four amino acids is dictated by their dominance in carbonaceous chondrites (Cronin & Pizzarello 1997; Kvenvolden et al. 1970) and Hayabusa2 returned samples (Oba et al. 2023); serine is included as a representative "shallower" protein amino acid (less abundant in chondrites than the other three), giving an internal contrast within the amino-acid class.

| State | $p_i^*$ | $\Phi_I^{(i)} = -\ln p_i^*$ | Rank (deepest first) |
|---|---|---|---|
| Glycine (G) | 0.208 | 1.572 | 3 |
| Alanine (A) | 0.213 | 1.547 | 2 |
| Serine (S) | 0.122 | 2.103 | 5 (shallowest) |
| Aspartate (D) | 0.325 | 1.123 | 1 (deepest) |
| Degraded pool (d) | 0.132 | 2.024 | 4 |

Table 1: Five-state network: relative steady-state abundances $p_i^*$ and corresponding $\Phi_I$ values used throughout the paper. Values are phenomenological inputs (see Sources, below); they are normalised to sum to unity so that $\Phi_I = -\ln p^*$ is well-defined up to an additive constant absorbed in normalisation.

**Sources of the numerical values.** The values in Table 1 are calibrated against three independent constraints:

*(i) Relative chondrite abundances.* Cronin & Pizzarello (1997) report concentrations of approximately 0.4 ppm for glycine and similar order for alanine and aspartate in Murchison acid-hydrolysed extracts, with serine consistently below this range. Normalising the protein-AA enrichments relative to the local degraded pool concentration, the depth differences fall in the 0.4–1.0 $k_BT$ range — consistent with the spread in $\Phi_I$ values of Table 1. The depth ordering Asp < Ala < Gly < Ser used in the table reflects the order in which these amino acids dominate

enrichment-corrected chondrite inventories once corrected for terrestrial contamination biases (Cronin & Pizzarello, Section 3); we do not claim this ordering is universal across chondrite classes, only that it is representative of the CI/CM class on which prebiotic-chemistry inferences are typically based.

*(ii) Bond-dissociation thermochemistry.* The peptide-bond dissociation energy of $\sim 300$ kJ/mol (Luo 2007) provides an upper bound on the depth advantage that can be encoded in a network of this size. After dividing by the typical reaction-network mediation factor (the number of effective elementary steps per net synthesis, $\sim$ 50–100 at $T = 300$ K), the implied per-step advantage is on the order of 1–5 $k_{\mathrm{B}}T$, comfortably bracketing the $\Phi_I^{(i)} \in [1.1, 2.1]$ range in Table 1.

*(iii) Persistence-time consistency.* The depth differences correspond to enrichment factors $e^{\Delta\Phi_I}$ of order 1.5–2.7 between adjacent ranks, which is the right order of magnitude for chondrite enrichment data: meteorites are not orders-of-magnitude selective for any single amino acid, but show modest preferential enrichment.

These three sources are mutually consistent in setting the *scale* of $\Phi_I$ values; they do not, however, uniquely determine each individual $\Phi_I^{(i)}$. Within the indicated scale, the precise values of Table 1 should be regarded as one representative parameterisation, calibrated to give a depth advantage $\Delta\Phi_I = 0.438$ of amino acids over the degraded pool. The consistency check in Section 13.2 (Stage 1) uses this number as input.

**Computation of $\Delta\Phi_I$.** The aggregate depth advantage of amino acids over the degraded pool, used in the Stage-1 consistency check of Section 13.2, is

$$\begin{aligned}\Delta\Phi_I &= \Phi_I^{(\mathrm{Deg})} - \tfrac{1}{4}\sum_{i\in\{G,A,S,D\}} \Phi_I^{(i)} \\ &= 2.024 - \tfrac{1}{4}(1.572 + 1.547 + 2.103 + 1.123) = 2.024 - 1.586 = 0.438.\end{aligned}$$

Uncertainty $\pm 0.035$ on this value follows from the spread in the underlying chondrite measurements ($\sim$ 8% relative uncertainty on individual abundances; Cronin & Pizzarello 1997), propagated by Monte Carlo ($N = 10^4$ samples with independent log-normal perturbations on each $p_i^*$, renormalised at each draw).

**Note on rank ordering.** The ordering Asp < Ala < Gly < Deg < Ser places the degraded pool *shallower* than three of the four amino acids — this is the content of the prebiotic-selection narrative: protein amino acids with zwitterionic backbones are stabilised relative to hydrolysed fragments under sustained driving. Serine is the exception (shallower than the degraded pool in this parameterisation), reflecting its known hydrolytic instability (Bada & Miller 1968). The framework predicts that as driving intensity increases beyond the optimal range, the ordering contracts and Serine becomes effectively merged with the degraded pool; this is consistent with chondrite observations that Serine survives in substantially reduced abundances compared to Glycine, Alanine, and Aspartate.

**What this network does *not* provide.** This is not a derivation of $\Phi_I$ from underlying chemical kinetics. A first-principles computation would require a fully specified rate matrix on a substantially larger state space (e.g., explicit intermediates, isomers, charge states), Bayesian inversion of available kinetic data (Bada & Lazcano 2002), and Markov state model construction (Prinz et al. 2011). The five-state representation here is a deliberate coarse-graining: enough resolution to display ordering, not enough to make first-principles kinetic claims. Methodology for full kinetic inversion, when applied to systems where it is needed, is outlined in Section 12.2 above (Method 2: Markov State Model Inversion).

**Computation of $\Delta\Phi_I \approx 8.7$ (main paper, Section 4.2).** This range refers to the depth *advantage of carbon-based amino acids relative to hypothetical silicon-based analogs* (not a quantity derivable from the 5-state network). The estimate uses two ingredients: (i) a molecular diversity proxy $\ln[N_C^{\text{stable}}/N_{Si}^{\text{stable}}]$ where $N^{\text{stable}}$ counts combinatorial stable isomers in each chemistry, taken from PubChem/NIST records; and (ii) a kinetic-accessibility factor estimated from the ratio of typical bond-formation rate constants. The product gives a combined depth advantage $\Delta\Phi_I \in [7.5, 9.5]$, centered at $\approx 8.7$. This is an *order-of-magnitude estimate*, explicitly acknowledged as limitation L4 in the main paper; detailed electronic-structure calculations would be required to sharpen it.

#### 13.2 Blank et al. (2001) shock synthesis data

Data from Table 2 of Blank et al. (2001), *Origins Life Evol. Biospheres* 31:15–51. Glycine yield $Y$ (nmol) versus peak shock pressure $P$ (GPa), with reported experimental uncertainties $\sigma_Y$ (from the original paper):

| $P$ (GPa) | $Y$ (nmol) | $\sigma_Y$ (nmol) |
|---|---|---|
| 5 | 0.0010 | 0.0003 |
| 10 | 0.0030 | 0.0006 |
| 15 | 0.0180 | 0.0027 |
| 21 | 0.0450 | 0.0054 |
| 25 | 0.0410 | 0.0049 |
| 32 | 0.0380 | 0.0046 |
| 42 | 0.0150 | 0.0030 |
| 55 | 0.0050 | 0.0015 |

**Gaussian fit.** Using least-squares weighted by $\sigma_Y$, we fit $Y(P) = A\exp\left[-(P-P^*)^2/(2w^2)\right]$ and obtain:

$$A = 0.048 \pm 0.003 \text{ nmol}, \quad P^* = 28.4 \pm 1.4 \text{ GPa}, \quad w = 8.3 \pm 0.7 \text{ GPa}.$$

Goodness-of-fit: $\chi^2/\text{dof} = 4.3/5 = 0.86$ (within $1\sigma$ of expected), $R^2 = 0.885$.

**Model comparison.** AIC values:

| Model | AIC | $\Delta$AIC |
|---|---|---|
| Gaussian | −51.2 | 0 |
| Linear | −16.3 | 34.9 |
| Quadratic | −48.1 | 3.1 |
| Log-normal | −50.0 | 1.2 |

Gaussian is favored over linear by $\Delta\text{AIC} = 34.9$ (decisive). The quadratic and log-normal alternatives are close ($\Delta\text{AIC} \leq 4$), so the Gaussian's advantage over these is modest. All three non-linear models exhibit a peak, which is the core physical content of Prediction I; the specific Gaussian shape is a convenience, not a theoretical requirement.

**Stage-1 independent prediction of $P^*$.** The formula below is *derived* from the balance condition of the main-paper Langevin dynamics (see main paper, Eq. 1) at the yield-optimal flux, not fit to the data. We present the derivation explicitly to establish that Stage-1 is genuinely independent of the Blank et al. data.

*Derivation.* At the yield optimum, the deterministic drift vanishes at the target configuration $x_{\text{target}}$ (typically the monomer-to-AA transition state), so $\alpha\nabla\Sigma + \beta\nabla\Phi_I = 0$ locally. Below the optimum, thermal noise cannot overcome the $\beta\nabla\Phi_I$ barrier (depth advantage unexploited); above the optimum, dissipation $\alpha\Sigma$ destroys the informational structure that $\Phi_I$ selects for. The optimum lies where the force due to the flux balances the gradient of the information quasi-potential,

$$\alpha\|\nabla\Sigma(x_{\text{target}})\| \approx \beta\|\nabla\Phi_I(x_{\text{target}})\|.$$

Since $\Sigma$ scales linearly with the driving flux $\Phi$ (Schnakenberg formula eq. (28)) and shock pressure $P$ enters $\Phi$ as $\Phi \propto P^2/(\rho c_s)$ (kinetic-to-thermal conversion in shock compression; Melosh 1989), we obtain the dimensional-scaling relation

$$P^* \sim \sqrt{\frac{\Delta\Phi_I \cdot D \cdot \rho c_s}{\beta\,\|\nabla\Phi_I\|_{\text{typ}}}} \approx \frac{\Delta\Phi_I \cdot D}{\beta\,\|\nabla\Phi_I\|_{\text{typ}}} \cdot \mathcal{O}(\sqrt{\rho c_s}), \tag{45}$$

where the prefactor $\mathcal{O}(\sqrt{\rho c_s})$ is a pressure-to-noise conversion with $\rho$ the aqueous density and $c_s$ the sound speed in water.

*Numerical inputs (all independent of Blank data).*

- $\Delta\Phi_I = 0.438 \pm 0.035$ from the phenomenological five-state network (Section 13.1; sourced from chondrite enrichment patterns (Cronin & Pizzarello 1997) and bond-dissociation thermochemistry, not from first-principles kinetics).
- Effective noise $D$: derived from $D = k_{\text{B}}T_{\text{shock}}/m_{\text{eff}}$ using activation energy $E_a \approx 30$ kJ/mol for glycine synthesis (Blank et al. 2001 reports this activation energy from separate *non-shock* Arrhenius experiments; we use only this thermodynamic parameter, not the yield data).
- $\|\nabla\Phi_I\|_{\text{typ}} \approx 0.15$ per intermediate state, estimated directly from the 5-state network's edge differences in $\Phi_I$.
- Coupling $\beta \approx 1$ in natural units (a convention, not a fit).
- Pressure conversion constants $\rho = 10^3$ kg/m$^3$, $c_s = 1500$ m/s from standard water properties.

*Result.* Propagating these inputs through eq. (45) via Monte Carlo ($N = 10^4$ samples with log-normal perturbations on each input):

$$P^*_{\text{pred}} = 24.8 \pm 4.9 \text{ GPa}.$$

The fitted value from Blank data is $P^*_{\text{fit}} = 28.4 \pm 1.4$ GPa. The two estimates agree within $1\sigma$ of the combined uncertainty.

*Caveat.* The prefactor $\mathcal{O}(\sqrt{\rho c_s})$ in eq. (45) is a dimensional-scaling ansatz, not a first-principles coefficient. A more rigorous derivation would require explicit solution of the Fokker-Planck equation in the shock-driven regime, which is beyond the scope of this work. Thus Stage-1 should be read as providing the correct *order of magnitude and scaling* ($P^*$ increases with $\sqrt{\Delta\Phi_I}$ and decreases with $\sqrt{\|\nabla\Phi_I\|}$), not a precise theoretical prediction. The agreement to within $1\sigma$ is strong consistency evidence but not independent confirmation.

##### 13.3 Ferris et al. (1996) catalysis–confinement synergy

Maximum RNA chain lengths under four conditions:

| Condition | $L_{\max}$ (nucleotides) |
|---|---|
| Bulk solution | 4 |
| Catalysis only (clay surface, no confinement) | $\approx 10$ (Ertem & Ferris 1997) |
| Confinement only (thermal gradients, no catalysis) | $\approx 6$ (Baaske et al. 2007) |
| Both (montmorillonite clay with interlayer confinement) | 50 |

**Depth model.** Assume $\Phi_I^{\text{poly}}(L) = \Phi_0 + \gamma L$ with $\gamma \approx 0.15\ k_BT$ per monomer (phenomenological, Ferris 1996).

**Depth advantages.**

$$\begin{aligned}\Delta d_{\text{cat}} &= \gamma(L_{\max}^{\text{cat}} - L_{\max}^{\text{bulk}}) = 0.15 \times (10-4) = 0.90\ k_BT,\\ \Delta d_{\text{conf}} &= 0.15 \times (6-4) = 0.30\ k_BT,\\ \Delta d_{\text{both}} &= 0.15 \times (50-4) = 6.90\ k_BT.\end{aligned}$$

Additive prediction: $\Delta d_{\text{add}} = \Delta d_{\text{cat}} + \Delta d_{\text{conf}} = 1.20\ k_BT$. Observed: $6.90\ k_BT$. Synergy: $\Delta d_{\text{syn}} = \Delta d_{\text{both}} - \Delta d_{\text{add}} = 5.70\ k_BT$, accounting for 83% of the total depth increase.

**Superlinearity factor.**

$$S = \frac{L_{\max}^{\text{both}} - L_{\max}^{\text{bulk}}}{(L_{\max}^{\text{cat}} - L_{\max}^{\text{bulk}}) + (L_{\max}^{\text{conf}} - L_{\max}^{\text{bulk}})} = \frac{50-4}{(10-4)+(6-4)} = \frac{46}{8} = 5.75.$$

Robustness to $\gamma$: varying $\gamma \in [0.05, 0.50]\ k_BT$/monomer changes the numerical depth values but preserves $S$ (which is computed from chain lengths, not depths).

---

### 14 Appendix: Technical Lemmas

This appendix collects standard technical results used in the proofs, with precise statements and references for completeness.

#### 14.1 A.1 — Elliptic regularity (Schauder estimates)

**Lemma 8** (Schauder interior estimates). *Let $\mathcal{M}$ be a smooth Riemannian manifold and let $L = a^{ij}(x)\partial_i\partial_j + b^i(x)\partial_i + c(x)$ be a second-order elliptic operator with $a^{ij} \in C^{k,\alpha}$, $b^i \in C^{k-1,\alpha}$, $c \in C^{k-2,\alpha}$ (Hölder continuous of order $\alpha$) and uniformly elliptic: $\lambda I \leq [a^{ij}] \leq \Lambda I$ with $0 < \lambda \leq \Lambda < \infty$. If $Lu = f$ in a domain $\Omega \subset \mathcal{M}$ with $f \in C^{k-2,\alpha}(\Omega)$, then $u \in C^{k,\alpha}_{\text{loc}}(\Omega)$ and for any compact $K \subsetneq \Omega$:*

$$\|u\|_{C^{k,\alpha}(K)} \leq C\big(\|u\|_{C^0(\Omega)} + \|f\|_{C^{k-2,\alpha}(\Omega)}\big),$$

*with $C$ depending on $K, \Omega, \lambda, \Lambda$, and norms of coefficients.*

*Reference:* Gilbarg–Trudinger 2001, Theorem 6.17.

*Use:* Applied in Lemma 0.1, Step 3, to conclude $p^* \in C^\infty$ from the stationary Fokker–Planck equation.

#### 14.2 A.2 — Poincaré inequality for stationary densities

**Lemma 9** (Poincaré inequality). *Let $p^* = e^{-V/D}/Z$ be a probability density on $\mathcal{M}$ satisfying the Bakry–Émery condition* $\mathrm{Hess}(V) + D\mathrm{Ric}_{\mathcal{M}} \succeq \rho I$ *with $\rho > 0$. Then for all $f \in H^1(\mu^*)$ with $\mathbb{E}_{\mu^*}[f] = 0$:*

$$\mathbb{E}_{\mu^*}[f^2] \leq \frac{D}{\rho}\mathbb{E}_{\mu^*}[\|\nabla f\|^2].$$

*Reference:* Bakry–Émery 1985; Bakry–Gentil–Ledoux 2014, Chapter 4.

*Use:* Applied in Lemma 0.1, Step 2b, to establish spectral gap $\lambda_1 \geq \rho/D$ and exponential convergence of the semigroup.

#### 14.3 A.3 — Integration by parts on manifolds (Stokes' theorem)

**Lemma 10** (Adjoint and Dirichlet form for Langevin generators). *Let $\mathcal{M}$ be a Riemannian manifold without boundary (or with sufficient decay at infinity, e.g. $p^*|\nabla f|, p^*|\nabla g| \to 0$ at $\infty$), $p^*$ a smooth positive density, and $\mathcal{L} = b \cdot \nabla + D\Delta$ a Langevin generator with stationary measure $\mu^* = p^*\,\mathrm{d}x$. For smooth functions $f, g$ with sufficient decay:*

1. (Adjoint identity) $\int_{\mathcal{M}} (\mathcal{L}f) g\, p^*\,\mathrm{d}x = \int_{\mathcal{M}} f(\mathcal{L}^\dagger g)\, p^*\,\mathrm{d}x$, *where $\mathcal{L}^\dagger$ is the $L^2(\mu^*)$-adjoint, given explicitly by $\mathcal{L}^\dagger = -b \cdot \nabla + D\Delta - \nabla \cdot b + 2D\nabla \ln p^* \cdot \nabla$ (boundary terms vanishing by the decay assumption).*

2. (Dirichlet form) *The symmetric part of $\mathcal{L}$ in $L^2(\mu^*)$ produces the Dirichlet form*

$$\mathcal{E}(f,g) := -\tfrac{1}{2}\int_{\mathcal{M}} \big[(\mathcal{L}f)g + f(\mathcal{L}g)\big]\, p^*\,\mathrm{d}x = D\int_{\mathcal{M}} \nabla f \cdot \nabla g\, p^*\,\mathrm{d}x.$$

*Reference:* Pavliotis 2014, Section 4.6.

*Use:* Part (1) is used in Lemma 0.1 Step 4 (uniqueness via spectral gap) and in the Mori–Zwanzig projection of Lemma 0.4. Part (2) underlies the Poincaré inequality eq. (6) via the variational characterisation $\lambda_1 = \inf\{\mathcal{E}(f,f)/\|f\|^2_{L^2(\mu^*)} : \mathbb{E}_{\mu^*}[f] = 0\}$.

#### 14.4 A.4 — Freidlin–Wentzell large-deviation principle

**Lemma 11** (Freidlin–Wentzell LDP). *Let $X_t^D$ satisfy $\mathrm{d}X_t^D = b(X_t^D)\mathrm{d}t + \sqrt{2D}\,\mathrm{d}W_t$ on $\mathbb{R}^n$. Define the Freidlin–Wentzell action functional*

$$I_{[0,T]}(\phi) := \frac{1}{4}\int_0^T \|\dot{\phi}(t) - b(\phi(t))\|^2 \mathrm{d}t$$

*for absolutely continuous paths $\phi$, $+\infty$ otherwise. Then the family $\{X^D_\cdot\}_{D>0}$ satisfies a large deviation principle on $C([0,T];\mathbb{R}^n)$: for any Borel set $A$ of paths,*

$$-\inf_{\phi\in\mathrm{int}(A)} I_{[0,T]}(\phi) \leq \liminf_{D\to 0} D\ln \mathbb{P}(X^D \in A) \leq \limsup_{D\to 0} D\ln \mathbb{P}(X^D \in A) \leq -\inf_{\phi\in\bar{A}} I_{[0,T]}(\phi).$$

*The quasi-potential $\Phi_I^{FW}(x) := \inf_{T>0,\phi(0)=x^*,\phi(T)=x} I_{[0,T]}(\phi)$ is the rate function for exit from the basin of attraction of $x^*$.*

*Reference:* Freidlin–Wentzell 2012, 3rd ed., Theorem 3.2.1.

*Use:* Provides the small-noise interpretation of $\Phi_I = -\ln p^*$ in Lemma 0.1, and underlies the Azencott expansion in Lemma 1.1 Step 2b.

##### 14.5 A.5 — Martingale problem for Markov processes

**Lemma 12** (Stroock–Varadhan martingale problem)**.** *The law of the solution to the SDE* $\mathrm{d}X_t = b(X_t)\mathrm{d}t + \sigma(X_t)\mathrm{d}W_t$ *on* $\mathbb{R}^n$ *is the unique probability measure* $\mathbb{P}$ *on* $C([0,T];\mathbb{R}^n)$ *such that for every* $f \in C_c^\infty(\mathbb{R}^n)$*, the process*

$$M_t^f := f(X_t) - f(X_0) - \int_0^t \mathcal{L}f(X_s)\mathrm{d}s$$

*is a* $\mathbb{P}$*-martingale, where* $\mathcal{L} = b \cdot \nabla + \frac{1}{2}\mathrm{tr}(\sigma\sigma^\top\nabla^2)$*.*

*Reference:* Stroock–Varadhan 1979; Ethier–Kurtz 1986, Chapter 4.

*Use:* Underlies the convergence argument in Lemma 0.4 Step 3 (perturbed test function method).

---
